# An Automated Radio-Telemetry System (ARTS) for Monitoring Small Mammals

**DOI:** 10.1101/2021.03.06.434221

**Authors:** Gerard Wallace, Marija Elden (née Gorinshteyn), Rachel Boucher (née Sheely), Steven Phelps

**Affiliations:** Department of Integrative Biology, University of Texas at Austin, 2415 Speedway, Austin, TX, 78712

**Keywords:** ARTS, Telemetry, Rodent, Prairie Vole, Ultradian

## Abstract

Point 1: The study of animals in nature is essential for developing an ecologically valid understanding of behavior. Small mammals, however, are often fossorial and exceedingly difficult to monitor in the wild. This limits both the taxonomic scope of field observation, and excludes species that are powerful models for the study of behavioral mechanisms.

Point 2: Here, we implement an automated radio telemetry system (ARTS) designed to track small fossorial mammals. Our ARTS uses an isotropic antenna array coupled with broadband receivers. We characterized transmission at our study site and tested the ARTS’ ability to track 48 prairie voles.

Point 3: We compared position estimates from nonlinear least squares, nonparameteric, and Bayesian trilateration methods and found Bayesian trilateration to have the smallest error. To examine the ability of the system to track biologically significant behavior we used ARTS data to investigate circadian rhythms of freely behaving prairie voles. We used Lomb-Scargle analysis to estimate periodic patterns from irregularly sampled time series of speed. Prairie voles demonstrated ultradian movement at periods of approximately 45 and 90 min, observations on a time scale not possible using data from traditional methods.

Point 4: This ARTS offers a new tool to observe rodent field behavior at time scales and in environments which have not been previously possible, such as investigating social and spatial behaviors on the scale of minutes, hours, and days in natural environments.

## Introduction

Ethologists and behavioral ecologists have long emphasized the importance of examining behavior in settings relevant to the ecology and evolution of a species (Tinbergen 1951). Increasingly, researchers in psychology and neuroscience have come to share this perspective; their approaches include examining the statistical properties of real-world sensory stimuli (Burge and Geisler, 2015), examining the mechanisms of memory in nature (Sonnenberg et al., 2019), and exploring the behavioral patterns that emerge among complex social groups (Hein et al., 2018). In parallel, researchers have begun analyzing exhaustive sets of behavioral data with the hope of more thoroughly characterizing the behavioral repertoire of a species (Anderson and Perona, 2014; Schaefer and Claridge-Chang, 2012). To support such advances, we describe a new implementation of an Automated Radio-Telemetry System (ARTS) for use with small animals in the field.

One challenge of examining behavior in natural settings is that many animal species are relatively small and secretive, making field observation difficult. Among fossorial muroid rodents, for example, knowledge of natural behavior is severely limited (Taborsky et al., 2015). The socially monogamous prairie vole, for example, has become a popular model for understanding the neurobiology of bonding and attachment. However, despite decades of excellent work in both the lab and field, we still know relatively little about the evolutionary and ecological pressures that promote monogamy. In prairie voles and other small rodents, field methods limit the ability of researchers to integrate ecological and evolutionary perspectives with the study of genetic, physiological and neurobiological mechanisms – an integration essential to understanding the animal in its world (Tinbergen, 1951).

Ecologists employ a variety of sensors for monitoring animals in the wild. These include global-positioning systems (GPS) tags, proximity tags, passive-integrated transponders (PIT) tags and automated radio-telemetry systems (ARTS). Each of these has strengths and limitations. Among these, GPS is the most common method for tracking animals in natural environments; however, many species are too small for GPS tags (Kays et al., 2015). Proximity tags record when two collars are near one another, generating high resolution spatially explicit records of social interaction. Unfortunately, proximity tags that fit small animals can only record several hours’ worth of data (Levin et al., 2015). Passive-integrated transponder tags have no batteries, are small and light, and have become a powerful tool for tracking small animals. For example, PIT tags have been deployed to automate access to bait stations, allowing spatial learning to be measured directly in the wild (Croston et al., 2016). Unfortunately, PIT tags require animals to come within 10 to 30 cm of the tag reader to be detected (Bridge and Bonter; Gibbons and Andrews, 2004); and although PIT tags are individually cheap, surveying large areas or high spatial resolutions with tag readers may be prohibitively expensive.

ARTS consist of an array of receiving antenna used to localize radio transmitters via triangulation. ARTS have been used to track activity in a variety of small bird, reptile, and mammal species in the field (Celis-Murillo et al., 2017; DeGregorio et al., 2016; DeGregorio et al., 2018; Hoffmann et al., 2018; Kays et al., 2011; Sperry et al., 2013; Steiger et al., 2013; Ward et al., 2013). For example, in field sparrows [*Spizella pusilla*] and yellow-breasted chats [*Icterina virens*] location predictions from ARTS have been used to study social behavior and reproductive tactics by observing extra-territorial forays, where birds are presumed to solicit extra-pair copulations (Celis-Murillo et al., 2017; Ward et al., 2014). Like alternative methods, ARTS have limitations. Most notable is difficulty estimating position based on received signal strengths – although the relationship between distance and signal strength has a clear physical description, in natural settings, reflections and other propagation irregularities add noise to the received signal strength. Prior ARTS have overcome this problem by focusing on tracking across very large scales, or by using rotating, directional antennae to localize a bearing rather than a position. Lastly, prior ARTS use sensitive narrow-channel receivers that detect the signal from an individual transmitter. Such sensitivity is not needed at the scale of small mammals, and comes at the cost of needing to sequentially cycle through each subject’s transmission frequency, greatly reducing the temporal precision of estimates.

Here we present a novel implementation of an ARTS system designed specifically for the automated, high-throughput monitoring of small animals. The ARTS system we deployed uses traditional VHF radiotransmitters coupled with ultra-wideband receivers receiving input from an array of omni-directional (isotropic) antennae. In this paper, we document signal propagation at our study site, then test the accuracy of various methods of estimating position based on the signal strength received by different antennae. These trilateration methods include nonlinear least squares (NLS), nonparametric (NP), and Bayesian models. We assessed these methods using a testing dataset consisting of detailed mapping of a single transmitter in multiple positions throughout the tracking areas, as well data obtained by traditional manual telemetry of 48 different prairie voles living within the outdoor enclosures. Finally, we demonstrate the ARTS can detect meaningful biological variation by identifying activity cycles on a time scale too short to have been detected using manual methods.

## Methods and Methods

### Study Site

The study site (Fig 1A) was located in Jackson Co., IL, within the natural range of prairie voles (Tamarin, 1985). This site has previously been used to investigate the behavioral and endocrine effects of population density in prairie voles and is described in detail in Blondel et al. (2016). The site covered an area of 0.4 hectares, consisting of four adjacent quadrants measuring 40 m by 30 m (0.1 hectare apiece). Removable gates between quadrants allow the area available to study animals to be manipulated. Predators and native rodents were excluded by two 60 cm tall aluminum flashing borders, a 3 m wide mowed “moat”, aviary netting and an electric fence. Sixteen baited traps (Sherman, Tallahassee, FL, USA) were present in the moat, to prevent exit of study animals or entry of wild rodents into the field site. To further prevent subject animals from escaping, the enclosure grasses and other plants were excluded from a perimeter within 0.5 m of all flashing boarders, creating open space which rodents are less likely to cross. Survey flags were placed in a 3 x 3 m resolution grid within the enclosure to assist location measurement during mapping and tracking. Coordinates within the enclosure were recorded on a Cartesian grid with an origin at the northwest corner on the enclosure. Distance south of the origin was the positive x, and distance east of the origin as was the positive y. The coordinate system was rotated *π*/2 radians clockwise to orient the x and y coordinates to Northing and Easting (the Cartesian coordinates of the Mercator projection).

**Fig 1.**
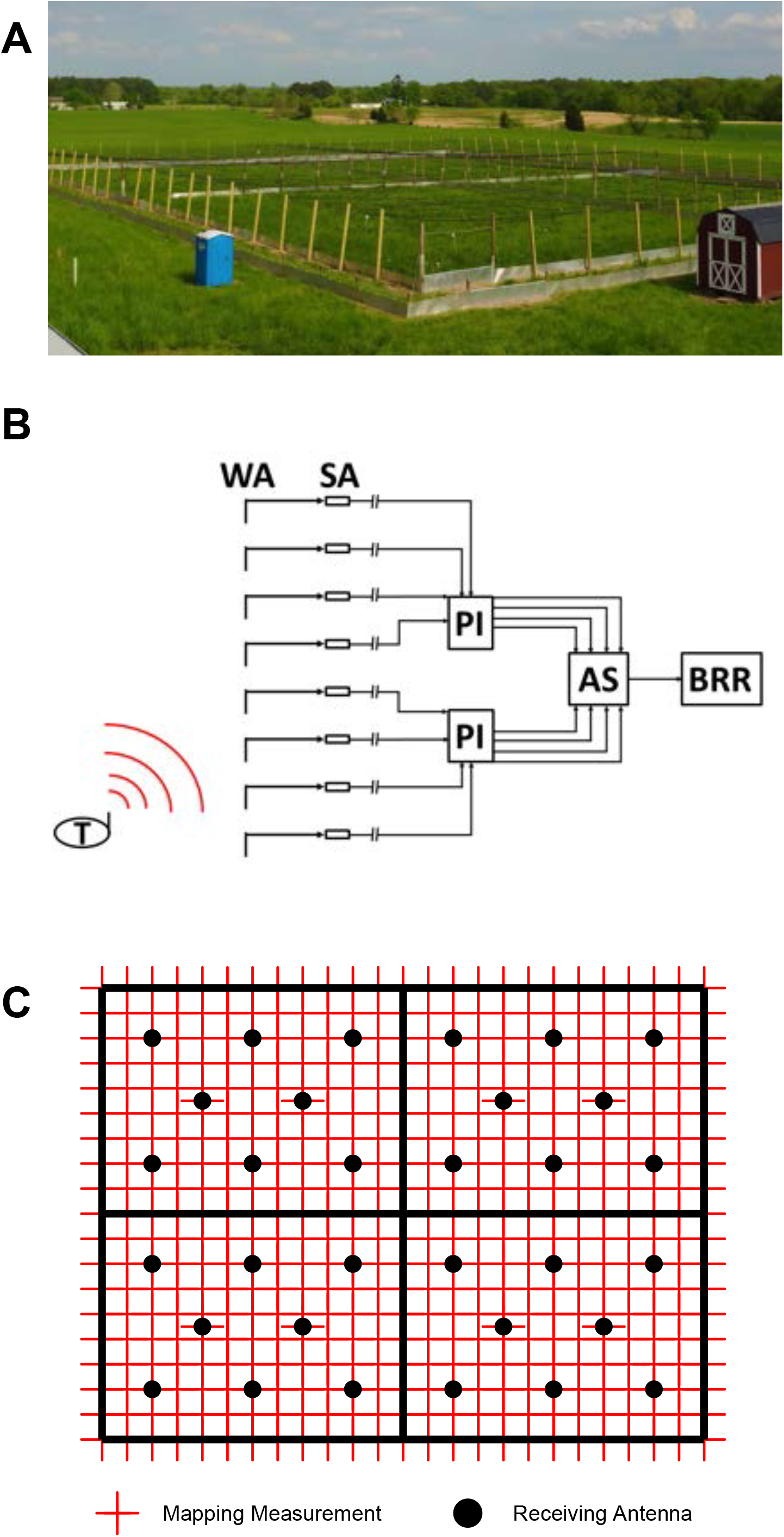
Field Site and Automated Telemetry System. (A) Jackson County, IL Field Site. (B) Automated telemetry system. Signal from radio transmitters (T) are recieved at Whip antenna (WA), signal is boosted at signal amplifier (SA), power is delivered to SA through in-lin power inserters (PI), WA are selected va an antenna switch (AS), broadband radio receiver (BRR) receives all transmitter frequencies within a 1 MHz band from selected WA. (C) Enclosure antenna layout (black) and 3 x 3 m radiomapping measurement grid (red).

**Fig 2.**
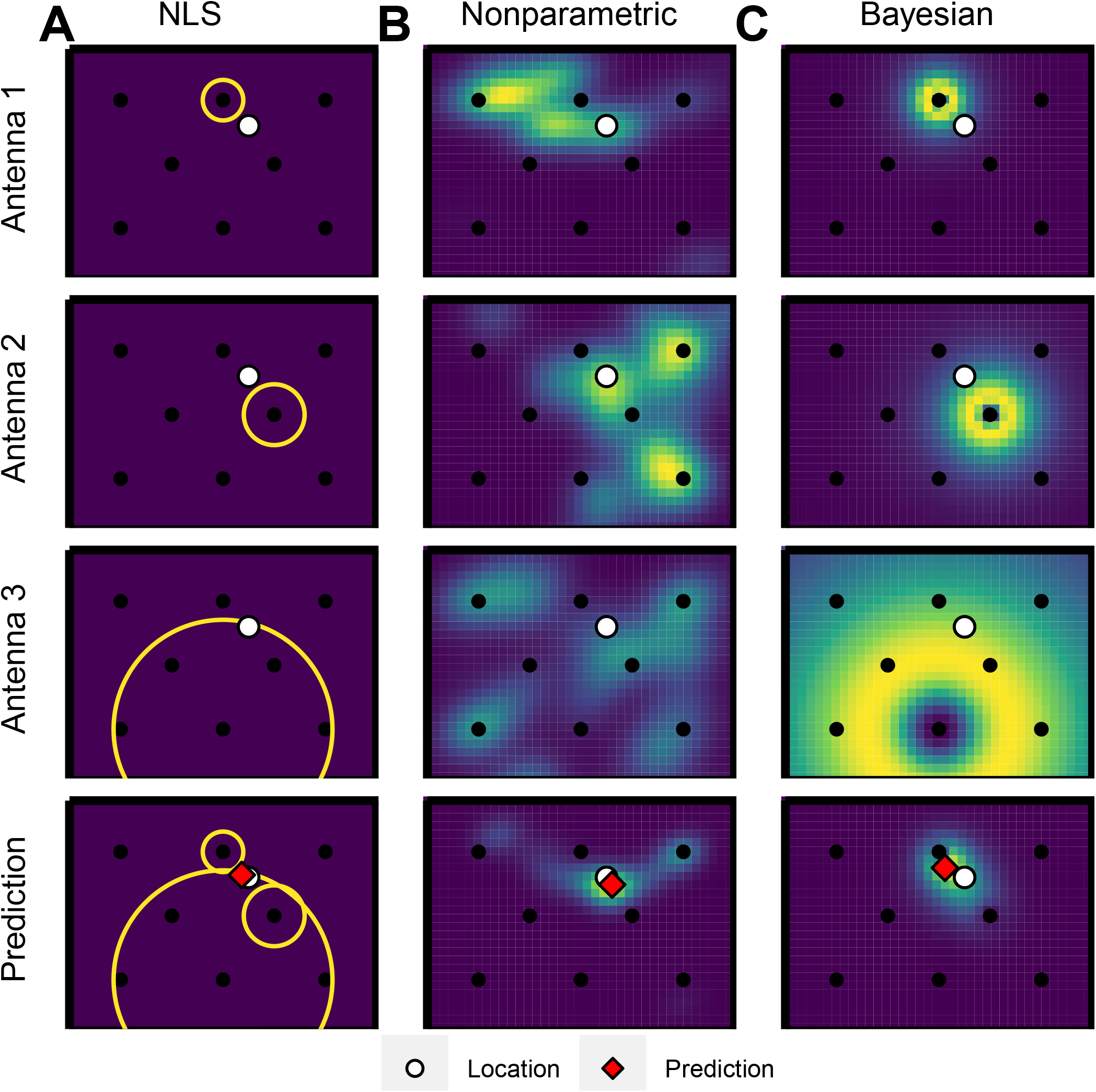
Localization Algorithm Illustration. (A) Nonlinear least squares trilateration. Location estimate minimizes the distance between projected distances from each antenna. (B) Nonparametric trilateration. Location estimate maximizes the product of the KDE of position for each antenna. (C) Bayesian trilateration. Location estimate maximizes the a posteriori joint probability of position.

### Animals

All animal procedures were approved by the University of Texas at Austin IACUC. Prairie voles [*Microtus ochrogaster* (Baird 1857)] derived from Jackson Co., IL founders were bred at the University of Texas at Austin. Voles were ear tagged at weaning (p21) for identification. Enclosures included locally common grasses such as rye, brome and fescue, and was seeded with patches of clover. Naturally available water was supplemented with two 11.6 L poultry fountains placed within each quadrant. Study animals and offspring were trapped 6 weeks following the start of tracking using 7.62 x 8.89 x 22.86 cm folding traps (Sherman, Tallahassee, FL, USA). Traps were baited with a mixture of seeds, oats and peanut butter. Forty-four traps were arrayed in a grid within each quadrant and armed between 18:00 and 8:00.

### Automated Radio-Telemetry System

The automated telemetry system consisted of an array of 32 isotropic receiving antennae connected to four broadband receivers (model Orion, Sigma 8 Inc) housed in the field lab (Fig 1C). To offset signal decay between the antennae and the receivers, signal amplifiers powered by inline power inserters, located within the lab, were placed on the receiving antenna (Fig 1B). The receivers monitored a 1 MHz band from a single antenna, and each receiver switched sequentially among eight antennae using antenna switches (model Hydra, Sigma 8 Inc). Data from separate receivers was concatenated offline. Each radio receiver truncated all signals above -57 dBm and censored all signals below -113 dBm. Isotropic transmitters (Wildlife Materials) were in the 151 MHz band. The transmitters delivered a -30 dBm pulse for 20 ms at a rate of 0.5 Hz. Transmissions were filtered by the Orion firmware using incoming signal by pulse width (15 to 25 ms), interpulse interval (1500 ms to 2500 ms), intra-burst power (±6 dBm) and inter-pulse power (±6 dBm).

### Radiomapping and Manual Radio Tracking

To characterize the relationship between received signal strength (RSS) and distance we “radiomapped” the field site. Received signal strength was measured across the enclosure at a resolution of 3 m x 3 m (475 locations, Fig 1C) using a test transmitter (model SOM 2028 HWSC, Wildlife Materials Inc.) at 151.115 MHz. To account for variation in RSS due to potential anisotropy in the transmitters, the test transmitter was rotated through four orientations at each measurement location. The test transmitter was measured for 60 sec in each orientation at each location.

To validate ARTS predictions voles were collared with transmitters (model SOM 2028 HWSC, Wildlife Materials) in the 151 MHz band, weighing no more than 2.5 g. All collar frequencies were separated by at least 0.01 MHz. Voles were observed 24 hrs before release to ensure collars were secure. Prairie vole territories are primarily established through female-female competition for resources (Ostfeld, 1985; Wolff, 1993) and males do not maintain a territory when unpaired (Getz and McGuire, 1993; McGuire et al., 2013). Thus, females were released first and males were released 24 hrs later to facilitate territory formation. Voles were released at low and high densities of 80 and 240 animals/hectare respectively. Each density consisted of twelve males and twelve females. The low-density treatment consisted of the Northwest, Northeast, and Southeast quadrants. The high-density treatment consisted of the Southwest quadrant. The voles were tracked manually for six weeks with a narrow-band receiver (model TRX-1000, Wildlife Materials Inc) and three-element folding yagi antenna via homing (Kenward, 2000). Weather permitting, voles were located up to twice daily. Time of localization was recorded to the nearest minute. Tracking order was randomized and split into three periods AM (6:00 -10:00), Noon (10:00 -14:00), and PM (14:00-18:00).

Home ranges were estimated by bivariate-normal kernel density estimates (KDE) using manual telemetry fixes in the R package “adehabitateHR” (Calenge, 2006). A vole’s nest site was taken to be the location corresponding to its kernel maximum. Kernel density estimates was recorded in the supplemental “2014_KDE_Fitting.R” script.

### Data Processing

As mentioned in the ARTS description, data matched inter-pulse interval and inter-pulse power of the transmitters; data not matching these criteria were excluded from analysis. Data from radiomapping and tracking were loaded, concatenated, and filtered in R (R Core Team, 2018). Following cleaning, data were binned to the nearest minute. The data cleaning process was recorded in the supplemental “Filtering.R” script with functions available in the “ARTSFunctions.R” file. Manual measures of transmitter position and automated measure of RSS were combined offline.

Our design provided two measures of the relationship between signal strength and position: First was the radio-mapping data, which provided relatively exhaustive measures of position and orientation for a single transmitter; and second, the manual telemetry of live animals, which includes a larger number of transmitters that capture the natural variation of conditions in the field, including animal orientation, distortion of signal strength by the animal body, visits to burrows, and the positions of researchers and equipment within the enclosures. To allow comparisons between ARTS estimates and known transmitter positions, localizations from manual telemetry were joined to corresponding data from the ARTS by time and frequency. Specifically, the temporally closest antenna contacts within five minutes of a manual localization were paired. Pairing was performed with the supplemental “ARTS_vs_Manual.R” script.

Localizations were randomly assigned to training or testing datasets. Training data sets were used to estimate the signal decay model and kernel density estimates of position. Testing data sets were used to compare the accuracy of telemetry models by comparing the ARTS position estimate to the known transmitter position in mapping and manual telemetry data. The test training partition was recorded in the supplemental “split_training_testing.R”.

### Position Estimation and Analysis

We compared three mathematical algorithms for estimating position based on received signal strength (RSS) of radio transmitters. In all three algorithms, we focus on the RSS of the three antennae that received the largest signal strength. (Initial analyses revealed that including additional antennae did not improve our results.) Non-linear least squares trilateration, uses mapping data to regress RSS against the log of distance for each antenna. This relationship predicts the distance of a transmitter to an antenna based on RSS, producing a circle of possible positions relative to each antenna. If the relationship between RSS and distance were error-free, there would be one position where the circles from the three antennae intersect; since it is not, we used NLS regression to estimate position. The second method, which we term Bayesian trilateration, uses a similar logic, but includes explicit considerations of error in the estimation of the relationship between RSS and distance. The third method, NP trilateration, does not assume isotropic decay of RSS with distance. Instead, we used the mapping data to estimate the probability that a transmitter was in a given location given its RSS. We used the joint distribution of these probabilities for the three antennae to estimate the most likely position. To compare the precision of these methods, we compared estimates to known positions of isolated transmitters in our mapping data, or to positions of transmitters attached to moving animals who had been tracked by manual telemetry.

To examine whether our methods could estimate novel, biologically meaningful variation in behavior, we estimated the periodicity of movements using a method known as the Lombs-Cargle Periodogram (Lomb, 1976; Scargle, 1982). The mathematical details of these three methods, the software used for their analysis, and the details of periodicity estimation, are provided in the supplement to this manuscript.

## Results

Signal propagation within the study site was investigated by measuring RSS in a 3 x 3 m grid across the study site. After filtering there were 87055 contacts with antenna corresponding to 2836 time points with known transmitter locations. Signals from at least three distinct antennae were observed at 91.5% of time points. The mapping transmitter could be detected by at least three distinct receiving antennae at 98.1% of measurement locations (Fig 3A). The median area of the minimum convex polygons of fixes for within the study site was 3296 ± 459 m^2^. The median of minimum convex polygon overlap was 29 ± 2 (range 22, 31; Fig 3B). As expected, antennae were approximately isotropic, and RSS decayed logarithmically with distance. Thus, a trilateration-based localization strategy was deemed feasible within the study site. We noted some antennae showed much shallower rates of decay or multiple RSS peaks, often at other antenna sites. This was interpreted as either noise from signal amplifiers short circuits or bleed through in the antenna switch (Fig S1-S4).

**Fig 3.**
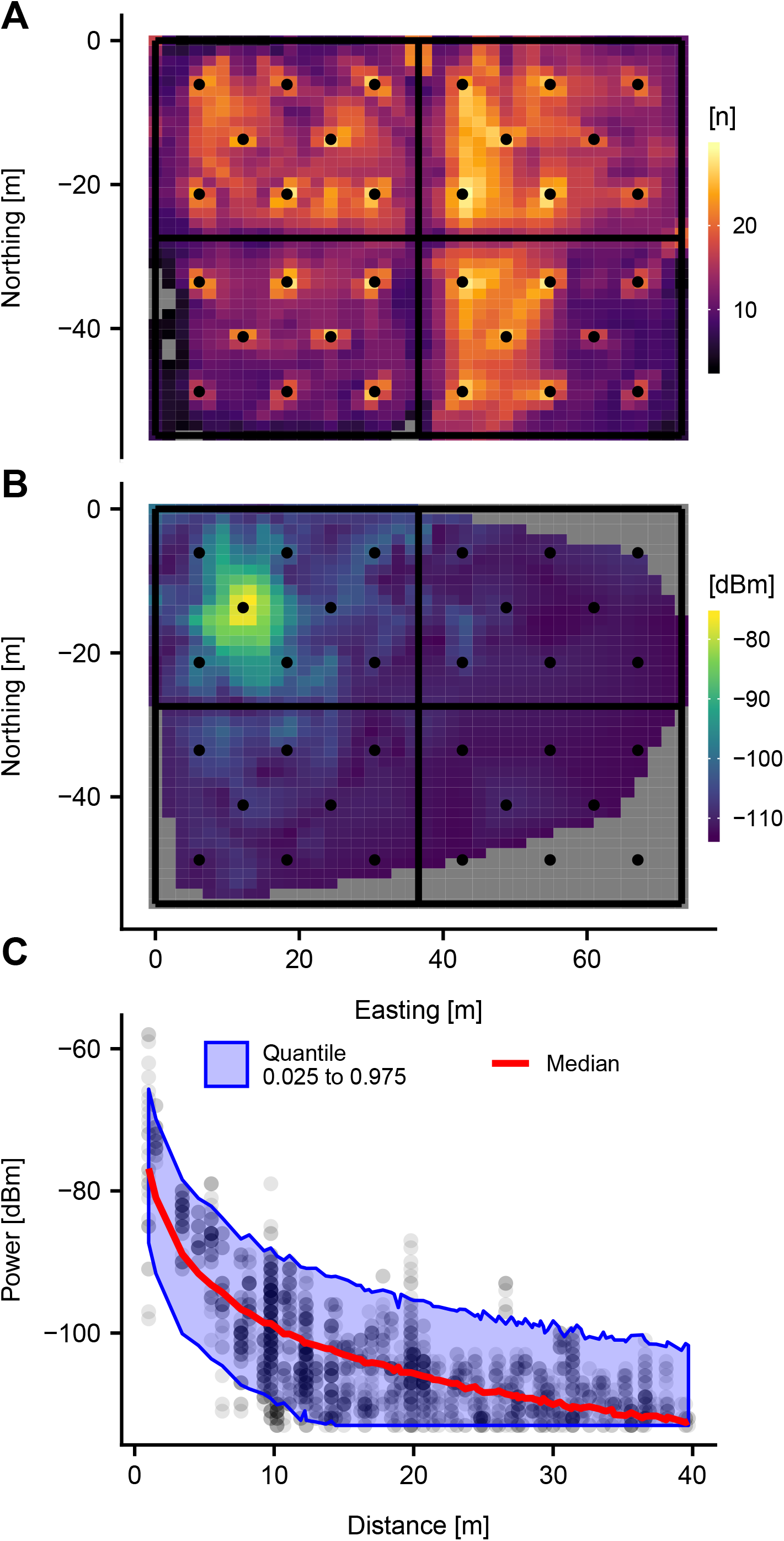
Empirical Characterization of Signal Propagation Within Study Enclosure. (A) Number of antenna receiving signal from a given location within the enclosure, grey denotes fewer than 3 unique antenna. (B) Receptive field for the antenna located at 12.2 Easting, -13.7 Northing. (C) Signal decay for the antenna located at 12.2 Easting, -13.7 Northing predicted by hierarchical Bayesian model. Truncated to 40 m.

Forty-eight prairie voles were radio collared and simultaneously tracked by the ARTS and manual telemetry for six weeks to test accuracy of ARTS based localization. 1292 fixes were recorded by manual telemetry, with a median of 15 ± 41.5 fixes per animal. After data filtering there were 217242 ARTS fixes (time points with RSS from at least three antennae) during the tracking period. The median number of ARTS fixes was 6820 ± 8970. Approximately 38% (497/1292) of manual telemetry fixes had an ARTS fix within 5 min. The median time between ARTS fixes was 2 ± 2 min.

Prairie voles may nest on the surface or in burrows approximately 5 to 25 cm underground (personal observation; Harper and Batzli, 1996b). Being underground has the potential to alter propagation in ways not accounted for in the radiomapping dataset. However, no effect of being near the nest the nest (manual fix within 2 m of manual KDE max) vs away from the nest was observed, accounting for distance between transmitter and antenna and collar frequency (as a random effect).(*β*_*nest*_ = 1.43, df = 1, p > 0.05) (Fig.S5).

Signal propagation may also be altered by researcher presence during tracking and subject animals acting as signal amplifiers (Levin et al., 2015; Naef-Daenzer et al., 2005). To account for this during manual tracking, data from both radiomapping and manual telemetry were used to train the Bayesian signal decay model and nonparametric KDE’s. In both datasets 50% of the localizations were randomly selected for training and 50% was used for testing. The training dataset contained 1521 fixes (1275 mapping, 246 tracking) and the testing dataset contained 1640 fixes (1397 mapping, 243 trapping).

The antenna parameter estimates from the Bayesian signal decay model are reported in table S1. Table 1 summarizes the distribution of errors in the testing dataset for each localization method. The Bayesian trilateration and NLS trilateration were substantially more accurate than nonparametric trilateration (Fig 4B). However, neither Bayesian nor NLS trilateration were large improvements on using the antenna as RFID detectors (assigning the location to the coordinate of the antenna with the strongest signal). For a subset of fixes in the testing data set (37), NLS trilateration did not converge. Both the Bayesian and nonparametric models estimate the probability distribution of a localization permitting evaluation of fix credibility contours. Bayesian trilateration preformed substantially better than nonparametric trilateration. Credibility contours were calculated for each fix in the testing data set from the estimated Bayesian and NP posterior densities (eg. the 50% contour outlines the isopleth defining the most likely 50% of the distribution of position). The NP contours were far too concentrated relative to the testing data, while the Bayesian contours showed only a modest negative bias in contour dispersion (Fig 4C). Further, the areas of the NP contours were systematically smaller than the Bayesian trilateration contours (Fig 4D). This likely reflects overfitting of the NP trilateration to the training data. Prediction errors for the Bayesian trilateration method was strongly related to minimum received signal strength within a fix (Fig 4D).

**Table 1.** Summary of Localization Error Distributions (m)

| Method | MSE (m<sup>2</sup>) | Q<sub>1</sub> | Mode | Median | Mean | Q<sub>3</sub> |
| --- | --- | --- | --- | --- | --- | --- |
| Bayes | 143.6 | 0.6 | 0.9 | 4.3 | 7.2 | 9.3 |
| RFID | 152.0 | 0.0 | 0.2 | 4.8 | 7.4 | 9.8 |
| NLS | 200.9 | 1.7 | 1.0 | 5.2 | 8.7 | 11.0 |
| NP | 245.3 | 5.9 | 5.9 | 10.1 | 12.5 | 16.7 |
| Random | 1364.8 | 20.5 | 35.4 | 32.8 | 33.3 | 45.3 |

**Fig 4.**
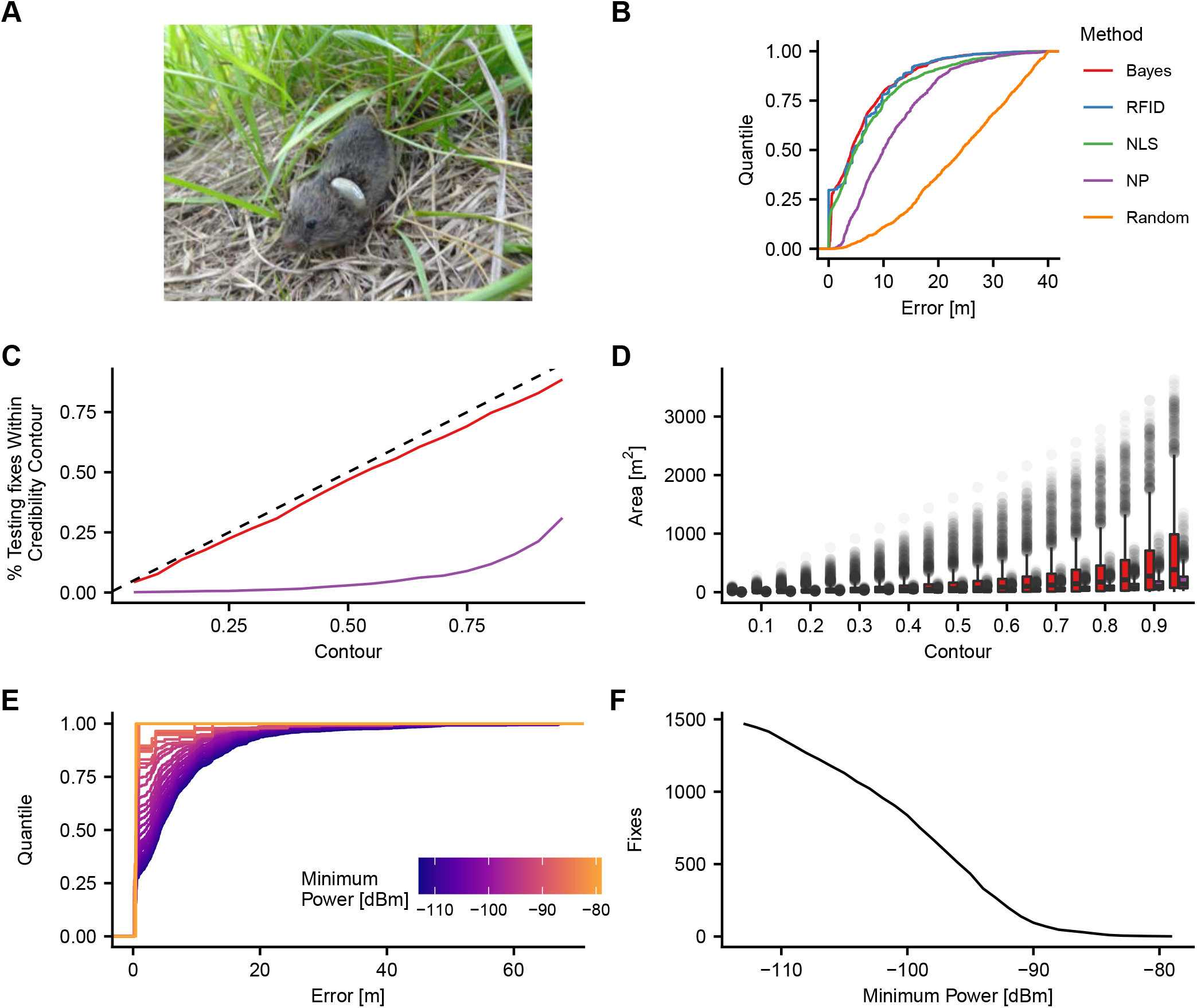
Localization Algorithm Accuracy. A) A prairie vole collared with a radio transmitter within the study enclosure. 48 collared voles where tracked by manual radiotelemetry to establish a training dataset. (B) Empirical cumulative density of error for NLS, Nonparametric, and Bayesian trilateration. (C) Proportion of fixes within the testing data set falling within a given contour level. (D) Contour area as a function of contour level. (E) Empirical cumulative density of error as a function of minimum RSS in the fix. (F) Number of localizations in the testing data as a function of minimum RSS.

The Bayesian model was tested for sensitivity to the amount of training data, and to outliers in the training dataset. The model was trained on fractions of the radiomapping and tracking datasets ranging from 10% to 90% and tested on the complementary fraction. Increasing or decreasing the amount of training data did not substantially influence the accuracy of the model (Fig S6A). To test for sensitivity to outliers the model was fit with the 50/50 training/testing split as described in the methods. The residuals from the training data were ranked. Data from fixes in the 80th through 99th percentiles were used to refit the model. Bayesian trilateration predictions were insensitive to the exclusion of outlying fixes in the training data (Fig S6B). Fix errors were sensitive to which antenna identity (Fig S6C), however the overall distribution of errors was insensitive to the contributions of individual antenna (Fig S6D).

Prediction errors were examined as a function of manual fix location (Fig S7A), revealing a mostly homogeneous distribution of error as a function of space within the enclosure. Differences in prediction error between density treatments or study area quadrants were not apparent.

Data from the second week of manual telemetry was analyzed for circadian patterns. The second week was selected because it represented a substantial duration where the voles were expected to have acclimated to the study area and which was not confounded by attrition due to lost collars or death. Because the Bayesian model had the lowest mean-squared error it was selected to analyze the ARTS data for circadian patterns. Speed was calculated as 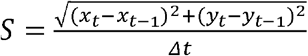. During the second week of tracking sufficient data were available for 24 animals (11 Low Density, 13 High Density). Time series data indicated that animals exhibit period bouts of activity, approximately every 45 min with slightly increased activity daytime activity (Fig 5A). Lomb-Scargle analysis revealed consistent periodic behavior in the low-density treatment (Fig 5BC), but not in the high-density treatment (Figs S8-11). All low-density windows showed peaks at approximately 5.67 MHz (a period of ∼2.94 min). This window activity was also present at the subharmonic at approximately 2.86 MHz (a period of 5.88 min). These peaks were considered artifacts, likely driven by antenna cycling. All low-density windows also showed a complex pattern of peaks ranging from periods of approximately 12 to 48 h, which were present in the LSP. This were also considered artifacts, likely driven by the data upload schedule. All low-density animals showed significant LSP peaks at periods of approximately 45 and 90 min (Fig 5 B).

**Fig 5.**
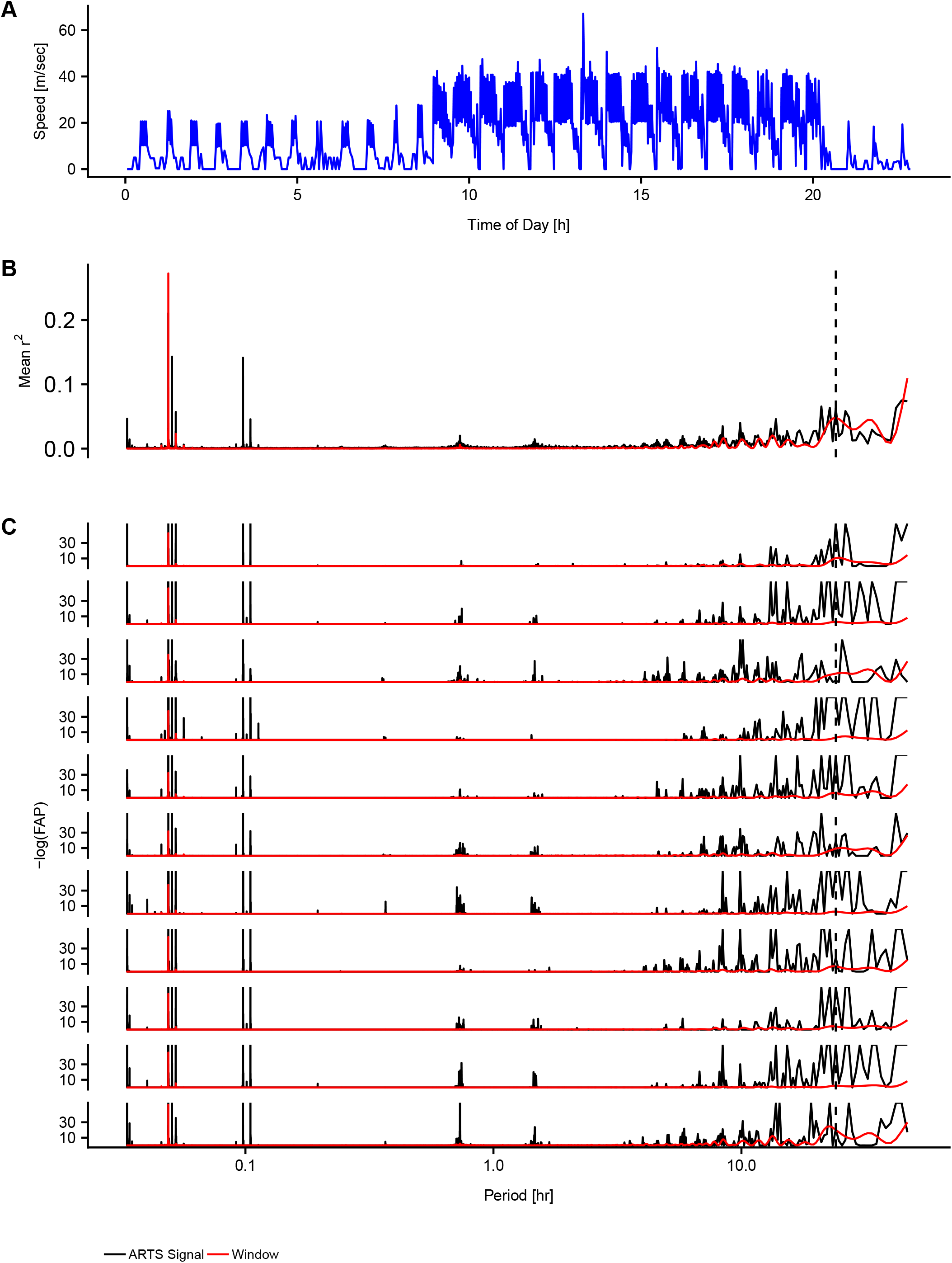
Circadian Activity Patterns Inferred with ARTS. (A) Speed by ARTS for animal 1.508 on 2014-10-10. (B) Mean Periodogram values for speed across animals in the low-density treatment. (C) False-alarm probabilities for the LSP of speed in the low-density treatment over the week of 10-10-2014.

## Discussion

To enable detailed observation of small rodents often studied the lab in natural environments we have deployed a novel ARTS. We used a group of wide-band radio telemetry receivers to enable the rapid estimation of position for ∼50 animals at a time, and explored a variety of alternative approaches to estimating position based on received signal strength. We found that we were able to deploy this system, that it had excellent temporal resolution. Its spatial resolution, though inferior to the precision of manual tracking, was able to provide meaningful and novel observations about the behavior of small mammals in the wild. Below we detail the strengths, limitations and uses of the ARTS system and the analysis tools we have assembled.

Our omnidirectional (isotropic) receiving antennae decayed roughly evenly with distance. We implemented three localization procedures, which we refer to as NLS, Bayesian, and NP trilateration. First, we found that the receiving system was sufficiently sensitive to reliably detect our transmitters across multiple transmitters. Indeed, every antenna covered an area that overlapped with at least three other antennae, and approximately 90% of our mapping positions were detected by at least 3 antennae. Examining RSS as a function of distance demonstrated that it decayed logarithmically with distance and that receiving antenna did not show any strong directionality.

In addition to mapping RSS from a single transmitter, we tracked 48 prairie voles concurrently by manual radio telemetry and ARTS to test the performance of localization algorithms. Data from mapping and tracking were pooled and randomly split into tracking and training data sets, allowing us to combine the extensive coverage from a single transmitter in our mapping data with the more diverse real-world data obtained from traditional manual telemetry. Locations from manual telemetry of live animals, were used as ground truth to build the signal decay model and assess the mean-squared error (MSE) of our models. We found that Bayesian trilateration had the lowest MSE while NP trilateration had the highest. The results of the Bayesian trilateration model were not sensitive to sampling or to the exclusion of outliers from training data.

Our system produced orders of magnitude more localizations (“fixes”) than is possible from manual telemetry. Previously deployed ARTS used directional antenna and narrow-band receivers which must cycle among tag frequencies; in contrast, the Orion broadband receiver can simultaneously sample a wide range of frequencies, allowing a given antenna to sample all tags within a study (48 in our case). This means that ARTS based on narrow-band receivers have been limited in their ability to detect the co-occurrence of tags, as observations are limited in temporal resolution by the time it takes to cycle though tag frequencies. If each tag is observed for one min (eg Steiger et al. 2013), then the best temporal resolution would be one localization per 48 min. Here we observed a median inter-fix interval of only 2 min. This methodological innovation enables the substantial gains in temporal resolution we report.

Compared to previous triangulation-based systems, our ARTS was much more accurate on an absolute scale. The most accurate triangulation-based ARTS we are aware of has reported average localization errors of approximately 30 m (Celis-Murillo et al., 2017; Ward et al., 2014). Our ARTS’ increased spatial accuracy is clearly due to the increased spatial resolution of our antenna array compared the receiving antenna configuration used in other studies. Our study area had a receiving antenna density of 80 antennae/ha and the range between antenna pairs was 9.8 m to 74.4 m. In contrast, previous ARTS implementations have pair-wise antenna distances of hundreds to thousands of meters.

To assess whether the ARTS could detect biologically meaningful variation in space use, we inferred circadian patterns of activity from the Bayesian trilateration predictions. We inferred the animals’ speeds, predicted from the ARTS data, and then used Lomb-Scargle analysis to detect periodic patterns of activity. Prairie voles are often described as nocturnal or sometimes as ultradian (having short activity cycles throughout the day) (Dewsbury, 1980; Grippo et al., 2007; Harper and Batzli, 1996a; Lewis and Curtis, 2016; Taymans et al., 1997). Moreover, laboratory conditions can alter activity cycles of captive rodents and complicate extrapolation to activity cycles in the wild. Because ultradian patterns of activity would be difficult to detect from slower ARTS methods, and impossible to detect from manual telemetry or trapping, this was an excellent means of assessing the utility of our ART system. Our estimates of speed (Fig 5A) revealed that prairie voles had a pronounced ultradian rhythm of roughly 0.75 and 1.5 h, and were also somewhat more active during daylight hours than at night. The circadian rhythm favoring daytime activity in our study was pronounced in some individuals, but not in others, and may contribute to seemingly conflicting data on the circadian cycles of prairie voles. In contrast, the strong ultradian pattern we detected is consistent with laboratory data on heart rate and locomotor behavior (Grippo et al. 2007; Lewis and Curtis 2016) and recent detailed RFID monitoring detecting activity across the day with peak at dusk (Sabol, et. al., 2018). An ultradian rhythm is also consistent with activity patterns common among other species voles, which are thought to eat every 2 to 4 h due to the low nutrient density of forbes and grasses (Tamarin, 1985).

Although our ARTS implementation has a limited ability to localize fixes obtained on a minute-by-minute basis, our data suggest several avenues for productive use. First, on longer time-scales, many fixes can be averaged, allowing a sliding-window kernel density estimate that can track changes in home-range or other aspects of space use on a scale of hours or days – a scale not plausible with manual methods which require weeks of data to fit. Second, by employing conservative detection criteria, the antennae can precisely localize when one or more individuals is present at an antenna, functioning as an elaborate RFID array. Indeed, the WILDSENSING aRFID system used to track badgers [*Melees melees*] operates on this principle (Dyo et al., 2012; Ellwood et al., 2017). Social networks (Ellwood et al., 2017; Smith et al., 2018; Sabol et al, 2018), pair-bonding (Psorakis et al., 2012; Sabol et al, 2018) and territory defense (Perony et al., 2012) can be inferred from such co-occurrence data. By placing antennae above a feeder, or at nest sites, one could combine the strengths of RFID measures with those of telemetry. These kinds of data would be useful for characterizing temporal patterns of activity, as we demonstrate in our assessment of ultradian cycles. Such data have uses well beyond activity measurement – allowing researchers to examine, for example, the correlated movement patterns of pairs or social groups, or the time an animal spends on a nest. Indeed, one unique strength of this system is that these approaches all rely on post-collection data processing, and so are not mutually exclusive. The relationships among, say, the temporal activity correlations of two animals, the extent of their home-range overlap, and the rate at which those same two animals are present together at an antenna each offers a distinct perspective on the nature and extent of the pair’s interactions. The ability to compare such data directly will allow a fuller characterization of social interactions in the wild and offer more reliable and detailed assessment of such behaviors than is currently available. In addition, the high temporal resolution of the data may allow detection of unique patterns of activity that are characteristic of specific behaviors, analogous to inferences made from data using accelerometers and automated pose estimation for example (Chambers et a. 2020, Pereira et al. 2019).

One avenue for future development of this and other ARTS systems is to replace trilateration with machine-learning based methods. Random forest classifiers, for example, have been used to develop “location fingerprints” that identify an animal’s position based on a unique combination of received signal strengths (Harbicht, et. al., 2017). Similarly, the structure of our data, with an array of received signal strengths for each transmitter, seem well suited for feedforward neural network methods. Our results with the nonparametric method, however, suggest that care needs to be taken when implementing such tools to avoid overfitting the training data.

In summary, we deployed a new ARTS using ultra-wideband receivers that are capable of simultaneously tracking dozens small animals with high temporal resolution. We show that a simple Bayesian trilateration procedure performs well, and that its limitations reflect the limitations of the hardware. We have validated the ability of the ARTS to track biologically meaningful behavior by using the ARTS to detect circadian activity patterns. By combining the various means of processing ARTS data, we see an exciting range of opportunities to use this system to enrich the ecological and mechanistic study of behavior.

## Supporting information

Analysis R Code

## Acknowledgements

We would like to acknowledge Ron Pulcher for the extensive guidance and physical assistance with field site maintenance and Cameron Grant from Sigma Eight Inc for assistance with Orion receiver setup and data recovery.

## Authors’ Contributions

*GW and SP designed the study; GW, MG and RS collected the data; GW performed the analysis and preparation the manuscript. SP contributed critically to the analysis and drafts. All authors approve publication*.

## Data availability

Data will be archived on Dryad before publication.

