## Supplement for "An Automated Radio-Telemetry System (ARTS) for Monitoring Small Mammals"

2021-03-06

#### Methods

#### Signal Decay Model

An idealistic model of signal decay is given by the Friss equation (Eqn 1) (Friis, 1946) . This model assumes an isotropicly radiating electromagnetic signal propagating in free-space.

$$\begin{matrix} & RSS=P_{t}+20*\log_{10}(\frac{\lambda}{4\pi d}) & (1) \\ & Where: & \\ & RSS=received signal strength & \\ & P_{t}=transmitter power & \\ & \lambda=signal wavelength & \\ & d=distance between transmitter and receiver & \end{matrix}$$

In practice the path-loss exponent differs from the free-space value of 2 due to the propagation environment. However this may be accounted for using regression to estimate an empirical decay coefficient (Eqn 2)(Goldsmith, 2005). Once values of $\beta_{0}$ and $\beta_{1}$ are the distance between the transmitter and antenna may be estimated (eq 3).

$$\begin{matrix} & RSS=\beta_{0}+\beta_{1}*10\log_{10}(d/d_{0})+\epsilon& (2) \\ & d=d_{0}{10}^{\frac{RSS-\beta_{0}}{10\beta_{1}}} & (3) \\ & Where: & \\ & d_{0}=reference distance & \\ & \beta_{0}=power received at d_{0} & \\ & \beta_{1}=empiracal path-loss exponent & \\ & \epsilon=gaussian noise & \end{matrix}$$

The relationship between distance and RSS was modeled using hierarchical Bayesian regression in JAGS (Plummer, 2003). Training data and model specification were exported from R to JAGS using the packages “rjags” (Plummer, 2016). Antenna were hierarchically clustered within their shared receiver. This is because the selection within an antenna switch is not perfect. Particularly strong signals may “bleed through” within the switch causing correlation between antenna on the same switch. Received signal strength was modeled as a linear function of the logarithm of distance with normally distributed noise (Eqn 4). Slopes and intercepts were drawn for each antenna from multivariate normal prior unique to each receiver (Eqn 5). The mean vector of each multivariate normal distribution for each receiver was drawn from a multivariate normal prior (Eqn 6). Each antenna’s RSS precision was drawn form a gamma prior (Eqn 7). The precision matrix was drawn from a Wishart prior (Eqn 8). The posterior distribution of distance given RSS for each antenna was drawn from a lognormal distribution (Eqn 9). The JAGS model was specified in the supplemental “model.4.txt” script.

$$\begin{matrix} & Model: & \\ & RRS_{i}\sim N(\beta_{0j}+\beta_{1,j}*10\log_{10}(d),\tau_{j}^{2}) & (4) \\ & \beta_{j}\sim MVN(\mu_{k},\Sigma^{-1}) & (5) \\ & Priors: & \\ & \mu_{k}\sim MVN(\mu_{0},\Sigma_{0}^{-1}) & (6) \\ & \tau_{j}^{2}\sim gamma(a_{0},b_{0}) & (7) \\ & \Sigma^{-1}\sim Wish(V_{0},n_{0}) & (8) \\ & Posterior Predictive: & \\ & d\sim logN\left( \frac{\log(10)(RSS_{i}-\beta_{0j})}{10\beta_{1}j},|\frac{\log(10)}{10\beta_{1j}*\tau_{j}}| \right) & (9) \\ & Where: & \\ & i=observations index & \\ & j=antenna index & \\ & k=receiver index & \\ & \tau_{j}=precision at antenna j & \\ & \beta_{j}=parameter vector containing \beta_{0}, \beta_{1}for antenna j & \\ & \Sigma^{-1}=precision matrix for parameter vector \beta_{j} & \\ & \{\mu_{0},\Sigma_{0}^{-1},V_{0},n_{0},a_{0},b_{0}\}=\mathrm{Hyperparameters} & \end{matrix}$$

**Nonlinear Least Squares Trilateration**

Trilateration is based on the distance between a transmitter and the receiving antenna. When a signal is received at an antenna a signal-decay model is used to project a circle around that antenna, with a center at the antenna and with radius determined by the signal decay model. The transmitters will be located at the intersection of the projected circles (Fig 2A). If antenna positions and distances are perfectly known, then the intersection of these three circles can be solved algebraically. In the presence of measurement error these circles will not uniquely intersect. It is possible to estimate the transmitter position by ordinary least-squares or nonlinear least-squares regression. The nonlinear least-squares solution was used here, as it is significantly more accurate than ordinary least-squares solution (Navidi et al., 1998). The nonlinear least-squares finds the point which minimizes the sum-of-squares of estimated distances (Eqn 10).

$$\begin{matrix} & \overset{^}{xy}=ArgMin\sum_{i=1}^{3} \left[ \overset{^}{d_{i}}-\sqrt{(x-x_{i})^{2}+(y-y_{i})^{2}} \right]^{2} & (10) \\ & Where: & \\ & \overset{^}{xy}=position estimate (Easting, Northing) & \\ & i=antenna index & \\ & \overset{^}{d_{i}}= estimate of distance between antenna i and transmitter & \\ & x_{i}=antenna i Easting & \\ & y_{i}=antenna i Northing & \end{matrix}$$

At each time point signal from the three antennas with the largest RSS were selected. For each antenna distance, $\overset{^}{d_{i}}$, was calculated using Eqn 5, implied from the empirical formula for RSS decay. Parameters $\beta_{0j}$ and $\beta_{1j}$ were taken from the expectation of the posterior distribution of $\beta_{j}$ obtained from the Bayesian signal decay model. The parameter estimate of position was obtained by minimizing equation 10, using the Levenberg–Marquardt algorithm in the R package “minpack.lm” (Elzhov et al., 2016). Observations were weighted according to $\frac{1}{d_{i}^{2}}$. The algorithm was allowed up to 100 iterations to reach covergence. Nonlinear least squares trilateration was implemented in the Trilaterate function in the “ARTSFunction.R” file.

#### Nonparametric Trilateration

To model signal decay without the assumption of isotropy implicit in the signal decay model nonparametric KDEs of the probability of position were fit for each antenna. Multivariate-normal kernels were used to smooth over three dimensions (Northing, Easting, and RSS) within the study area for each antenna. Densities were evaluated across a 1 x 1 m resolution grid spanning the enclosure. Kernels for the probability of location given RSS were estimated from the training dataset using the R package “ks” (Duong, 2018). Bandwidth matrices were automatically selected using the multivariate plug-in selector of Wand and Jones (1994) implemented in the Hpi function. Kernels for each antenna were fit with the function “CalcTablesNP” in the “ARTSFunction.R” file.

#### Bayesian and Nonparametric “Trilateration”

At each time point signal from the strongest three distinct antennae were selected. For the Bayesian model at each antenna the posterior pdf of distance obtained from the signal decay model was projected onto a 1 x 1 m resolution grid spanning the study area. To localize the transmitter using Bayesian and Nonparametric models the signal at each antenna was assumed to be independent. Thus, the joint probability of position given the signal at each antenna should be the product of the densities at each antenna (Eqn 11). The point estimate of position was taken to be the maximum of the element-wise product of pdfs across selected antenna at the corresponding RSS (Eqn 12, Fig 2B & 2C). Products were normalized to one.

$$\begin{matrix} & Assuming independent antenna & \\ & Pr(xy|RSS_{1:3})=\prod_{i=1}^{3} Pr(xy|RSS_{i}) & (11) \\ & \overset{^}{xy}=ArgMax[Pr(xy|RSS_{1:3}))] & (12) \\ & Where: & \\ & i=antenna index & \end{matrix}$$

### Circadian Inference

Periodic patterns in time series are often analyzed via Fourier analysis, which applies the Fourier transform to a signal in the time domain and yields the signal power in the frequency domain. Real world signals are analyzed using the discrete Fourier Transform. This means that the analyzed signal is actually a sample of the continuous-time signal (ie the convolution of the continuous signal and the Dirac comb sampling window). Because the signal of interest is convolved with a sampling window the Fourier transform also reflects a substantial amount of correlated noise when the sampling function is irregular. Because the sampling functions take the value of zero at all unobserved points it becomes impossible to deconvolve irregularly sampled signals with the discrete Fourier transform. Unfortunately, due to logistical and technical challenges most animal behavior data recorded in the field will not be uniformly sampled.

The Lomb-Scargle Periodogram (LSP) can approximate the Fourier transform for irregularly sampled data (Lomb, 1976; Scargle, 1982). The LSP has been a useful tool for detecting circadian behavior in the lab and field and has previously been applied to the analysis of ARTS data (Glynn et al., 2006; Péron et al., 2016; Ruf, 1999; Steiger et al., 2013; Zielinski et al., 2014).

The LSP is equivalent to fitting a linear model consisting of a sine and a cosine term to the signal for each frequency in consideration (Eqn 13). Once the model is fit, the periodogram spectral density (PSD) is calculated from the $\chi^{2}$ goodness-of-fit statistic (Eqn 14) and normalized to the $\chi^{2}$ goodness-of-fit statistic for null hypothesis (eg. the sample variance) resulting in a power that is equivalent to the model $r^{2}$ (Eqn 15). Frequencies may be scanned from 0 Hz to $f_{max}$ without aliasing. For irregularly sampled data $f_{max}$ is equal to half of the frequency of the greastest common factor of sampling intervals, which may be much greater than the commonly used half of the frequency of the average sampling interval (VanderPlas, 2017). The normalized LSP power was directly from time series data (Eqn 16) as this is faster than computing the liner model $\chi^{2}$ goodness-of-fit statistic at each frequency (Zechmeister and Kürster, 2009). The normalized LSP power was calculated using the function GenLS in the “ARTSFrunctions.R” file.

$$\begin{matrix} & \overset{^}{y}(f,t_{i})=A\cos\omega(t_{i}-\tau)+B\sin\omega(t_{i}-\tau)+c & (13) \\ & Where: & \\ & i=observation index & \\ & t_{i}=time at observation i & \\ & \overset{^}{y}(f,t_{i})=signal estimate at time t & \\ & f=\mathrm{frequency} & \\ & \omega=2\pi f & \\ & A=sine term amplitude parameter & \\ & B=cosine term amplitude parameter & \\ & c=intercep parameter & \\ & \tau=phase offset parameter & \end{matrix}$$

$$\begin{matrix} & P(f)=\frac{1}{2}(\chi_{0}^{2}-\chi^{2}(f)) & (14) \\ & P(f)_{normalized}=\frac{\chi_{0}^{2}-\chi^{2}(f)}{\chi_{0}^{2}} & (15) \\ & Where: & \\ & P(f)=Lomb-Scargle PSD & \\ & \chi^{2}(f)=model Goodness-of-Fit=W\sum_{i=1}^{n} w_{i}(y-\overset{^}{y})^{2} & \\ & \chi_{0}^{2}=null Goodness-of-Fit=W\sum_{i=1}^{n} w_{i}(y_{i}-\overline{y})^{2} & \\ & w_{i}=observation weights=\frac{1}{W\sigma_{i}^{2}} & \\ & W=\sum1/\sigma_{i}^{2} & \\ & \sigma_{i}^{2}=gaussian noise & \end{matrix}$$

$$\begin{matrix} & P(f)_{normalized}=\frac{1}{YY}[\frac{YC_{\tau}^{2}}{CC_{\tau}}+\frac{YS_{\tau}^{2}}{SS_{\tau}}] & & & (16) \\ & \mathrm{note}\tau subscript indicates times shifted by \tau& & & \\ & Where: & & & \\ & Y=\sum w_{i}y_{i} & & & \\ & C=\sum w_{i}y_{i}\cos(\omega t_{i}) & & & \\ & S=\sum w_{i}y_{i}\sin(\omega t_{i}) & & & \\ & YY=\overset{^}{YY}-Y\cdot Y & \overset{^}{YY} & =\sum w_{i}y_{i}^{2} & \\ & YC=\overset{^}{YC}-Y\cdot C & \overset{^}{YC} & =\sum w_{i}y_{i}\cos(\omega t_{i}) & \\ & YS=\overset{^}{YS}-Y\cdot S & \overset{^}{YS} & =\sum w_{i}y_{i}\sin(\omega t_{i}) & \\ & CC=\overset{^}{CC}-C\cdot C & \overset{^}{CC} & =\sum w_{i}\cos^{2}(\omega t_{i}) & \\ & SS=\overset{^}{SS}-S\cdot S & \overset{^}{SS} & =\sum w_{i}\sin^{2}(\omega t_{i}) & \\ & CS=\overset{^}{CS}-C\cdot S & \overset{^}{CS} & =\sum w_{i}\cos(\omega t_{i})\sin(\omega t_{i}) & \\ & tan(2\omega\tau)=\frac{2CS}{CC-SS} & & & \end{matrix}$$

The false-alarm probability (FAP) is the probability that an observed power in the scanned frequency range was generate by chance (Eqn 17). The null distributions for hypothesis tests of LSPs cannot be calculated exactly for irregularly sampled data and require either approximation or calculation by computationally intensive bootstrapping. Baluev (2008) approximated the Davies limit of upper-bound of p-values for the FAP (Note that Baluev’s parameterization of the null normalized LSP power differs by a factor of $\frac{2}{df}$ from Cumming et al. (1999) and Zechmeister and Kürster (2009))(Eqn 18). This approximation is quick to compute and in good agreement with bootstrapping form uniformly sampled data. Further, it tends to produce conservative, slightly increased, FAP estimates compared to bootstrapping for irregularly sampled data (Baluev, 2008; VanderPlas, 2017).

$$\begin{matrix} & FAP=1-(1-Pr(Z<z_{obs}))^{m} & (17) \\ & Max(FAP)\approx1-Pr(Z<z_{obs})e^{-\tau} & (18) \\ & Where: & \\ & Pr(Z<z_{obs})=1-(1-Z)^{df/2} & \\ & \tau=\gamma A(f_{max})(Z)^{\frac{N-df-2}{2}}(1-Z)^{\frac{df-1}{2}} & \\ & & \\ & \gamma=\frac{1}{2\pi}\frac{\Gamma(\frac{N-1}{2})}{\Gamma(\frac{df}{2})}\left( \frac{1}{\pi} \right)^{\frac{N-df-2}{2}} & \\ & & \\ & A(f_{max})=2\pi^{3/2}f_{max}\sqrt{4\pi Var(t)}) & \\ & & \\ & Z=normalized Lomb-Scargle Spectral Density & \\ & t=time, & \\ & f_{max}=upper frequency limit & \\ & m=number of independent frequencies scanned & \\ & z_{obs}=observed Lomb-Scargle periodogram power & \\ & \alpha=acceptible Type-I Error Rate & \\ & df=degrees of freedom & \end{matrix}$$

Patterns of circadian behavior in freely behaving voles were inferred over the second week of manual radio telemetry from 2014-10-07 00:00:00 through 2014-10-13 23:59:00. Lomb-Scargle periodograms for speed and distance from nest site were computed. Generalized Lomb-Scargle periodograms were calculated in R following the algorithm layed out in Zechmeister and Kürster (2009). The scanned frequency grid ranged from 1.65310^{-6} Hz to 0.008333 Hz with an oversampling factor of five. The Davies upper bound approximation of p-values (Baluev, 2008) was calculated in R, following VanderPlas (2017) in the FAP_davies function in the “ARTSFunction.R” file. Sampling windows were calculated via the discrete Fourier Transform.

### Software

Data were analyzed in R (R Core Team, 2018). In addition to the packages mentioned above the following R packages were used. The packages “data.table”, “dplyr”, and “dtplyr” (Dowle and Srinivasan, 2018; Wickham, 2017; Wickham et al., 2017) were used for general data manipulation and joins. Dates and times were formated with “lubridate” (Grolemund and Wickham, 2011). Plots were generated with “ggplot2”, “akima”, “ggforce”, “GGally”, and “cowplots” (Akima and Gebhardt, 2016; Pedersen, 2016; Schloerke et al., 2017; Wickham, 2009; Wilke, 2017). Contour geometry for KDEs and PDFs were evaluated using “sp” and “rgeos” (Bivand and Rundel, 2017; Pebesma and Bivand, 2005).

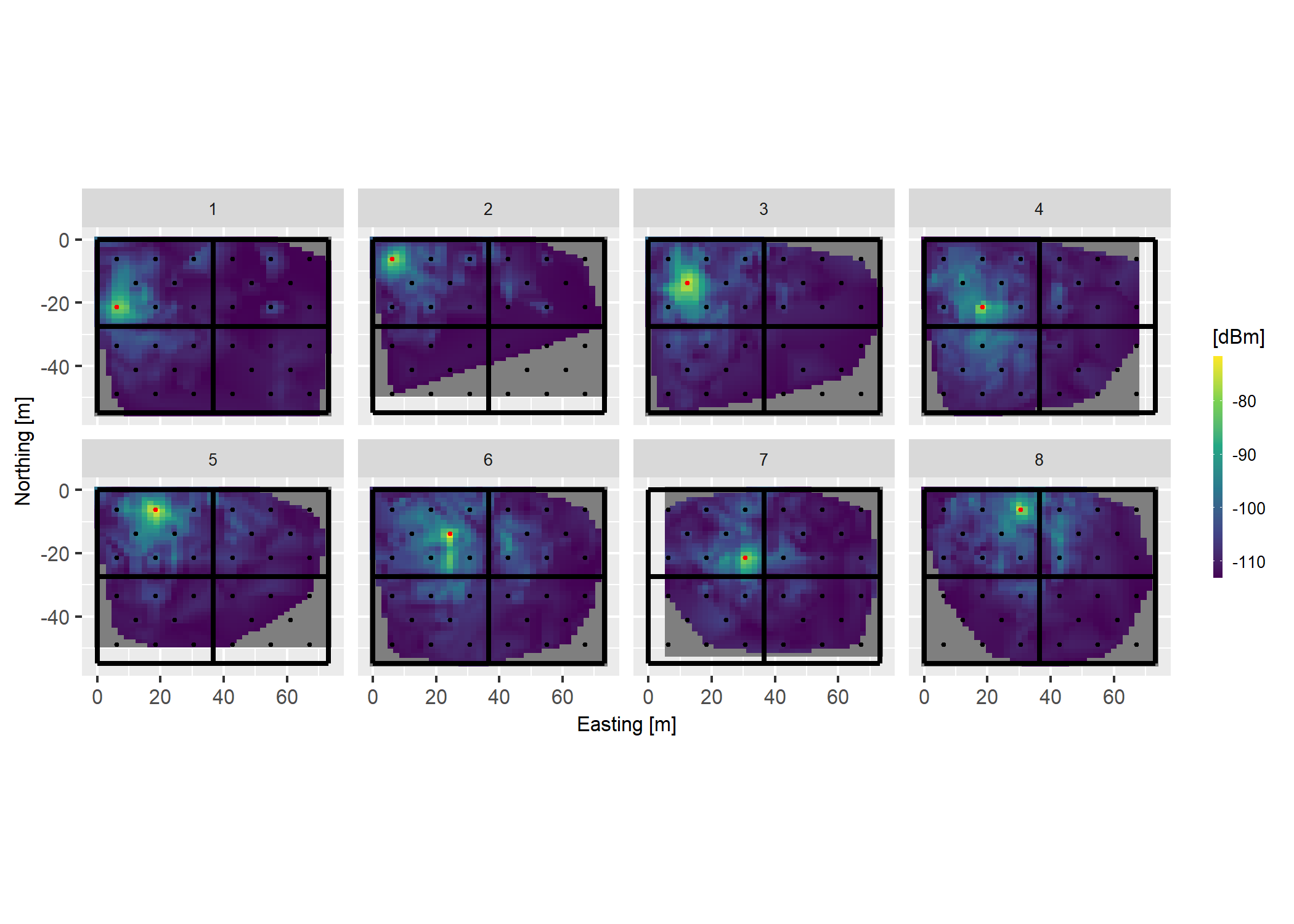

**Fig. S1. Antenna Receptive Fields for Quadrant 1 (NW).** Antenna one through eight linear interpolation of RSS in the radiomapping dataset. Red indicates focal antenna.

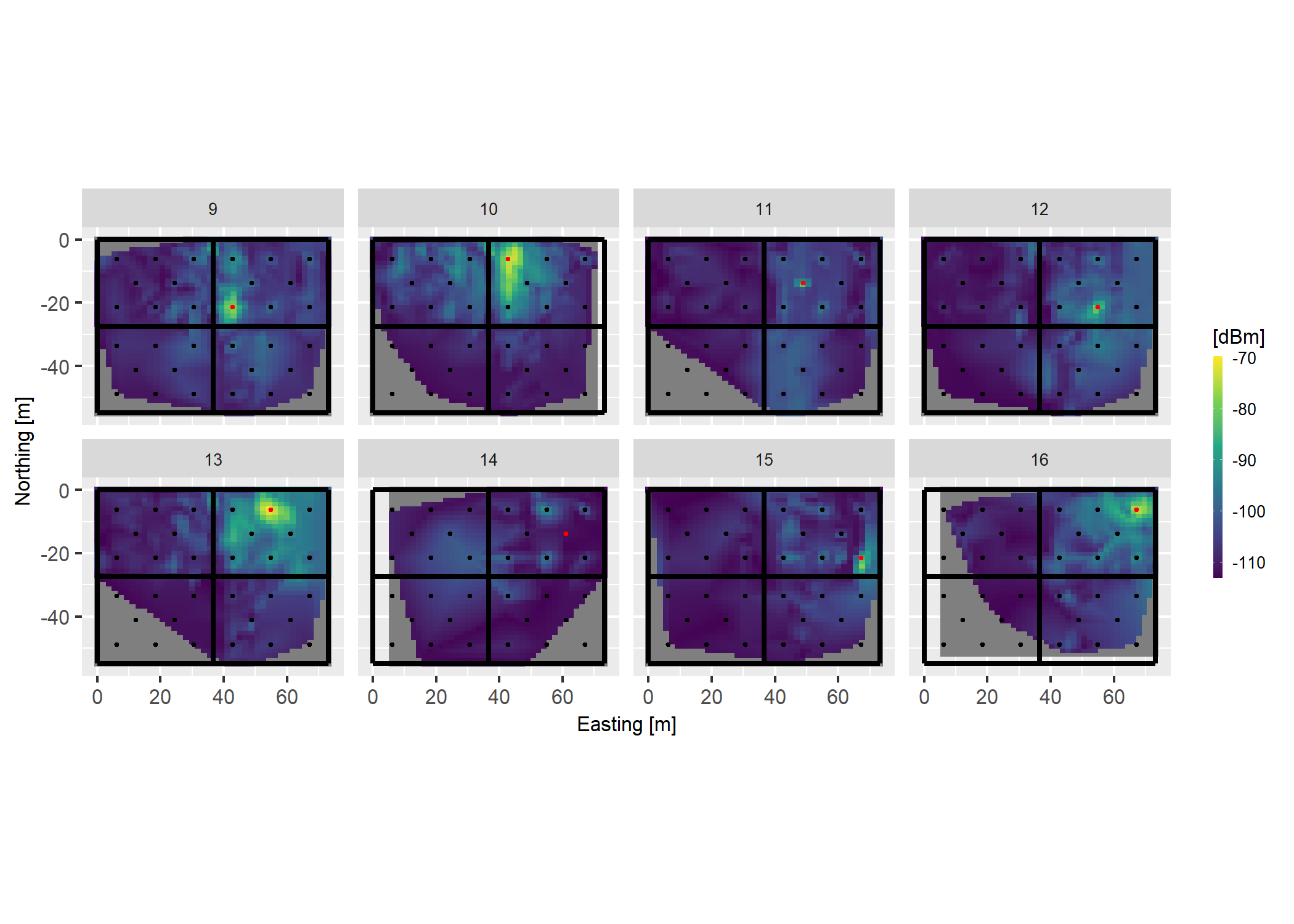

**Fig. S2. Antenna Receptive Fields for Quadrant 2 (NE).** Antenna nine through sixteen linear interpolation of RSS in the radiomapping dataset. Red indicates focal antenna.

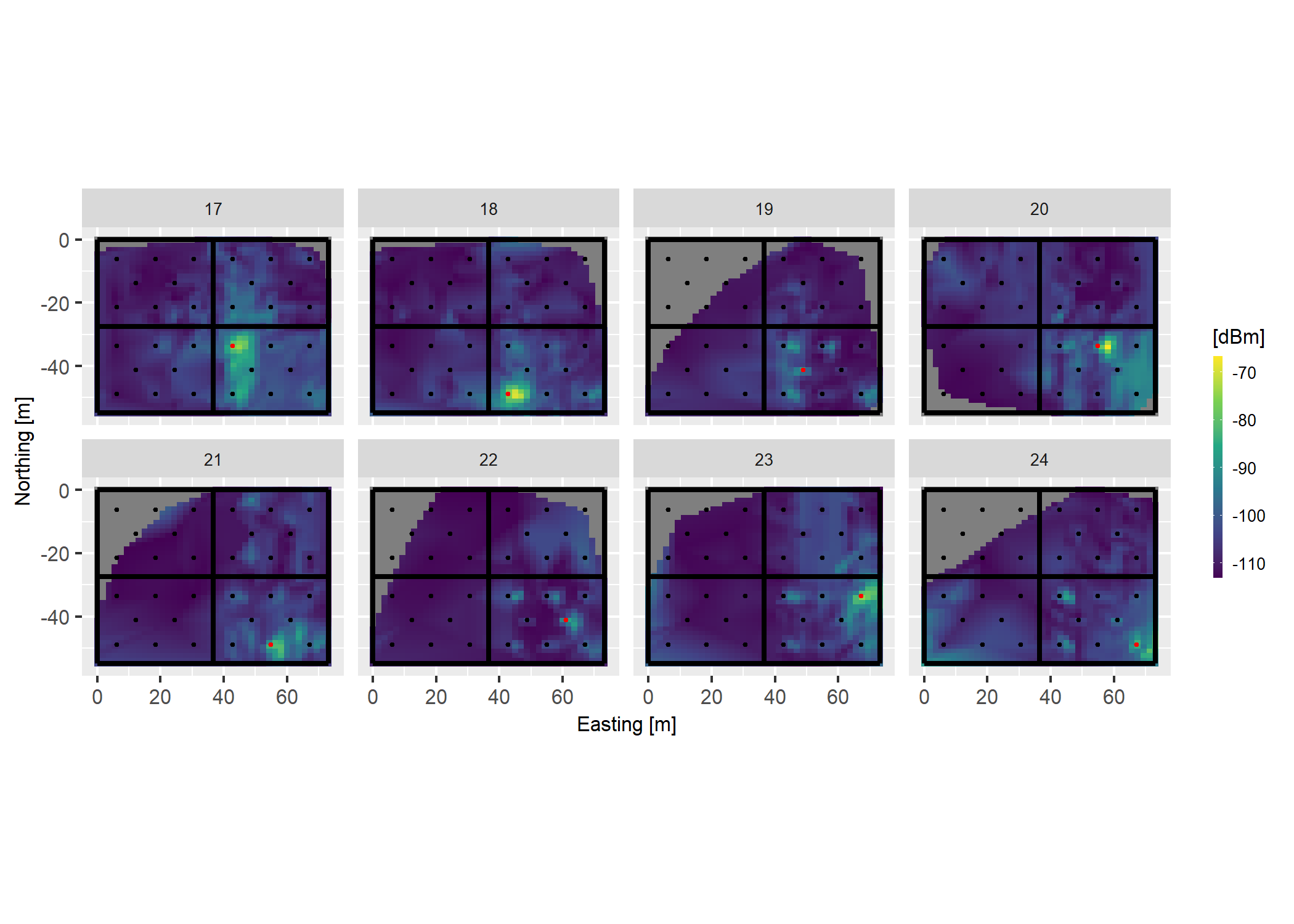

**Fig. S3. Antenna Receptive Fields for Quadrant 2 (SE).** Antenna 17 through 24 linear interpolation of RSS in the radiomapping dataset. Red indicates focal antenna.

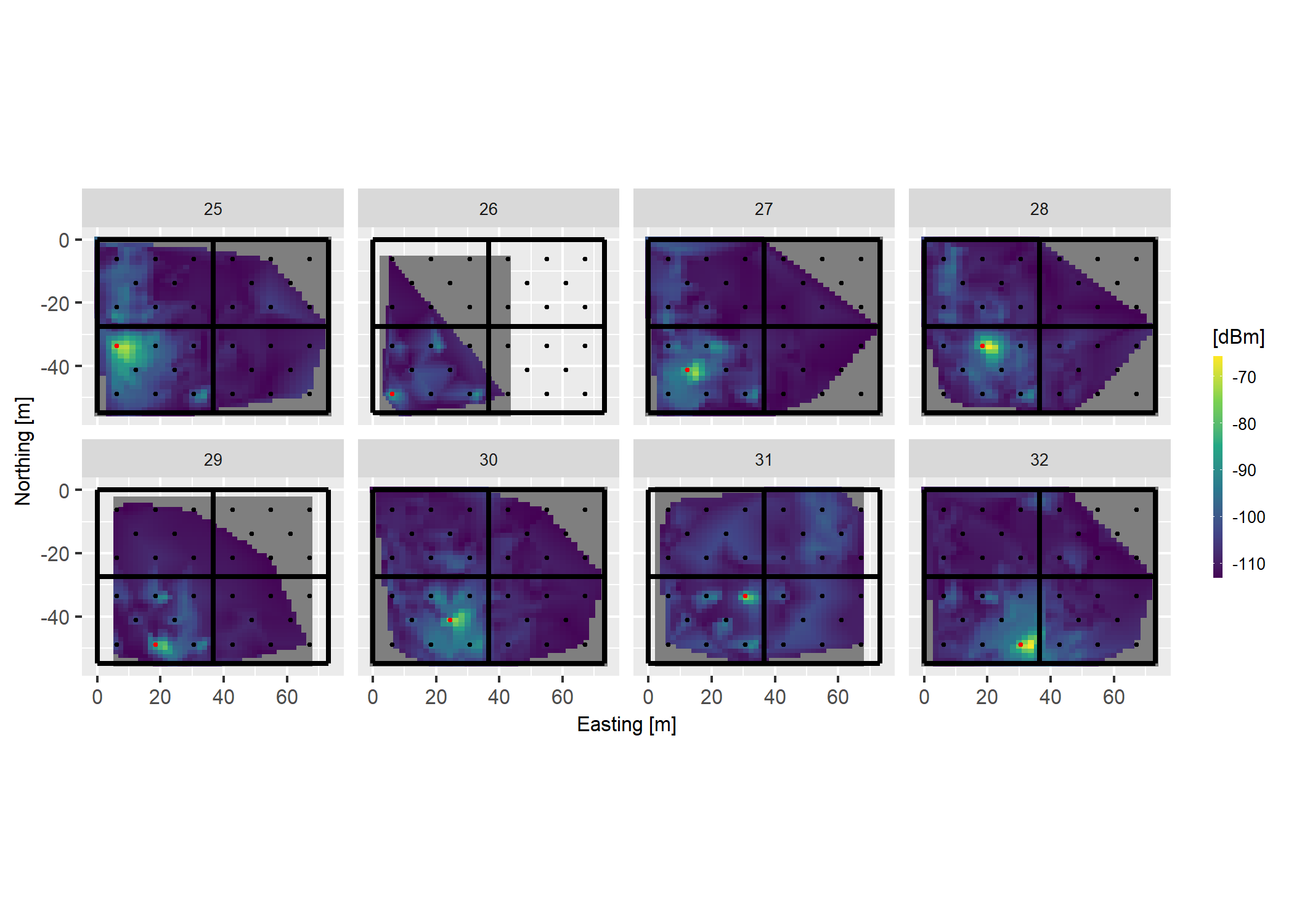

**Fig. S4. Antenna Receptive Fields for Quadrant 2 (SE).** Antenna 25 through 32 linear interpolation of RSS in the radiomapping dataset. Red indicates focal antenna.

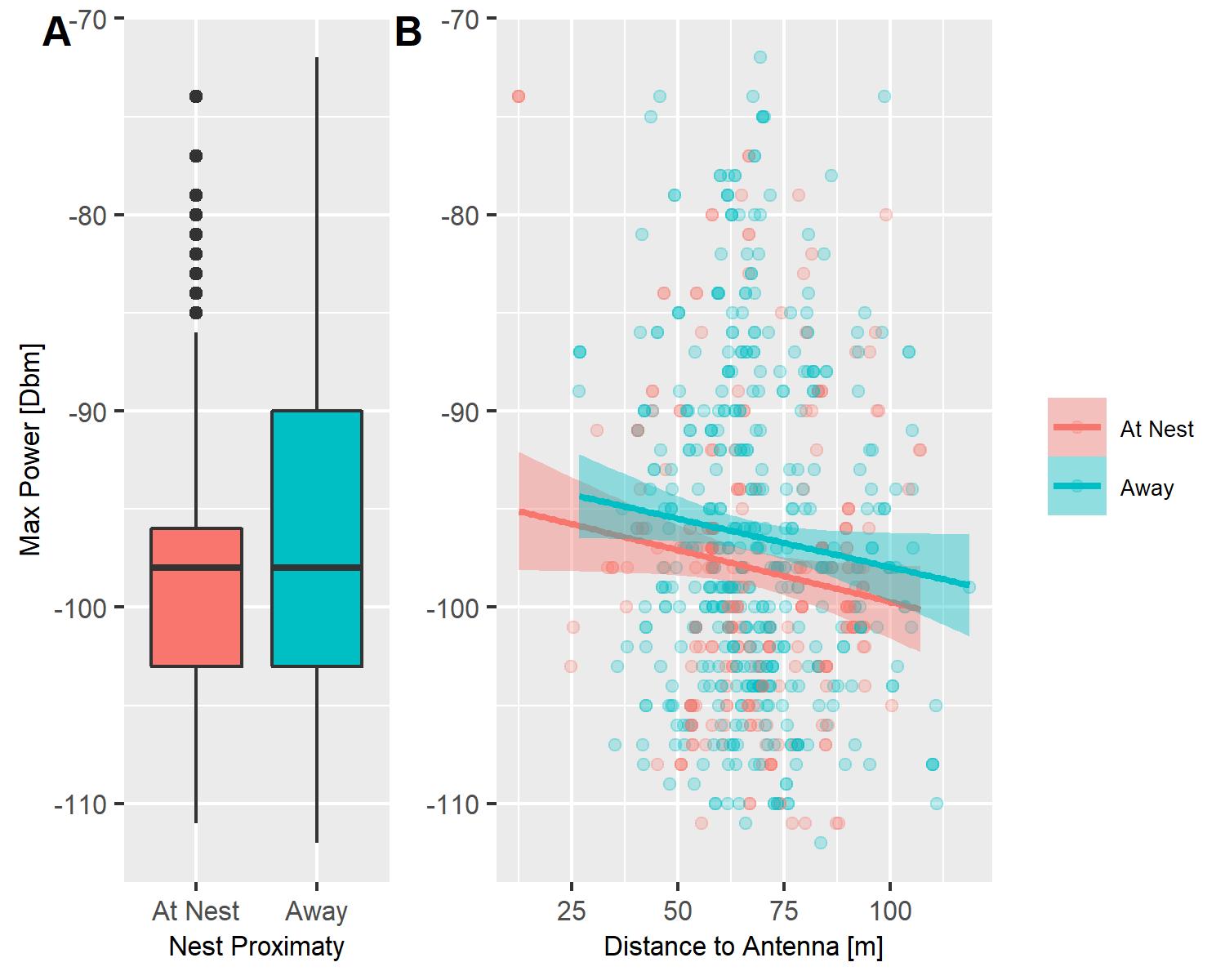

**Fig. S5. Effect of Nest Proximaty on RSS.** Maximum received power per fix was calcuated. (A) Power near and away from the estimated nest site (maxima of manual trackingKDE). (B) Power near and way from the estimated nest site accounting for distance to the antenna receiving the strongest sign in a given fix.

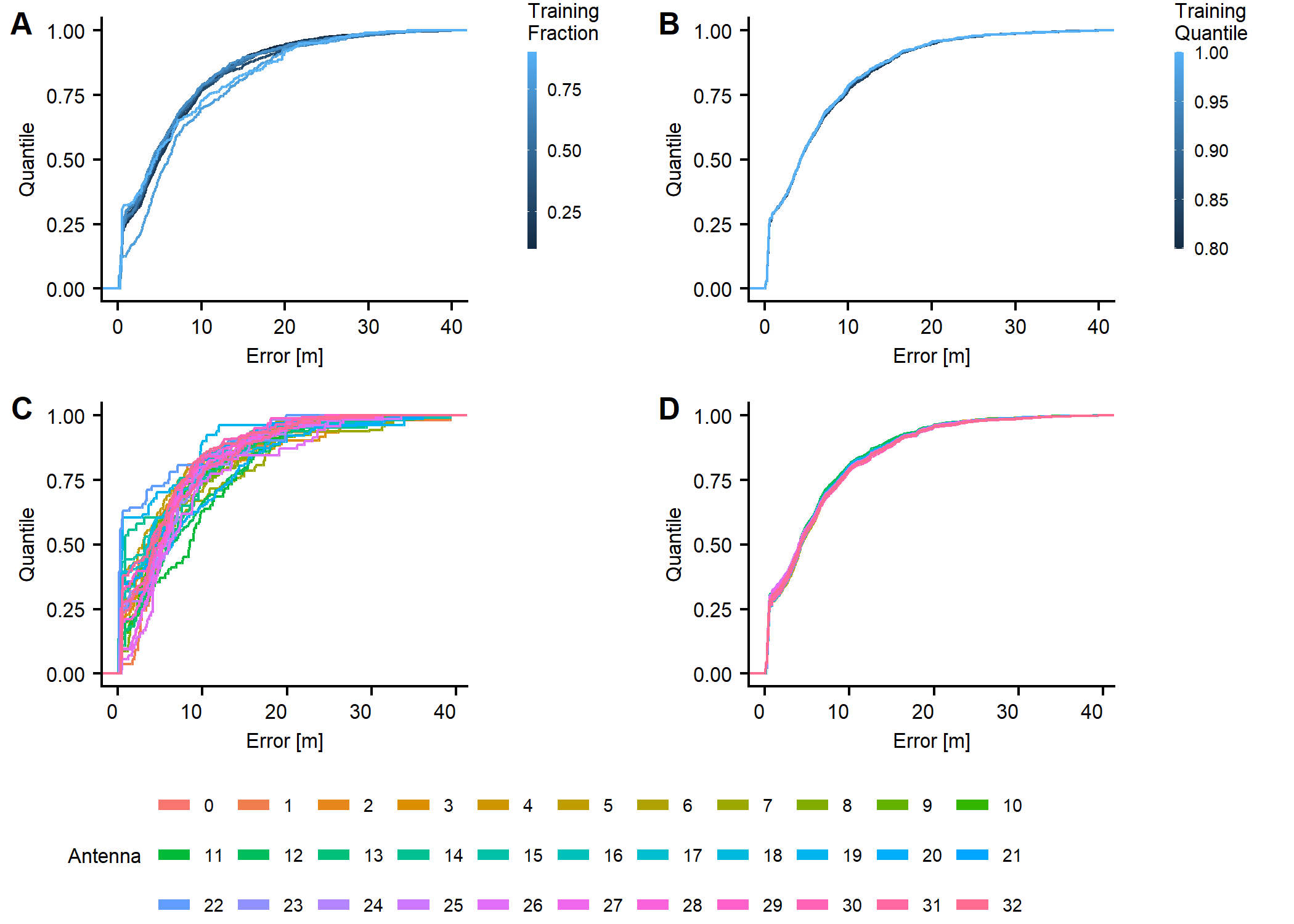

**Fig. S6. Fix Accuracy: Sensitivity to Sample, Outliers, and Antenna.** (A) Empirical cumulative density of error for the Bayesian trilateration model trained on random samples from 10% to 90% of manual fixes and tested on the remaining 90% to 10% of manual fixes. (B) Empirical cumulative density of error for the Bayesian trilateration model trained by excluding points in the 0.8 to 0.99 quantile of the error distribution. (C) Empirical cumulative density of error for fixes including a given antenna (D) Empirical cumulative density of error for fixes excluding each antenna.

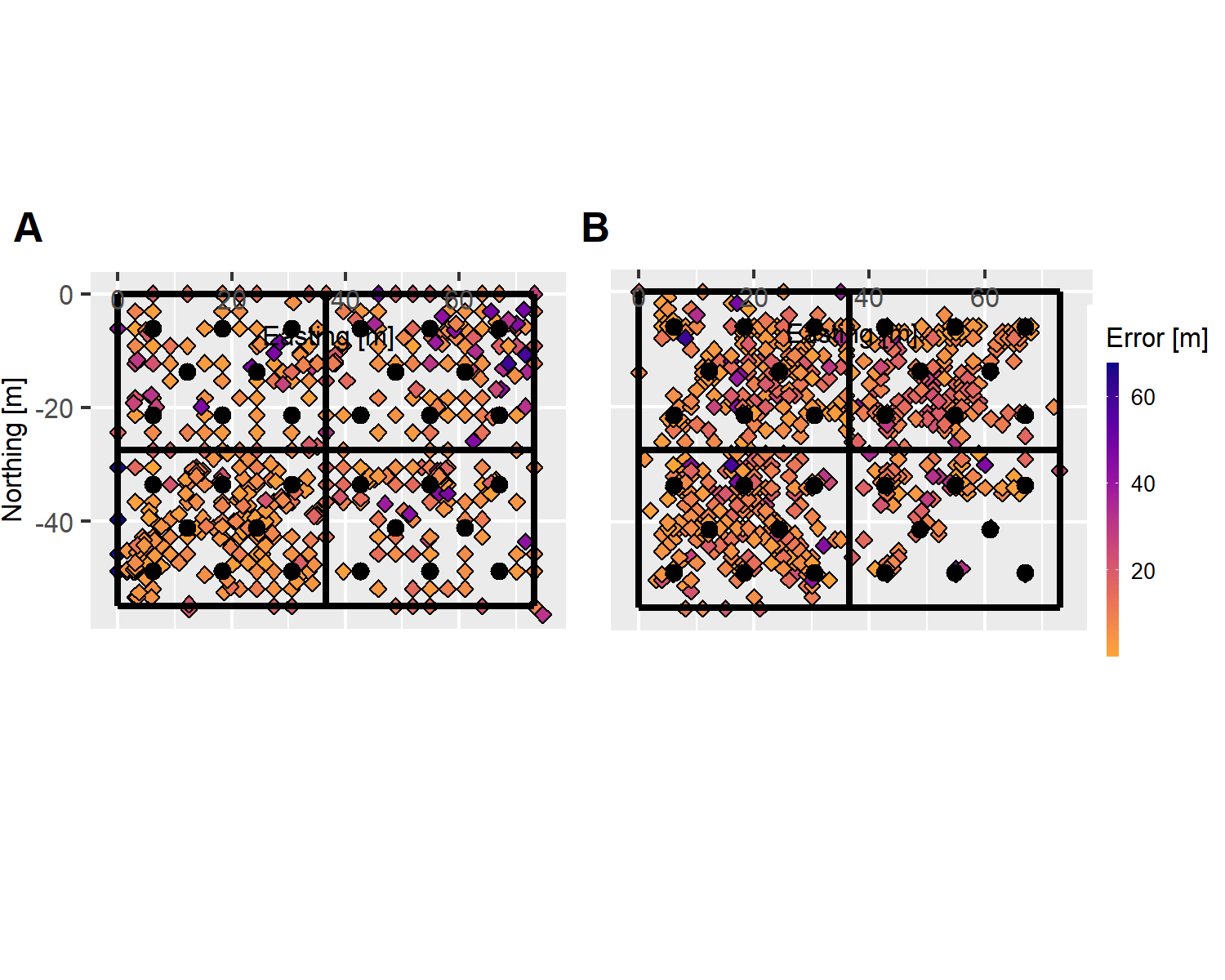

**Fig. S7. Bayesian Trilateration Accuracy as a Function of Space.** The colormap represents euclidean distance between know transmitter location and location predicted by Bayesian trilateration (A) Points represent known relocations coordinates from the testing data set. (B) Points represent coordinates predicted by Bayesian trilateration.

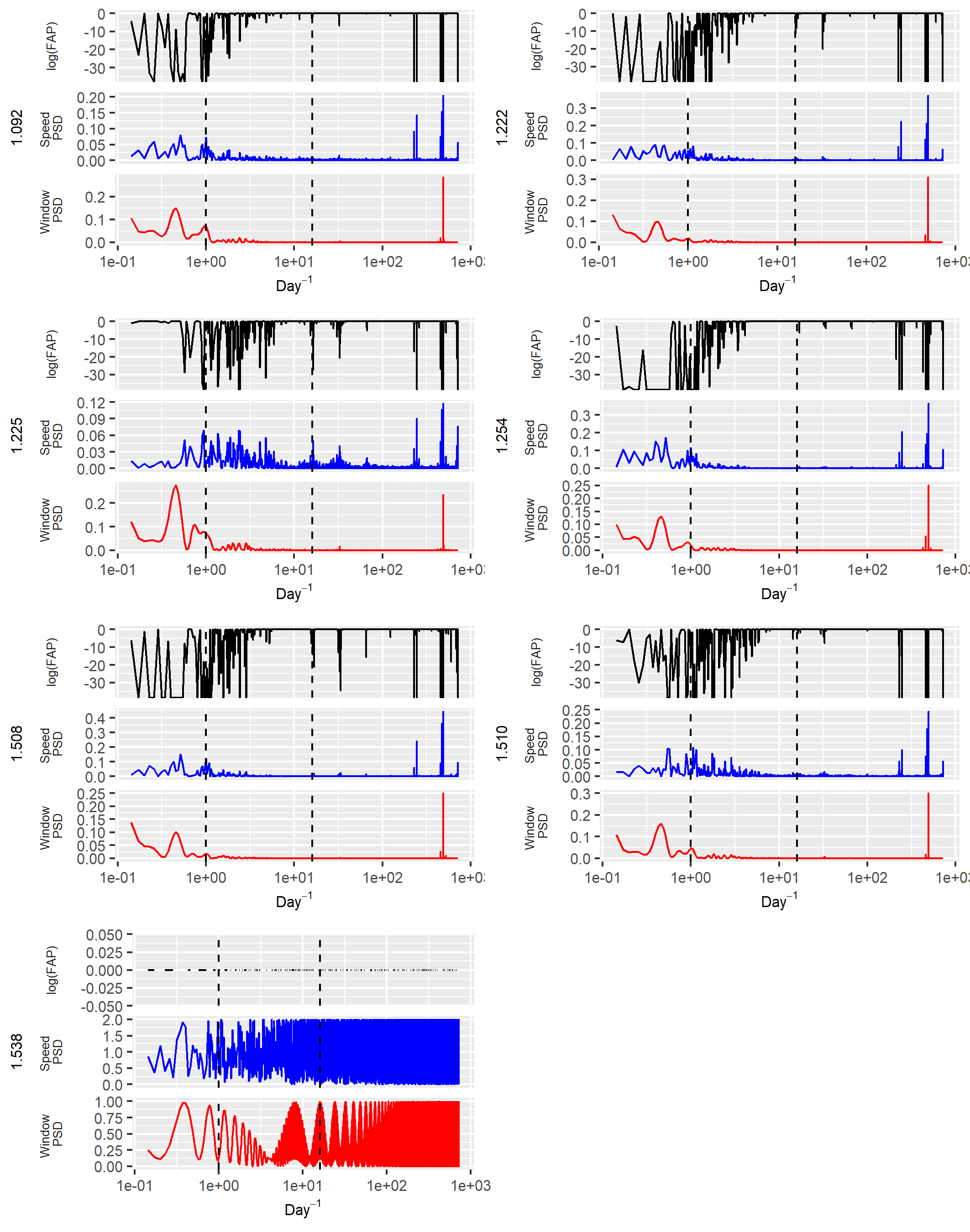

**Fig. S8. Lomb-Scargle Periodograms for Low Density Males** Lomb-Scargle Periodograms of speed were calculated from Bayesian trilaterion predictions over the week of 10-10-2014. Dashed lines indicate periods of 24 hours and 90 min. Animal 1.538 only had 3 observations and was excluded from further analysis (Note the sampling window to dominate the LSP).

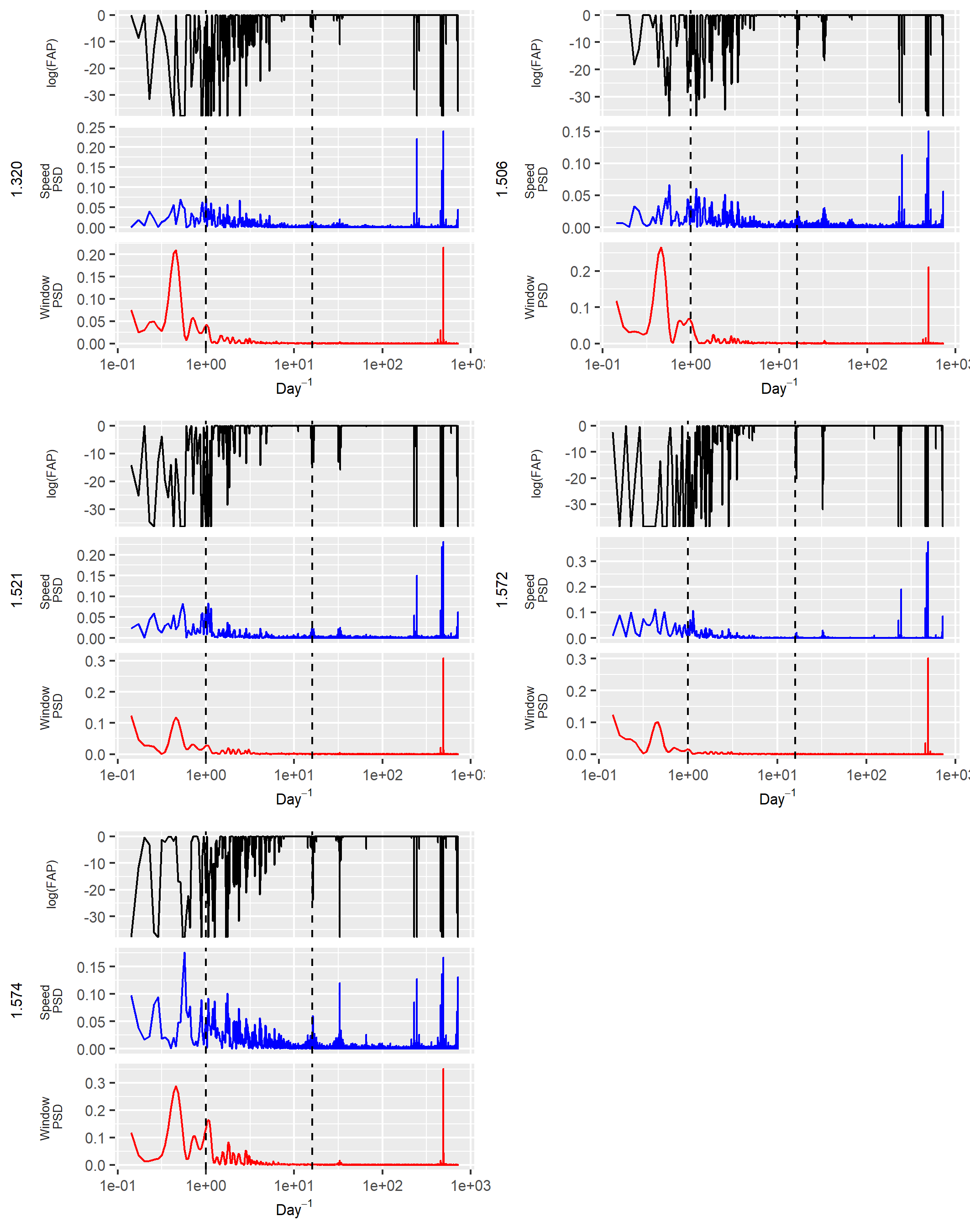

**Fig. S9. Lomb-Scargle Periodograms for Low Density Females** Lomb-Scargle Periodograms of speed were calculated from Bayesian trilaterion predictions over the week of 10-10-2014. Dashed lines indicate periods of 24 hours and 90 min.

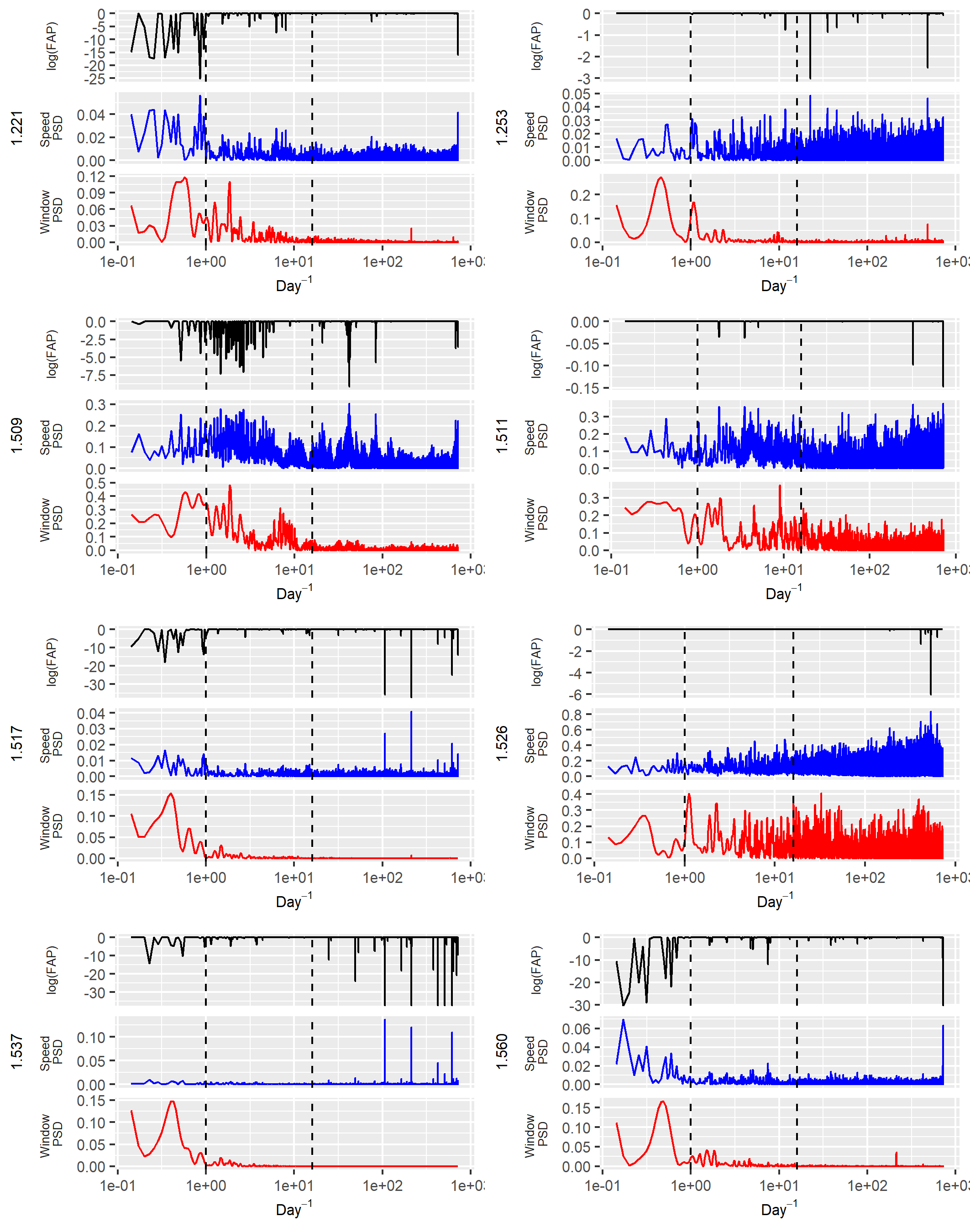

**Fig. S10. Lomb-Scargle Periodograms for High Density Males** Lomb_Scargle Periodograms of speed were calculated from Bayesian trilaterion predictions over the week of 10-10-2014. Dashed lines indicate periods of 24 hours and 90 min.

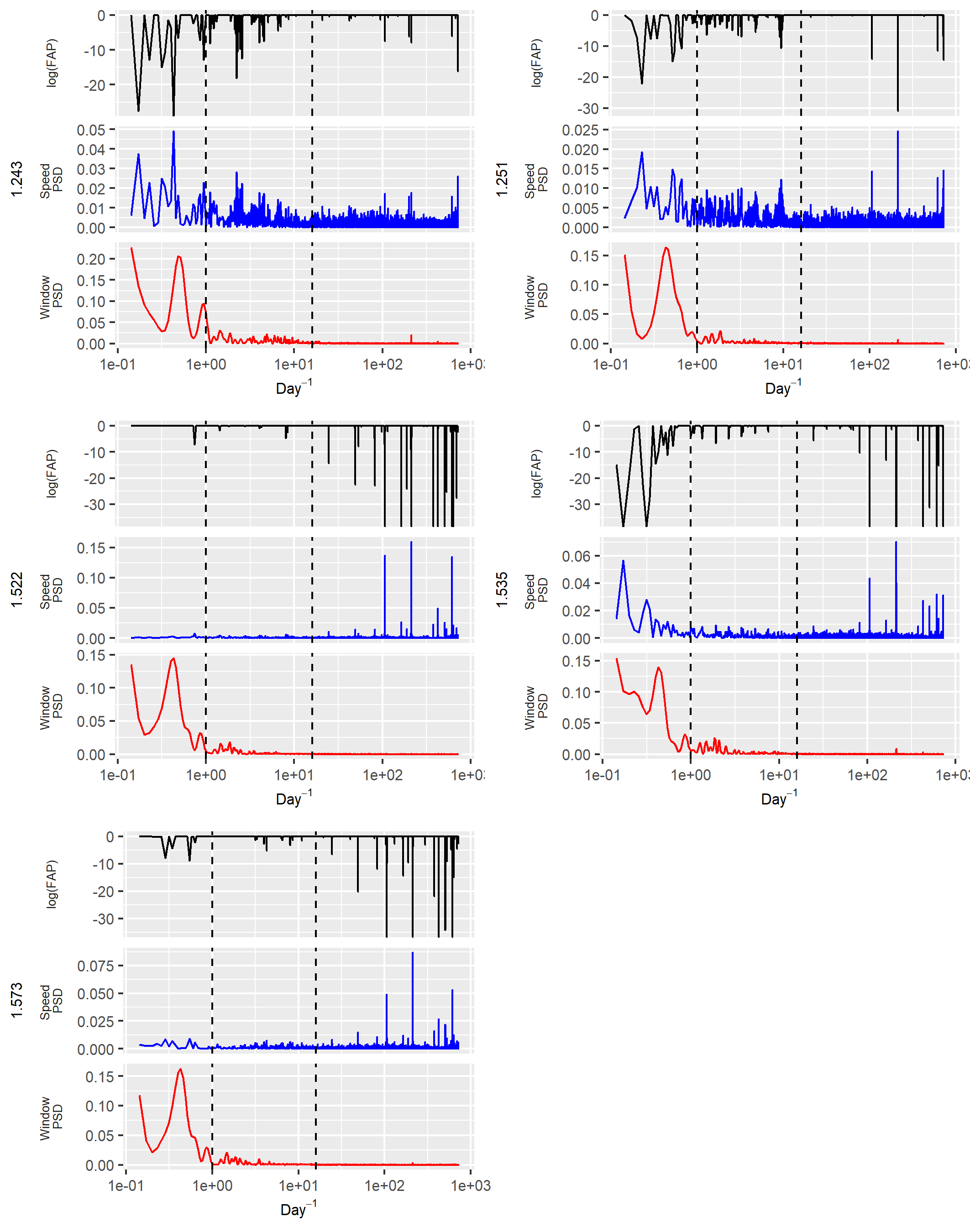

**Fig. S11. Lomb-Scargle Periodograms for High Density Females** Lomb_Scargle Periodograms of speed were calculated from Bayesian trilaterion predictions over the week of 10-10-2014. Dashed lines indicate periods of 24 hours and 90 min.

| Antenna | $\beta_{0}$ | $\beta_{1}$ | $\tau^{2}$ | $\sigma$ |
| --- | --- | --- | --- | --- |
| 1 | -84.5 | -1.59 | 0.037 | 5.2 |
| 2 | -83.4 | -1.68 | 0.022 | 6.7 |
| 3 | -78.4 | -2.04 | 0.034 | 5.4 |
| 4 | -85.4 | -1.58 | 0.045 | 4.7 |
| 5 | -82.2 | -1.79 | 0.041 | 4.9 |
| 6 | -85.4 | -1.53 | 0.052 | 4.4 |
| 7 | -81.8 | -1.94 | 0.034 | 5.4 |
| 8 | -81.9 | -1.85 | 0.031 | 5.7 |
| 9 | -85.1 | -1.45 | 0.026 | 6.2 |
| 10 | -73.3 | -2.19 | 0.023 | 6.6 |
| 11 | -91.0 | -1.06 | 0.025 | 6.3 |
| 12 | -81.9 | -1.70 | 0.021 | 6.9 |
| 13 | -68.7 | -2.43 | 0.036 | 5.3 |
| 14 | -107.4 | -0.12 | 0.024 | 6.5 |
| 15 | -91.5 | -1.07 | 0.016 | 7.9 |
| 16 | -79.3 | -1.69 | 0.025 | 6.3 |
| 17 | -80.7 | -1.67 | 0.025 | 6.3 |
| 18 | -84.9 | -1.51 | 0.022 | 6.7 |
| 19 | -102.6 | -0.42 | 0.025 | 6.4 |
| 20 | -91.8 | -0.89 | 0.027 | 6.0 |
| 21 | -95.7 | -0.68 | 0.022 | 6.7 |
| 22 | -107.7 | 0.08 | 0.016 | 7.9 |
| 23 | -80.0 | -1.81 | 0.016 | 8.0 |
| 24 | -88.9 | -1.15 | 0.012 | 9.0 |
| 25 | -77.7 | -2.01 | 0.027 | 6.1 |
| 26 | -89.1 | -1.39 | 0.015 | 8.3 |
| 27 | -84.0 | -1.70 | 0.022 | 6.7 |
| 28 | -89.5 | -1.30 | 0.026 | 6.2 |
| 29 | -91.4 | -1.14 | 0.017 | 7.7 |
| 30 | -86.4 | -1.50 | 0.036 | 5.3 |
| 31 | -84.7 | -1.54 | 0.023 | 6.6 |
| 32 | -84.3 | -1.59 | 0.039 | 5.1 |

**Table S1. Antenna Model Parameters.** Antenna posterior distribution parameter means for intercept, slope, and precision parameters. Precision is also reparameterized as standard deviation for clarity.
