## Supplementary material for "An Automated Radio-Telemetry System (ARTS) for Monitoring Small Mammals": Analysis R Code: 2014_KDE_Fitting.R

```
#load packages
library(here)
library(adehabitatHR)
library(sp)
library(tidyverse)
library(dtplyr)
library(data.table)
library(lubridate)

func_dir <- here::here("src", "ARTSFunctions.R")
source(func_dir)

#load 2014 Manual Tracking Data
data.dir <- here::here("data", "2014 Animal Data", "Manual Telemetry 2014.csv")
data_2014 <- read_csv(data.dir, 
                      col_names = TRUE,
                      col_types = cols(time = col_character()))

#load id table
id.dir <- here::here("data", "2014 Animal Data/identification 2014.csv")
id_2014 <- read_csv(id.dir, col_names = TRUE) %>%
  dplyr::select(id, retag)

#In manual telemetry data ID corresponds to tag at time of release not JCMS
#Mouse ID. 
id_2014 <- mutate(id_2014, manual_id = ifelse(is.na(retag), id, retag))

#Join tracking data to ids
data_2014 <- inner_join(data_2014, id_2014, by = c("ID" = "manual_id")) %>%

#Convert from feet to meters convert date and time to POSIXct
  mutate(time = mdy_hms(paste(date, time), tz = "America/Chicago"),
         x = x * 0.3048, 
         y = y * 0.3048) %>%
  dplyr::select(-ID, -date) 

#Male UDs, unnormalized
M_UD <- KDE2pass(filter(data_2014, Sex == "M"))

#female UDs, unnormalized
F_UD <- KDE2pass(filter(data_2014, Sex == "F"))

#Save UDs for later analysis
S4_manual_KDEs_2014 <- unlist(list(M_UD, F_UD), recursive = FALSE)
save(S4_manual_KDEs_2014, file = here::here("results", 
                                            "S4_manual_KDEs_2014.RData"))

#Save KDE_data
write_csv(data_2014, here::here("results",
                                "Manual_Telemetry_2014_tidy.csv"))
```
