## Supplementary material for "An Automated Radio-Telemetry System (ARTS) for Monitoring Small Mammals": Analysis R Code: analysis_makefile.R

```
library(here)

source(here::here("src", "2014_KDE_Fitting.R"))
rm(list = ls())
source(here::here("src", "Filtering.R"))
rm(list = ls())
source(here::here("src", "Automated_vs_Manual.R"))
rm(list = ls())
source(here::here("src", "split_training_testing.R"))
rm(list = ls())
source(here::here("src", "JAGS_Regression.R"))
rm(list = ls())
source(here::here("src", "ARTS_training.R"))
rm(list = ls())
source(here::here("src", "ARTS_testing.R"))
rm(list = ls())
source(here::here("src", "size_sensitivity.R"))
rm(list = ls())
source(here::here("src", "outlier_sensitivity.R"))
rm(list = ls())
source(here::here("src", "ARTS_prediction_2014.R"))
rm(list = ls())
source(here::here("src", "Lomb-Scargle_Analysis.R"))
rm(list = ls())
```
