## Supplementary material for "An Automated Radio-Telemetry System (ARTS) for Monitoring Small Mammals": Analysis R Code: ARTS_prediction_2014.R

```
#Fit locations for 2014 season
library(here)
source(here::here("src", "ARTSFunctions.R"))
library(data.table)
library(dtplyr)
library(tidyverse)
library(lubridate)
library(ggridges)
library(maptools)


load(here::here("results", "TrackingAug2_bayes_tables.RData"))

#Load Filtered Automated Data 
ARTS_2014 <- read_csv(here::here("results", "Tracking_2014.csv")) %>%
  mutate(Time = with_tz(Time, tz = "America/Chicago"),
         Time = round_date(Time, unit = "min"),
         Ant = AntEdit(.))

predictions <- ARTSPrediction(ARTS_2014, TrackingAug2_bayes_tables)

#labeling fitting method and training data
predictions <- mutate(predictions, 
                      training = "TrackingAug2",
                      method = "bayes",
                      ant_selection = "top3") %>%
  #Assign animal numbers to data
  inner_join(distinct(dplyr::select(ARTS_2014, ID, Freq, Time)))

#Load Manual KDEs, S4 objects, saved as .RData 
KDE.dir <- here::here("results", "S4_manual_KDEs_2014.RData")
load(KDE.dir)

#For each Manual fix find distance to KDE global maxima
KDE_max <- rbindlist(lapply(S4_manual_KDEs_2014, function(x){
  filter(as.data.frame(x@coords), x@data$ud == max(x@data$ud))
}))

#Label KDE_max
KDE_max <- mutate(KDE_max, ID = names(S4_manual_KDEs_2014)) %>%
  rename(KDE_max_x = Var2, KDE_max_y = Var1)

#join predictions and KDE_max
predictions <- inner_join(predictions, KDE_max)

#Compute distance to nest and speed
predictions  <- predictions %>% 
  group_by(ID) %>%
  mutate(nest_dist = sqrt((x - KDE_max_x)^2 + (y - KDE_max_y)^2),
         step_size = sqrt((x - lag(x))^2 + (y - lag(y))^2),
         step_time = as.numeric(Time - lag(Time)),
         speed = step_size / step_time) 

#Assign sex, and density to fixes
treatment_df <- read_csv(here::here("data", 
                                   "2014 Animal Data",
                                   "Release 2014.csv")) %>%
  dplyr::select(id, density) %>%
  rename(ID = id, Density = density)

colony_df <- read_csv(here::here("data", 
                                 "2014 Animal Data",
                                 "Colony Data 2014.csv")) %>%
  rename(ID = `Mouse ID`) %>%
  dplyr::select(ID, Sex)

predictions <- predictions %>% 
  inner_join(colony_df) %>%
  inner_join(treatment_df)

#crepuscule/sunriset take coordinates in long, lat NOT lat, long
coord <- matrix(c(-89.4948, 37.8884), ncol = 2)
angle <- 18

predictions <- mutate(predictions,
               dawn = crepuscule(coord,
                                 Time, 
                                 solarDep = angle, 
                                 direction="dawn", 
                                 POSIXct.out = TRUE)$time,
               sunrise = sunriset(coord,
                                  Time, 
                                  direction="sunrise", 
                                  POSIXct.out = TRUE)$time,
               sunset = sunriset(coord,
                                 Time, 
                                 direction="sunset", 
                                 POSIXct.out = TRUE)$time,
               dusk = crepuscule(coord,
                                 Time, 
                                 solarDep = angle, 
                                 direction="dusk", 
                                 POSIXct.out = TRUE)$time,
               period = case_when(Time < dawn | Time > dusk ~ "Night",
                                  Time > sunrise & Time < sunset ~ "Day",
                                  Time > dawn & Time < sunrise ~ "Dawn",
                                  Time > sunset & Time < dusk ~ "Dusk"))

write_csv(predictions,
          here::here("results", "ARTS_predictions_2014.csv"))
          


  


  
```
