## Supplementary material for "An Automated Radio-Telemetry System (ARTS) for Monitoring Small Mammals": Analysis R Code: ARTS_testing.R

```
#Script Begins
library(here)
library(tidyverse)
library(data.table)
library(dtplyr)
library(lubridate)
library(ggplot2)
library(GGally)

func_dir <- here::here("src", "ARTSFunctions.R")
source(func_dir)

#Intermediate Results directory (contains filtered training data, bayesian regression, etc)
results_dir <- here::here("results")

# Load Training Data and Bayesain parameter coeffs
training_data <- c("raw", "shift", "TrackingAug", "TrackingAug2")
testing_ls <- list()
BHR_model_coeffs <- list()
for(i in training_data) {
  #Load training data
  testing_ls[[i]] <-
    switch(i,
           raw = here::here("results", "Manual_ARTS_2014.csv"),
           shift = here::here("results", "Manual_ARTS_2014.csv"),
           TrackingAug = here::here("results", "Manual_ARTS_2014_TrackingAug.csv"),
           TrackingAug2 = here::here("results", "Testing_TrackingAug2.csv")
    ) %>%
    read_csv(.)%>%
  
    #bin times note that this decrease temporal resolutsion some locations in mapping
    #data are assigned to the same time
    mutate(Time = with_tz(Time, tzone = "America/Chicago"),
           Time = round_date(Time, unit = "min"),
           Ant = AntEdit(.)) %>%
    rename(trans_x = x, trans_y = y)
    
    #Exclude Manual fixes greater than 5 min from an ARTS fix
    if(i == "raw" | i == "shift") {
      testing_ls[[i]] <- testing_ls[[i]] %>%
                             filter(time_diff <= 5)
    }
  
  #Load bayesian regression coeffs
  load(paste0(results_dir, "/", i, "_BHR_model_coeffs.RData"))
  BHR_model_coeffs[[i]] <- get(paste0(i, "_BHR_model_coeffs"))
  rm(list = paste0(i, "_BHR_model_coeffs"))
}

#Load Cached model tables
method <- c("bayes", "NP")
bayes_tables <- list()
NP_tables <- list()
for (i in training_data){
  for (j in method){
    #Check for cached tables
    table_dir <- paste0(results_dir, "/", i, "_", j,"_tables.RData")
    if(file.exists(table_dir)){
      load(table_dir)
      if(j == "bayes"){
        bayes_tables[[i]] <- get(paste(i, j, "tables", sep ="_"))
      }
      if(j == "NP"){
        NP_tables[[i]] <- get(paste(i, j, "tables", sep ="_"))
      }
    } else {
      print("table not cached")
    }
    rm(list =  paste(i, j, "tables", sep ="_"))
  }
}

#Fit model predictions/residuals
method <- c("tri", "bayes", "NP", "random", "strongest")
ant_selection <- c("top3", "all")
model_predictions <- vector(mode = "list", length(method)*length(training_data)*length(ant_selection))
index <- 1
for(i in training_data) {
  for(j in method) {
    for(k in ant_selection) {
      #Splitting by Freq and Time
      fixes <- split(testing_ls[[i]], 
                     testing_ls[[i]]$Freq)
      fixes <- lapply(fixes, function(x) x[order(x$Time),])
      fixes <- lapply(fixes, function(x) {
        split(x, as.factor(as.character(x$Time)))
      })
      
      #Select only fixes with 3 unique antenna and 1 transmitter location
      fixes <- lapply(fixes, function(x){
        x[which(sapply(x, function(x) {
          length(unique(x$Ant)) >= 3 & 
          length(unique(interaction(x$trans_x, x$trans_y))) == 1
        }))]
      })
      
      if(k == "top3"){
        fixes <- lapply(fixes, function(x) {
          lapply(x, Top3)
        })
      }
      
      
      #NLS Fitting
      if(j == "tri") {
        prediction <- lapply(fixes, function(x) {
          lapply(x, function(x, model) {
            try(Trilaterate(x, model))}, 
            model = BHR_model_coeffs[[i]])
        })
        
        #Format fitting varialbe names
        prediction <- FormatNLS(prediction)
        prediction <- mutate(prediction,
                             entropy = as.numeric(NA),
                             training = i,
                             method = j,
                             ant_selection = k) 
      }
      
      #bayeian fitting
      if(j == "bayes"){
        prediction <- lapply(fixes, function(x) {
          lapply(x, PositionPDF, tables = bayes_tables[[i]])
        })
        
        prediction <- lapply(prediction, function(x) {
          lapply(x, MAP.est)
        })
        
        #Format fitting varialbe names
        prediction <- FormatMAP(prediction)
        prediction <- mutate(prediction, 
                             training = i,
                             method = j,
                             ant_selection = k)
      }
      
      #NP fitting 
      if(j == "NP") {
        prediction <- lapply(fixes, function(x) {
          lapply(x, PositionPDF, tables = NP_tables[[i]])
        })
        
        prediction <- lapply(prediction, function(x) {
          lapply(x, MAP.est)
        })
        
        prediction <- FormatMAP(prediction)
        prediction <- mutate(prediction, 
                             training = i,
                             method = j,
                             ant_selection = k)
      }
      
      #Random fitting
      if(j == "random") {
        prediction <- lapply(fixes, bind_rows) %>%
          bind_rows() %>%
          group_by(Time, Freq) %>%
          summarize(x = runif(1, 0, 180 * 0.3048),
                    y = runif(1, 0, 240 * 0.3048)) %>%
          mutate(entropy = as.numeric(NA),
                 training = i,
                 method = "random",
                 ant_selection = k
                 )
      }
      
      if(j == "strongest") {
        #Presence absences at strongest antenna
        prediction <- lapply(fixes, bind_rows) %>%
          bind_rows() %>%
          group_by(Time, Freq) %>%
          filter(Power == max(Power)) %>%
          #Assign ties to geometic mean of ant locations
          summarize(x = exp(1/n() * sum(log(ant_x))), 
                    y = exp(1/n() * sum(log(ant_y))),
                    entropy = as.numeric(NA),
                    training = i,
                    method = "strongest",
                    ant_selection = k)
      }
      
      prediction <- prediction %>%
        rename(pred_x = x, pred_y = y)%>%
        tbl_df() %>%
        inner_join(testing_ls[[i]]) %>%
        dplyr::select(pred_x,
                      pred_y,
                      trans_x, 
                      trans_y,
                      entropy,
                      Time,
                      Freq,
                      training,
                      method,
                      ant_selection)%>%
        distinct()
      
      model_predictions[[index]] <- prediction
      index <- index + 1
      rm(prediction)
    }
  }
}

model_predictions <- bind_rows(model_predictions) %>%
  mutate(error = sqrt((trans_x - pred_x)^2 + (trans_y - pred_y)^2))

write_csv(model_predictions,
          path = here::here("results", "testing_error.csv"))

model_predictions %>%
  group_by(training, method, ant_selection) %>%
  summarize(mode_error = CalcMode(error),
            median_error = median(error, na.rm = T),
            mean_error = mean(error, na.rm=T),
            time_points = nrow(.),
            fix_num = sum(!is.na(pred_x)),
            fix_drop = sum(is.na(pred_x))) %>%
  print.data.frame()

if(!dir.exists(here::here("docs", "ARTS_Testing"))) {
  dir.create(here::here("docs", "ARTS_Testing"))
}

#ECDF of errors
model_predictions %>%
  filter(!(method %in% c("random", "strongest") & ant_selection == "top3")) %>%
  mutate(method = interaction(method, ant_selection), sep = "_") %>%
ggplot(aes(error, color = method)) +
  stat_ecdf(size = 1) +
  facet_grid(training~.) +
  labs(x = "Error [m]",
       y = "Quantile",
       title = "Density 2014: Emperical CDF of Error") +
  theme(text = element_text(size=16),
        axis.title=element_text(face="bold"))
ggsave(here::here("docs",
                  "ARTS_Testing",
                  "error_ecdf.pdf"), 
       device = "pdf")

#Histograms of Error
model_predictions %>%
filter(!(method %in% c("random", "strongest") & ant_selection == "top3")) %>%
  mutate(method = interaction(method, ant_selection, sep = "_")) %>%
ggplot(aes(error, fill = method)) +
  geom_histogram(aes(y = ..density..)) +
  facet_grid(training~method) +
  labs(x = "Error [m]",
       title = "Density 2014: Error") +
  theme(text = element_text(size=16),
        axis.title=element_text(face="bold"))
ggsave(here::here("docs",
                  "ARTS_Testing",
                  "error_hist.pdf"), 
       device = "pdf")

#error correlation by method 
for(i in training_data){
  model_predictions %>%
    filter(!(method %in% c("random", "strongest") & ant_selection == "top3"),
           training == i) %>%
    mutate(method = interaction(method, ant_selection, sep = "_")) %>%
    dplyr::select(Time, Freq, method, error) %>%
    spread(method, error) %>%
    dplyr::select(-Time, -Freq) %>%
    ggpairs( lower = list(continuous = wrap("points", shape = 1)),
             title = paste("Residual correlation", i, "training data"))
  ggsave(here::here("docs",
                    "ARTS_Testing",
                    paste0(i, "_error_corr.pdf")), 
         device = "pdf") 
}

model_predictions %>%
  filter(!(method %in% c("random", "strongest") & ant_selection == "top3")) %>%
  mutate(method = interaction(method, ant_selection, sep = "_")) %>%
  ggplot(aes(x = pred_x, y = pred_y, color = error)) +
    geom_point() +
    facet_grid(training ~ method) +
    scale_color_gradient(low = "green", high = "red")+
    labs(title = "Error vs predicted position") +
    theme(text = element_text(size=16),
          axis.title=element_text(face="bold"))
ggsave(here::here("docs",
                  "ARTS_Testing",
                  "error_vs_predxy.jpeg"), 
       device = "pdf") 

model_predictions %>%
  filter(!(method %in% c("random", "strongest") & ant_selection == "top3")) %>%
  mutate(method = interaction(method, ant_selection, sep = "_")) %>%
  ggplot(aes(x = trans_x, y = trans_y, color = error)) +
    geom_point() +
    scale_color_gradient(low = "green", high = "red") +
    facet_grid(training ~ method)+
  labs(title = "Error vs known position")+
    theme(text = element_text(size=16),
          axis.title=element_text(face="bold"))
ggsave(here::here("docs",
                  "ARTS_Testing",
                  "error_vs_knownxy.jpeg"), 
       device = "pdf") 

   
# #Error at Nest
# model_predictions %>%
#   mutate(method = interaction(method, ant_selection, sep = "_")) %>%
#   filter(method == "bayes_top3") %>%
#   ggplot(aes(x = ifelse(nest_dist_KDE < 2, "At Nest", "Away"), y = error)) +
#   geom_boxplot() +
#   labs(x = "Manual Fix Location",
#        y = "Error [m]",
#        title = "Do Burrows Atler Signal Propagation?") +
#   theme(text = element_text(size=18),
#         axis.title=element_text(face="bold")) +
#   ggsave("Nest_Away_Boxplot.pdf", device = "pdf")
#
# #Error vs Distance to Nest
# filter(plot_data, key == "bayes_top3") %>%
#   ggplot(aes(x = nest_dist_KDE, y = error)) +
#   geom_point() +
#   geom_smooth(method = "lm") +
#   labs(x = "Distance to Nest from Manual Fix [m]",
#        y = "Error [m]",
#        title = "Error vs Distance to Nest") +
#   theme(text = element_text(size=18),
#         axis.title=element_text(face="bold")) +
#   ggsave("error_vs_nestdist.pdf", device = "pdf")
#
# #Entropy by fix method
# ggplot(plot_data, aes(x = key, y = entropy, fill = key)) +
# geom_boxplot() +
# labs(x = "Fix Method",
#      y = "Entropy",
#      title = "Fix-Method Dispersion") +
# theme(text = element_text(size=18),
#       axis.title=element_text(face="bold")) +
# ggsave("Entropy_Boxplot.pdf", device = "pdf")
#
# #Error vs Distance to Nest
# filter(plot_data, key == "bayes_top3") %>%
#   ggplot(aes(x = error, y = entropy)) +
#   geom_point() +
#   geom_smooth(method = "lm") +
#   labs(x = "Error [m]",
#        y = "Entropy",
#        title = "Error vs Entropy") +
#   theme(text = element_text(size=18),
#         axis.title=element_text(face="bold")) +
#   ggsave("entropy_vs_error.pdf", device = "pdf")
#
# #Nest Distance vs Entropy
# filter(plot_data, key == "bayes_top3") %>%
#   ggplot(aes(x = nest_dist_KDE, y = entropy)) +
#   geom_point() +
#   geom_smooth(method = "lm") +
#   labs(x = "Distance to Nest from Manual Fix [m]",
#        y = "Entropy",
#        title = "Distance to Nest vs Entropy") +
#   theme(text = element_text(size=18),
#         axis.title=element_text(face="bold")) +
#   ggsave("Nest_Away_ent_Boxplot.pdf", device = "pdf")
```
