## Supplementary material for "An Automated Radio-Telemetry System (ARTS) for Monitoring Small Mammals": Analysis R Code: ARTS_training.R

```
#Script Begins
library(here)
library(tidyverse)
library(data.table)
library(dtplyr)
library(lubridate)
library(GGally)

func_dir <- here::here("src", "ARTSFunctions.R")
source(func_dir)

#Intermediate Results directory (contains filtered training data, bayesian regression, etc)
results_dir <- here::here("results")

# Load Training Data and Bayesain parameter coeffs
training_data <- c("raw", "shift", "TrackingAug", "TrackingAug2")
training_ls <- list()
BHR_model_coeffs <- list()
for(i in training_data) {
  #Load training data
  training_ls[[i]] <-
    switch(i,
      raw = here::here("results", "Mapping_2014_raw.csv"),
      shift = here::here("results", "Mapping_2014_shift.csv"),
      TrackingAug = here::here("results", "Mapping_2014_TrackingAug.csv"),
      TrackingAug2 = here::here("results", "Training_TrackingAug2.csv")
      ) %>%
      read_csv(.) %>%
      mutate(Time = with_tz(Time, tzone = "America/Chicago"),
             Time = round_date(Time, unit = "min"),
             Ant = AntEdit(.)) 

  #Load bayesian regression coeffs
  load(paste0(results_dir, "/", i, "_BHR_model_coeffs.RData"))
  BHR_model_coeffs[[i]] <- get(paste0(i, "_BHR_model_coeffs"))
  rm(list = paste0(i, "_BHR_model_coeffs"))
}

#Load Antenna Coordinates
antenna_dir <- here::here("data", "All Radiomapping Keys", "Antenna Key.csv")
ant_coor<- read_csv(file = antenna_dir) %>%
  
#Label antenna 1-32, convert feet to meters
  mutate(Ant = AntEdit(.),
         X = X * 0.3048,
         Y = Y * 0.3048)

#Calculate and Cache model tables
method <- c("bayes", "NP")
bayes_tables <- list()
NP_tables <- list()
for (i in training_data){
  for (j in method){
    #Check for cached tables
    table_dir <- paste0(results_dir, "/", i, "_", j,"_tables.RData")
      if(file.exists(table_dir)){
        load(table_dir)
        if(j == "bayes"){
          bayes_tables[[i]] <- get(paste(i, j, "tables", sep ="_"))
        }
        if(j == "NP"){
          NP_tables[[i]] <- get(paste(i, j, "tables", sep ="_"))
        }
      } else {
        #Calculate tables
        if (j == "bayes") {
          table <- BHR_model_coeffs[[i]] %>%
          CalcTablesBayes(ant_coor)
          bayes_tables[[i]] <- table
        }
        if(j == "NP"){
          #Assing training data
          table <- training_ls[[i]] %>%
            mutate(Ant = AntEdit(.)) %>%
            CalcTablesNP()
            NP_tables[[i]] <- table
        }
        
        #Assign variable name and save
        assign(paste(i, j, "tables", sep ="_"), table)
        save(list= paste(i, j, "tables", sep ="_"),
             file = paste0(results_dir, "/", i, "_", j, "_tables", ".RData"))
        rm(table)
      }
    rm(list =  paste(i, j, "tables", sep ="_"))
  }
}

#Fit model predictions/residuals
method <- c("tri", "bayes", "NP")
ant_selection <- c("top3", "all")
model_predictions <- vector(mode = "list", length(method)*length(training_data)*length(ant_selection))
index <- 1
for(i in training_data) {
  for(j in method) {
    for(k in ant_selection) {
      #Splitting by Freq and Time
      fixes <- split(training_ls[[i]], 
                        training_ls[[i]]$Freq)
      fixes <- lapply(fixes, function(x) x[order(x$Time),])
      fixes <- lapply(fixes, function(x) {
        split(x, as.factor(as.character(x$Time)))
      })
      
      #Select only fixes with 3 unique antenna and 1 transmitter location
      fixes <- lapply(fixes, function(x){
        x[which(sapply(x, function(x) {
          length(unique(x$Ant)) >= 3 & 
          length(unique(interaction(x$trans_x, x$trans_y))) == 1
          }))]
      })
      
      if(k == "top3"){
        fixes <- lapply(fixes, function(x) {
          lapply(x, Top3)
        })
      }
      
      
      #NLS Fitting
      if(j == "tri") {
        prediction <- lapply(fixes, function(x) {
          lapply(x, function(x, model) {
            try(Trilaterate(x, model))}, 
            model = BHR_model_coeffs[[i]])
        })
        
        #Format fitting varialbe names
        prediction <- FormatNLS(prediction)
        prediction <- mutate(prediction,
                      entropy = as.numeric(NA),
                      training = i,
                      method = j,
                      ant_selection = k) 
      }
      
      #bayeian fitting
      if(j == "bayes"){
        prediction <- lapply(fixes, function(x) {
          lapply(x, PositionPDF, tables = bayes_tables[[i]])
        })
        
        prediction <- lapply(prediction, function(x) {
          lapply(x, MAP.est)
        })
        
        #Format fitting varialbe names
        prediction <- FormatMAP(prediction)
        prediction <- mutate(prediction, 
                             training = i,
                             method = j,
                             ant_selection = k)
      }
      
      #NP fitting raw
      if(j == "NP") {
        prediction <- lapply(fixes, function(x) {
          lapply(x, PositionPDF, tables = NP_tables[[i]])
        })
        
        prediction <- lapply(prediction, function(x) {
          lapply(x, MAP.est)
        })
        
        prediction <- FormatMAP(prediction)
        prediction <- mutate(prediction, 
                             training = i,
                             method = j,
                             ant_selection = k)
      }
      
      prediction <- prediction %>%
        rename(pred_x = x, pred_y = y)%>%
          tbl_df() %>%
          inner_join(training_ls[[i]]) %>%
          dplyr::select(pred_x,
                        pred_y,
                        trans_x, 
                        trans_y,
                        entropy,
                        Time,
                        Freq,
                        training,
                        method,
                        ant_selection)%>%
          distinct()
      
      model_predictions[[index]] <- prediction
      index <- index + 1
      rm(prediction)
    }
  }
}

model_predictions <- bind_rows(model_predictions) %>%
  mutate(residual= sqrt((trans_x - pred_x)^2 + (trans_y - pred_y)^2))

write_csv(model_predictions,
          path = here::here("results", "training_residuals.csv"))

CalcMode <- function(x){
  d <- density(x, na.rm = T)
  d$x[which(d$y == max(d$y))]
}

model_predictions %>%
  group_by(training, method, ant_selection) %>%
  summarize(mode_residual = CalcMode(residual),
            median_residual = median(residual, na.rm = T),
            mean_residual = mean(residual, na.rm=T),
            time_points = nrow(.),
            fix_num = sum(!is.na(pred_x)),
            fix_drop = sum(is.na(pred_x))) %>%
  print.data.frame()

if(!dir.exists(here::here("docs", "ARTS_Training"))) {
  dir.create(here::here("docs", "ARTS_Training"))
}
#ECDF of residuals
ggplot(model_predictions, aes(residual, color = interaction(method, ant_selection, sep = "_"))) +
  stat_ecdf(size = 1) +
  facet_grid(training~.) +
  labs(x = "Residual [m]",
       y = "Quantile",
       title = "Mapping 2014: Emperical CDF of Residuals",
       color = "method") +
  theme(text = element_text(size=16),
        axis.title=element_text(face="bold"))
ggsave(here::here("docs",
                  "ARTS_Training",
                  "residuals_ecdf.pdf"), 
       device = "pdf")

#Histograms of Residuals
ggplot(model_predictions, aes(residual, fill = interaction(method, ant_selection, sep = "_"))) +
  geom_histogram(aes(y = ..density..)) +
  facet_grid(training~interaction(method, ant_selection, sep = "_")) +
  labs(x = "Residual [m]",
       title = "Mapping 2014: Residuals",
       fill = "method") +
  theme(text = element_text(size=16),
        axis.title=element_text(face="bold"))
ggsave(here::here("docs",
                  "ARTS_Training",
                  "residual_hist.pdf"), 
       device = "pdf")

#residual correlation by method for raw data
for(i in training_data){
  model_predictions %>%
    filter(training == i) %>%
    mutate(method = paste(method, ant_selection, sep ="_")) %>%
    dplyr::select(Time, Freq, method, residual) %>%
    spread(method, residual) %>%
    dplyr::select(-Time, -Freq) %>%
    ggpairs( lower = list(continuous = wrap("points", shape = 1)),
             title = paste("Residual correlation", i, "training data"))
  ggsave(here::here("docs",
                    "ARTS_Training",
                    paste0(i, "_residual_corr.pdf")), 
         device = "pdf") 
}


ggplot(model_predictions, aes(x = pred_x, y = pred_y, color = residual)) +
  geom_point() +
  facet_grid(training ~  interaction(method, ant_selection, sep = "_")) +
  scale_color_gradient(low = "green", high = "red")+
  labs(title = "Residual vs predicted position") +
  theme(text = element_text(size=16),
        axis.title=element_text(face="bold"))
ggsave(here::here("docs",
                  "ARTS_Training",
                  "residuals_vs_predxy.jpg"),
       height = 7,
       width = 7,
       device = "jpg")

ggplot(model_predictions, aes(x = trans_x, y = trans_y, color = residual)) +
  geom_point() +
  scale_color_gradient(low = "green", high = "red") +
  facet_grid(training ~  interaction(method, ant_selection, sep = "_"))+ 
  labs(title = "Residual vs known position")+
  theme(text = element_text(size=16),
        axis.title=element_text(face="bold"))
ggsave(here::here("docs",
                  "ARTS_Training",
                  "residuals_vs_knownxy.jpg"),
       height = 7,
       width = 7,
       device = "jpg")
```
