## Supplementary material for "An Automated Radio-Telemetry System (ARTS) for Monitoring Small Mammals": Analysis R Code: ARTS_vs_Manual.R

```
#Load required packages 
library(here)
library(tidyverse)
library(dtplyr)
library(data.table)
library(lubridate)
library(parallel)
library(adehabitatHR)
library(sp)

#Source functions
func_dir <- here::here("src", "ARTSFunctions.R")
source(func_dir)

#Load Filtered Automated Data 
ARTS_2014.dir <- here::here("results", "Tracking_2014.csv")
ARTS_2014 <- read_csv(ARTS_2014.dir) %>%
  mutate(Time = with_tz(Time, tz = "America/Chicago"))

#Load Manual Fixes make sure to load the tidy data set with dates and
# times in the Y-m-d H:M:S format and distances in meters not feet
Manual_2014.dir <- here::here("results",
                              "Manual_Telemetry_2014_tidy.csv")
Manual_2014 <- read_csv(Manual_2014.dir) %>%
  mutate(time = with_tz(time, tz = "America/Chicago"))


#Load Manual KDEs, S4 objects, saved as .RData 
KDE.dir <- here::here("results", "S4_manual_KDEs_2014.RData")
load(KDE.dir)

#Convert Automated and Manual data to data tables to improve speed
ARTS_2014 <- data.table(ARTS_2014)
Manual_2014 <- data.table(Manual_2014)

#Bin Automated data to nearest minute
ARTS_2014 <- mutate(ARTS_2014, Time = round_date(Time, unit = "min")) %>%
  
  #Count distinct contacts per minute
  group_by(ID, Freq, Time) %>%
  mutate(contacts = n_distinct(Site, Ant)) %>%
  
  #Remove contacts < 2
  filter(contacts > 2) %>%
  ungroup()

#set up parallel cluster
cl <- makeCluster(detectCores() - 1)

#Calculate Nearest times
tdiffs_2014 <- ManualvsARTS_Time(ARTS_2014, Manual_2014, cluster = cl)

stopCluster(cl)

#Calculate Geometric means
tdiffs_2014 <- tdiffs_2014 %>%
  group_by(id) %>%
  mutate(geom_x = exp(mean(log(x))),
         geom_y = exp(mean(log(y))),
         nest_dist_geom = sqrt((x - geom_x)^2 + (y - geom_y)^2))

#For each Manual fix find distance to KDE global maxima
KDE_max <- rbindlist(lapply(S4_manual_KDEs_2014, function(x){
  filter(as.data.frame(x@coords), x@data$ud == max(x@data$ud))
}))

#Label KDE_max
KDE_max <- mutate(KDE_max, id = names(S4_manual_KDEs_2014)) %>%
            rename(KDE_max_x = Var2, KDE_max_y = Var1)

#join tdiffs and KDE_max
Manual_ARTS_2014 <- inner_join(tdiffs_2014, KDE_max) %>%
  mutate(nest_dist_KDE = sqrt((KDE_max_x - x)^2 + (KDE_max_y - y)^2),
         geom_KDE_dist = sqrt((KDE_max_x - geom_x)^2 + (KDE_max_y - geom_y)^2))

Man_ARTS_plot <- ggplot(Manual_ARTS_2014, aes())

# Man_ARTS_plot +
#   geom_histogram(aes(x = time_diff)) +
#   labs(title = "Manaul vs Automated Fixes") +
#   xlab("Time Difference [min]") + 
#   theme(plot.title = element_text(hjust = 0.5))
# 
# Man_ARTS_plot +
#   geom_histogram(aes(x = time_diff)) +
#   labs(title = "Manaul vs Automated Fixes") +
#   xlab("Time Difference [min]") + 
#   theme(plot.title = element_text(hjust = 0.5)) +
#   xlim(NA, 60) +
#   theme(axis.title=element_text(size=18,face="bold"))
# ggsave("Hist Time to Manual fix 1 hr.pdf",
#        device = "pdf",
#        width = 6,
#        height = 4,
#        units = "in"
#        )
# 
# Man_ARTS_plot +
#   geom_histogram(aes(x = time_diff)) +
#   labs(title = "Manaul vs Automated Fixes") +
#   xlab("Time Difference [min]") + 
#   theme(plot.title = element_text(hjust = 0.5)) +
#   xlim(NA, 10)
# 
# Man_ARTS_plot +
#   geom_histogram(aes(x = geom_KDE_dist)) +
#   labs(title = "Geometric Mean vs KDE Max") +
#   xlab("Difference [m]") + 
#   theme(plot.title = element_text(hjust = 0.5))
# 
# Man_ARTS_plot +
#   geom_histogram(aes(x = nest_dist_KDE)) +
#   labs(title = "Distance from KDE Max") +
#   xlab("Distance [m]") + 
#   theme(plot.title = element_text(hjust = 0.5))+ 
#   theme(plot.title = element_text(hjust = 0.5)) +
#   theme(axis.title=element_text(size=18,face="bold"))
# ggsave("Hist Dist to KDE Max.pdf",
#        device = "pdf",
#        width = 6,
#        height = 4,
#        units = "in"
# )
# 
# Man_ARTS_plot +
#   geom_histogram(aes(x = nest_dist_KDE)) +
#   labs(title = "Distance from KDE Max") +
#   xlab("Distance [m]") + 
#   theme(plot.title = element_text(hjust = 0.5)) +
#   xlim(NA, 10)
# 
# Man_ARTS_plot +
#   geom_point(aes(x = nest_dist_KDE, y = nest_dist_geom)) +
#   geom_smooth(aes(x = nest_dist_KDE, y = nest_dist_geom), method = "lm") +
#   labs(title = "Distance from Nest: Geometric Mean vs KDE Max") +
#   xlab("KDE Max Distance [m]") + 
#   ylab("Geometric Mean Distance [m]") +
#   theme(plot.title = element_text(hjust = 0.5)) +
#   theme(axis.title=element_text(size=18,face="bold"))
# ggsave("Geom vs KDE Max.pdf",
#        device = "pdf",
#        width = 6,
#        height = 4,
#        units = "in"
# )

#Join ARTS data to Manual_ARTS_2014 to get nearest contacts for each manual fix
#._join on data.tables has problems with more than 1 key in dtply 0.5.0. 
#Convert to tbl_df to join and convert back
Manual_ARTS_2014 <- data.table(inner_join(as.data.frame(Manual_ARTS_2014),
           as.data.frame(ARTS_2014), 
           by = c("id" = "ID", "Time" = "Time")))

#Path to antenna key
antennakey_path <-  here::here("data", "All Radiomapping Keys/Antenna Key.csv")
#Load antenna key
ant_key <- read_csv(antennakey_path) 

#Convert Ant key to meters
ant_key <- ant_key %>%
  mutate(ant_x = X *0.3048, ant_y = Y *0.3048) %>%
  dplyr::select(-X, -Y)

#join Manual_ARTS_2014 to ant_key to get antenna coordinates
Manual_ARTS_2014 <- data.table(inner_join(as.data.frame(Manual_ARTS_2014), 
                                         ant_key))

#Calculate distance between Nest Site and ant
Manual_ARTS_2014 <- mutate(Manual_ARTS_2014,
                          ant_nest_dist = sqrt((KDE_max_x-ant_x)^2 + 
                                                 (KDE_max_y - ant_y)^2)) %>% 
  arrange(id, Time) %>%
  rename(manual_time = time)

write_csv(Manual_ARTS_2014, 
          here::here("results", "Manual_ARTS_2014.csv"))

# #Summarize Max power and nest_ant distance for each contact
# Max_Power_Summary <- Manual_ARTS_2014 %>%
#   group_by(id, manual_time) %>%
#   filter(Power == max(Power))
# 
# 
# Man_ARTS_Power_plot <- ggplot(filter(Manual_ARTS_2014, time_diff <= 5), aes())
# 
# #Distribution of power at nest vs away
# Man_ARTS_Power_plot +
#   geom_histogram(aes(x = Power,
#                      y = ..density..,
#                      fill = ifelse(nest_dist_KDE < 2, "At Nest", "Away"))) +
#   facet_wrap(~ ifelse(nest_dist_KDE < 2, "At Nest", "Away")) +
#   labs(title = "Power at Nest vs Power Away") +
#   xlab("Power [Dbm]") +
#   theme(plot.title = element_text(hjust = 0.5)) +
#   scale_fill_discrete(guide = guide_legend(title = NULL)) +
#   theme(axis.title=element_text(size=18,face="bold"))
# ggsave("Power at Nest vs Power Away.pdf", 
#        device = "pdf", 
#        width = 6, 
#        height = 4, 
#        units = "in")
# 
# ggplot(filter(Max_Power_Summary, time_diff <= 5), aes(x = Power,
#                       y = ..density..,
#                       fill = ifelse(nest_dist_KDE < 2, "At Nest", "Away"))) +
#   geom_histogram() +
#   facet_wrap(~ ifelse(nest_dist_KDE < 2, "At Nest", "Away")) +
#   labs(title = "Max Power per contact at Nest vs Power Away") +
#   xlab("Max Power [Dbm]") +
#   theme(plot.title = element_text(hjust = 0.5)) +
#   scale_fill_discrete(guide = guide_legend(title = NULL))
# ggsave("Max Power per contact at Nest vs Power Away.pdf", 
#        device = "pdf", 
#        width = 11, 
#        height = 8.5, 
#        units = "in")
# 
# 
# ggplot(Manual_ARTS_2014, aes(x = ant_nest_dist, 
#                             y = Power)) +
#   geom_point(aes(color = ifelse(nest_dist_KDE < 2, "At Nest", "Away")),
#              alpha = 1/4) +
#   labs(title = "Power at Nest vs Nest-Antenna Distance", col = NULL) +
#   xlab("Distance between antenna and nest [m]") +
#   ylab("Power [Dbm]") +
#   theme(plot.title = element_text(hjust = 0.5)) +
#   theme(axis.title=element_text(size=18,face="bold"))
# ggsave("Power at Nest vs Nest-Antenna Distance.pdf",
#        device = "pdf",
#        height = 4,
#        width = 6,
#        units = "in")
# 
# ggplot(Max_Power_Summary, aes(x = ant_nest_dist, 
#                             y = Power)) +
#   geom_point(aes(color = ifelse(nest_dist_KDE < 2, "At Nest", "Away")),
#              alpha = 1/4) +
#   labs(title = "Max Power at Nest vs Nest-Antenna Distance",
#        y = "Power [Dbm]",
#        x = "Distance between antenna and nest [m]",
#        col = NULL)
# ggsave("Max Power at Nest vs Nest-Antenna Distance.pdf",
#        device = pdf,
#        height = 11,
#        width = 8.5,
#        units = "in")
  
  
  
```
