## Supplementary material for "An Automated Radio-Telemetry System (ARTS) for Monitoring Small Mammals": Analysis R Code: ARTSFunctions.R

```
#Rotation
#Take cartesian xy coordinates and rotates by theta units of pi.
Rotate <- function(x, y, theta){
  if (length(x) != length(y)){
    stop("x and y coordinates differ in length")
  } 
  data <- matrix(c(x, y), nrow = 2, byrow = TRUE)
  R <- matrix(c(cospi(theta), sinpi(theta), -sinpi(theta), cospi(theta)), ncol = 2)
  result <- R%*%data
  rownames(result) <- c("x", "y")
  data.frame(t(result))
}

#Helper function to Rotate coordinates within data frames.
##Args: 
##  df:A dataframe, tibble, data.table, etc
##   x: Character string specifying x coordinate columns
##   y: Character string specifying y coordinate columns
##theta: Units of pie to rotate
##Returns: A the original data frame with   
RotateDF <- function(df, x, y, theta){
  #Check that each column pairing is mapped to a rotation
  if (length(x) != length(y)) {
    stop("Number of x and y coordinates do not match")
  }
  
  
  #Rotations
  for(i in seq_along(x)) {
    trans <- Rotate(df[[x[i]]], 
                    df[[y[i]]],
                    theta = theta)
    df[, x[i]] <- trans$x
    df[, y[i]] <- trans$y
  }

  df
}

#Computes KDEs using mean href bandwidth for all animals of the same sex
#Args
#xy: A spatialpointsdataframe with id, x and y coordinates
#Returns
#A list of spatial
KDE2pass <- function(xy){
  require(dplyr)
  require(adehabitatHR)
  require(sp)
  
  #Format xy
  xy <- dplyr::select(xy, id, x, y) %>%
    group_by(id) %>%
    filter(n() >= 5) %>%
    ungroup()
  
  #Formatting data to spatial points dataframes
  #note: dplyr etc will not work on S4 objects
  coordinates(xy) <- c("x","y")
  
  #Constructing a 55 x 74 m grid with resolution 1 m^2  resolution
  x.lattice <- seq(0,55, by=1) 
  y.lattice <- seq(0,74, by=1) 
  xy.grid <- expand.grid(x=x.lattice, y=y.lattice) 
  coordinates(xy.grid) <- ~x+y
  gridded(xy.grid) <- TRUE 
  
  
  pass1 <- kernelUD(xy, h = "href", grid = xy.grid, extent = 0)
  pass1.h <- lapply(pass1, slot, "h")
  pass1.h.vector <- sapply(pass1.h, "[[", 1)
  h.ave <- mean(pass1.h.vector)
  pass2 <- kernelUD(xy, h = h.ave, grid = xy.grid, extent = 0)
}  

# Filters rows with duplicate Time, Site, Ant, and Freq by maximum 
# power. Optionally filters by interpulse interval (IPI) and interpulse-power 
# matching (PPM).
# 
# Args:
#   x: An POSIXct formated orion output.
#   IPI.min: Minimum interpulse interval (seconds).
#   IPI.max: Maximum interpulse interval (seconds).
#   PPM: Pulse-power matching (dBm).
#
# Returns: Dataset filter by duplicates, IPI and PPM
#
FilterNoise <- function(data, IPI.min, IPI.max, PPM){
  #load required packages
  require(dplyr)
  require(dtplyr)
  require(data.table)
  
  data <- data %>%
    
    #Convert to data.table
    tbl_dt() %>%
    
    #Remove duplicate contacts
    distinct() %>% 
    
    #Choose Contacts within Transmitter IPI if internal orion filtering on
    filter(Code == 0 | (Code > IPI.min & Code < IPI.max)) %>%
    
    #Choose strongest contact at given time
    group_by(Time, Site, Ant, Freq) %>% 
    filter(Power == max(Power)) %>% 
    
    #Sort by time ascending
    ungroup() %>%
    group_by(Site, Ant, Freq) %>% 
    arrange(Time) %>%
    ungroup()
  
  if (nargs() == 4) {
  
  #Split data into channels (unique combinations of Site, Antenna, Freq)  
  data <- with(data,
                split(data, 
                list(as.factor(Site), as.factor(Ant), as.factor(Freq)), 
                drop = TRUE))
  
  #Create Filtering Logical Index for each channel
  signal <- unlist(lapply(data, function(x, ...) {
    index <- logical(nrow(x))
    
    #Convert IPI.min and IPI.max to seconds 
    f_min <- floor(IPI.min / 1000)
    c_max <- ceiling(IPI.max / 1000)
    for(i in 1:nrow(x)){
      
      #Check for orion interal filtering
      if(x$Code[i] != 0) {
        index[i] <- TRUE
        next
      }
      
      #IPI Logical 
      td <- abs(x$Time[i] - x$Time)
      IPILogical <- td > f_min & td <= c_max
      
      #PPM LOgical
      pd <- abs(x$Power[i] - x$Power)
      PPMLogical <- pd <= PPM
      
      index[which(IPILogical & PPMLogical)] <- TRUE
    }
    index
  }, IPI.min = IPI.min, IPI.max = IPI.max, PPM = PPM))
  
  #Recombind data
  data <- rbindlist(data)
  
  #Filter Data
  filter(data, signal)
  } else{
    data
  }
}

# Creates a radio map by filtering orion mapping data by timestamp at known
# transmitter locations, then computes distance between transmitter location
#
# Args:
#   x: A POSIXct formated orion output
#   y: A POSIXct formated timestamp key
#   z: A key of antenna locations (ft)
#   int: int radiomapping sampling interval (sec)
#
# Returns: 
#   Orion contacts during periods of known transmitter coordinates, given 
#   antenna coordinates,  and the distance between transmitter and 
#   antenna 

CreateMap <- function(x, y, z, int) {
  #load required packages
  require(dplyr)
  require(dtplyr)
  require(data.table)
  
  x <- tbl_dt(x)
  y <- tbl_dt(y)
  z <- tbl_dt(z)
  z <- rename(z, ant_x = X, ant_y = Y)
  
  #Set mapping stop time
  y <- mutate(y, Stop = Start + int)
  
  #join mapping data and antenna a coordinates
  data <- inner_join(x, z)
  rm(x)
  rm(z)
  
  #Create table of positions at time
  y <- with(y, split(y, as.factor(paste(Start))))
  y <- lapply(y, function(x) {
    
    #Create a data table 
    Time <- seq.POSIXt(x$Start, x$Stop, by = "sec")
    trans_x <- rep(x$X, length(Time))
    trans_y <- rep(x$Y, length(Time))
    direction <- rep(x$Direction, length(Time))
    data.table(Time, direction, trans_x, trans_y)}
    )
  y <- rbindlist(y)
  
  #join data and times
  data <- inner_join(data, y)
  rm(y)
  
  #calculate distance
  data <- data %>%
    mutate(distance = sqrt((ant_x - trans_x)^2 + (ant_y - trans_y)^2)) %>%
    mutate(distance = signif(distance, 3))
}

#Assigns Animal IDs and trims orion tracking data by the dates collars were in 
#the enclosure
  # Args
  #   data: Orion data with Date and Time in POSIXct format in a singe column 
  #   named "Time"
  #   id_key: A table associating id, collar, orion frequency and start and stop
  #   of tracking. Start and stop formatted as POSIXct and named "start" and 
  #   "stop"
  # Returns
  #   Orion data with ids filtered by start and stop
    
AssignID <- function(data, id_key){
   #load required packages
  require(dplyr)
  require(dtplyr)
  require(data.table)
  
  data <- tbl_dt(data)
  id_key <- tbl_dt(id_key)
  
  #Create table of ids at time
  id <- with(id_key, split(id_key, 
                          list(as.factor(id), 
                               as.factor(Freq), 
                               as.factor(start)),
                          drop = TRUE))
  id <- lapply(id, function(x) {
    
    #Create a data table
    Time <- seq.POSIXt(x$start, x$stop, by = "sec")
    ID <- rep(x$id, length(Time))
    Freq <- rep(x$Freq, length(Time))
    data.table(Time, ID, Freq)})
  
  id <- rbindlist(id)
  
  #join id and data
  inner_join(id, data)
}

BuildJAGS <- function(df, model_dir, seed = NULL, ...) {
  require(rjags)
  require(MASS)
  require(MCMCpack)
  
  if(!is.null(seed)) {
    set.seed(seed)
  }
  
  #Format Data for BUGS
  p <- df$Power
  n <- length(p)
  n.site <- max(df$Site)
  n.ant <- length(unique(interaction(df$Ant, df$Site)))
  d <- ifelse(df$distance < 1, 1, df$distance)
  log10d <- 10 * log10(d)
  Site = df$Site
  Ant = AntEdit(df)
  
  #Define Hyperparameters
  #Bayesian Heirarchy by Ant and Site (Conjugate MVN prior on betas)
  Omega <- matrix(c(1,0,0,1), nrow=2)/10000
  prec.site <- matrix(c(1,0,0,1), nrow=2)/1000
  site.mu.0 <- c(0,0)
  
  BHR_data <- list(n = n, 
                   n.site = n.site, 
                   n.ant = n.ant,
                   p = p, 
                   Site = Site, 
                   Ant = Ant, 
                   log10d = log10d, 
                   Omega = Omega, 
                   prec.site = prec.site, 
                   site.mu.0 = site.mu.0)
  
  #Initialize parametes
  BHR_inits <- function () { list(
    beta.ant = MASS::mvrnorm(n.ant, c(0,0), matrix(c(10, 0, 0, 10), nrow = 2)),
    beta.site = MASS::mvrnorm(n.site, c(0,0), matrix(c(10, 0, 0, 10), nrow = 2)),
    R.site = MCMCpack::rwish(2, matrix(c(1,0,0,1), nrow=2)),
    tau.power = rgamma(n.ant, 1, 1))
  }
  
  BHR_params <- c("beta.ant", 
                  "beta.site", 
                  "R.site", 
                  "cov.site",
                  "tau.power", 
                  "var.power")
  
  #Specify Model in JAGS
  BHR_model <- jags.model(file = model_dir,
                          data = BHR_data, 
                          inits = BHR_inits, 
                          n.chains = 3,
                          n.adapt = 1000)
  #Burn In
  update(BHR_model, 1000)
  
  #coda fomratted chains
  samples <- coda.samples(BHR_model, 
                          variable.names = BHR_params,
                          n.iter = 10000)
  samples
}

#Format Coda Sample means as a list af matrices (similar to the ouptput of Coefs function)
JAGSCoefs <- function(samples){
  require(stringr)
  require(rjags)
  sample_stats <- summary(samples)$statistics
  labels <- rownames(sample_stats)
  coef_names <- unique(str_extract(labels, "[[:alpha:]]+.[[:alpha:]]+"))
  coefs <- list()
  for (i in coef_names) {
    coef_index <- str_detect(labels, i)
    coef_est <- sample_stats[coef_index, "Mean"]
    nr <- max(as.numeric(str_extract(labels[coef_index], "\\d++")))
    coefs [[i]] <- matrix(coef_est, nrow = nr)
  }
  coefs
}

#FixeAnt swaps Site 1 Ant 6 and Site 1 and 7 labels.  Raw data indicates that
#phycials lines were switched at power inserter or antenna switch
FixAnt <- function(data){
  replace <- logical(nrow(data))
  replace[data$Site == 1 & data$Ant == 6] <- TRUE
  data[data$Site == 1 & data$Ant == 6,"Ant"] <- 7
  data[data$Site == 1 & data$Ant == 7 & replace == FALSE,"Ant"] <- 6
  data
}

#define TDiff function, calculates time nearest time in b and time difference
#Args 
#a: A data table with a POSIXct column labeled time
#b: A data table with a POSIXct column labeled Time
#Returns
#A with columns for nearest time in b and time difference between a and
#nearest time in b
TDiff <- function(a, b){
  require(dplyr)
  #Allocate memory for for loop
  t <- as.POSIXct(rep(NA, nrow(a)), tz = tz(a$time))
  d <- rep(NA, nrow(a))
  attributes(d)$units <- "mins"
  attributes(d)$class <- "difftime"
  
  for(i in seq_along(a$time)) {
    
    #Find difference
    d_sign <- abs(difftime(a$time[i], unique(b$Time), units = "mins"))
    d[i] <- min(d_sign)
    
    #Find automated time
    dt <- b %>%
      distinct(Time) %>%
      filter(d_sign == d[i])
    t[i] <- dt$Time
  }
  
  #add automated time and time diff back to manual data
  mutate(a, Time = t, time_diff = d)
}

#For each Manual fix find nearest Automated Fix in time
# Args
#   ARTS_data: Filtered automated telemetry data
#   Manual_data: Tidied manual telemetry data
# Returns
#   Manual telemetry data school with columns for closest Automated telemetry
#   fix and time difference
ManualvsARTS_Time <- function(ARTS_data, Manual_data, cluster = NULL){
  require(lubridate)
  require(dplyr)
  
  #Check intersection of IDs between manual and automated data
  common_ids <- intersect(unique(Manual_data$id), unique(ARTS_data$ID))
  
  #select times for common ids
  Manual_times <- Manual_data %>% 
    filter(id %in% common_ids) %>%
    arrange(id, time)
  ARTS_times <- ARTS_data %>% 
    filter(ID %in% common_ids) %>% 
    distinct(ID, Time) %>%
    arrange(ID, Time)
  
  #Spilt data by id
  Manual_times <- with(Manual_times, split(Manual_times, id))
  ARTS_times <- with(ARTS_times, split(ARTS_times, ID))
  
  #Determine how many cores to use
  if(is.null(cluster)) {
    #Calculate closest automated fix and time difference
      results <- mapply(TDiff, a = Manual_times, b = ARTS_times)
  } else {
    require(parallel)
    clusterEvalQ(cluster,
                 {library(here)
                  library(dplyr)
                  library(dtplyr)
                  library(data.table)
                  library(lubridate)
                  source(here::here("src", "ARTSFunctions.R"))
                 }
    )
    results <- clusterMap(cluster, TDiff, a = Manual_times, b = ARTS_times)
  }
  rbindlist(results)
}

#EditAnt: Edit antenna notation so each antenna in the Orion dataset is uniquely 
#identified
# Args
#   x: An orion output
#
# Returns
#   x$Ant:  A vector of edited antenna labels.  
#     Site 1: Ant 1-8
#     Site 2: Ant 9-16
#     Site 3: Ant 17-24
#     Site 4: Ant 25-32

AntEdit <- function(x) {
  ant <- 9
  for (i in 2:4){
    for(j in 1:8){
      x[x$Site == i &
          x$Ant == j, "Ant"] <- ant
      ant <- ant + 1
    }
  }
  x$Ant
}

# CalcTableBayes: Calculates lognormal pdf of positon for each antenna at each
# received power within the enclosure.
#   Args
#     model: Bayesian model coeffecients
#     antkey:  Key of antenna locations
#     
#   Returns
#     tables: An array with the dimensions [x, y, Power] containing the normalized
#     densities of position.
CalcTablesBayes <- function(model, antkey){
  pdf.info <- list()
  grid.x <- 0:55
  grid.y <- 0:74
  power.range <- -57:-113
  
  for(i in 1:32){
    grid <- outer(grid.x, grid.y, function(a,b) {
      sqrt((a-antkey$X[i])^2+(b-antkey$Y[i])^2)
    })
    mean.p0 <- model$beta.ant[i,1]
    mean.eta <- model$beta.ant[i,2]
    mean.var <- 1/model$tau.power[i]
    densities <- array(0, c(length(grid.x), length(grid.y),length(power.range)), 
                       dimnames = list(grid.x, grid.y, power.range))
    
    for(j in 1:length(power.range)){
      mu <- (power.range[j] - mean.p0)/(10*mean.eta)*log(10)
      sd <- abs(sqrt(mean.var)/(10*mean.eta)*log(10))
      dens <- dlnorm(grid, mu, sd)
      densities[,,j] <- dens/sum(dens)
    }
    pdf.info[[i]] <- list(p0 = mean.p0, eta = mean.eta, var = mean.var, 
                          grid = grid, densities = densities)
  }
  names(pdf.info) <- 1:32
  pdf.info
}

# CalcTablesNP: Calculates nonparameteric pdf of positon for each antenna at each
# received power within the enclosure.
# 
#   Args
#     data: Radiomapping data
#     
#   Returns
#     tables: An array with the dimensions [x, y, Power] containing the normalized
#     densities of position.
CalcTablesNP <- function(data){
  require(ks)
  
  smoothing.data<- split(data, as.factor(data$Ant))
  
  #format for kde
  smoothing.data<- lapply(smoothing.data, function(x){
    matrix(c(x$trans_x, x$trans_y, x$Power), nrow(x), ncol = 3)
  }
  )
  
  #evaluation grid
  grid.x <- 0:55
  grid.y <- 0:74
  power.range <- -57:-113
  grid.xyz <- expand.grid(grid.x, grid.y, power.range)
  
  #fit kde
  emp.dens <- lapply(smoothing.data, function(x){
    kde(x, eval.points = grid.xyz)
  }
  )
  
  #organize kde data
  dens.array.list <- lapply(emp.dens, function(x){
    H <- x$H
    dimnames(H) <- list(c("x", "y", "power"), c("x", "y", "power"))
    KDE <- array(data = x$estimate,
                 dim = c(length(grid.x), 
                         length(grid.y), 
                         length(power.range)), 
                 dimnames = list(grid.x, grid.y, power.range))
    list(H = H, KDE = KDE)
  }
  )
  
  #Normalize Densities
  dens.array.list <- lapply(dens.array.list, function(x){
    for(i in 1:dim(x$KDE)[3]){
      x$KDE[,,i] <- x$KDE[,,i]/sum(x$KDE[,,i])
    }
    list(H = x$H, KDE = x$KDE)
  }
  )
  dens.array.list
}

# Top3: Filters 3 strongest signals 
# Args
#   data: A list consisting of lists of Orion contants by time for a single 
#         freqency
# Returns
#   A a list consisting of list of the three stongest orion contacts at each 
#   timestamp
Top3 <- function(data) {
  if(length(unique(data$Time)) > 1) {
    stop("More than one time point detected")
  }
  
  data <- data[order(data$Power, decreasing = TRUE), ]
  a <- data[1, ]
  b <- data[!(data$Ant == a$Ant), ]
  b <- b[order(b$Power, decreasing = TRUE), ]
  b <- b[1, ]
  c <- data[!(data$Ant == a$Ant) & !(data$Ant == b$Ant), ]
  c <- c[order(c$Power, decreasing = TRUE), ]
  c <- c[1, ]
  y <- rbind(a, b, c)
  y
}

#PositionPDF: Calculates the PDF of position within the enclosure, normalized to
#enclosure 
# Args:
#   data: A dataframe of Orion contacts at a single time 
#   tables: A list of the array of PDFs of posistion within the enclosures by 
#   antenna.  
#           
# Returns
#   A list of PDFs of position for each timestamp or "Numeric Underflow" if 
#   the PDF collapses to 0
PositionPDF <- function (data, tables){
  dens <- list()
  for (i in 1:nrow(data)) {
    #list index for antenna info
    ant <- data$Ant[i]
    
    #list index for table postion in antenna info
    t <- length(tables[[ant]])
    
    #array index for density of desired Power by name
    d <- paste(data$Power[i])
    
    #desired density  
    dens[[i]] <- tables[[ant]][[t]][,,d]
  }
  
  position.pdf <- Reduce("*", dens)
  position.pdf <- position.pdf / sum(position.pdf)
  if (any(is.na(position.pdf))) {
    warning("Numeric Underflow")
  }
  position.pdf
}

# MAP.est: Calculates the maximum and entropy of a pdf a given timestamp
# Args
#   pdf: A list of pdfs by timestamp
# Returns
#   A dataframe containing the cartesian coordiante of the maximum and the 
#   shannon entropy of the distribution

MAP.est <- function(a) {
  if (length(a) == 0L | (any(is.na(a)))) {
    x <- NA
    y <- NA
    E <- NA
    df <- data.frame(x=x, y=y, entropy=E)
    warning("Numeric Underflow")
  } else {
    #row (x), col (y) index of the max density
    max<- which(a == max(a), arr.ind = TRUE)
    
    #values at the max index
    x <- as.numeric(dimnames(a)[[c(1, max[1,1])]])
    y <- as.numeric(dimnames(a)[[c(2, max[1,2])]])
    
    #Need a log function which returns zero when the pdf is 0 (for the case
    # 0 * log(0)), instead of -inf.
    log0 <- function(p){
      y <- ifelse(p == 0, 0, log(p))
      y
    }
    
    E <- -1 * sum(a * log0(a))
    df <- data.frame(x=x, y=y, entropy=E)
  }
  df
}

#Format MAP estimates into data frames
FormatMAP <- function(x) {
  require(lubridate)
  if(!is.list(x)){
    stop("x must be a list ")
  }
  #Freq i
  for (i in seq_along(x)){
    
    #Time j
    
    for(j in seq_along(x[[i]])) {
      x[[c(i,j)]]$Time <- ymd_hms(names(x[[i]][j]), 
                                  tz = "America/Chicago")
      x[[c(i,j)]]$Freq <- as.numeric(names(x[i]))
    }
  }
  x <- lapply(x, rbindlist)
  rbindlist(x) %>%
    as.tibble()
}


#Trilaterate predicts location gien a model of signal decay and an Orion
#output
Trilaterate <-function(data, model){
  require(dplyr)
  require(minpack.lm)
  
  model <- tbl_dt(model$beta.ant)
  names(model) <- c("beta0", "beta1")
  model <- mutate(model, Ant = 1:32)
  data <- inner_join(data, model) %>%
    mutate(pred_d = 10^((Power - beta0)/(beta1 *10)))
  
  result <- nlsLM(pred_d ~ sqrt((x - ant_x)^2 + (y - ant_y)^2), 
                  data = data, 
                  start = list(x = mean(data$ant_x), y = mean(data$ant_y)), 
                  upper= c(180*0.3048, 240*0.3048),
                  lower= c(0,0),
                  weights = 1/(pred_d^2),
                  control = nls.lm.control(maxiter = 100))
}


#Format trilateration estimates into data frames
FormatNLS <- function(x) {
  require(lubridate)
  
  x <- lapply(x, function(x) {
    lapply(x, function(x) {
      if(class(x) == "try-error") {
        x <- data.frame(x = NA, y = NA)
      } else {
        data.frame(t(coef(x)))
      }
    }
    )
  }
  )
  
  #Freq i
  for (i in seq_along(x)){
    
    #Time j
    for(j in seq_along(x[[i]])) {
      x[[c(i,j)]]$Time<- ymd_hms(names(x[[i]][j]),
                                 tz = "America/Chicago")
      x[[c(i,j)]]$Freq <- as.numeric(names(x[i]))
    }
  }
  x <- lapply(x, rbindlist)
  x <- rbindlist(x)
  x
}

# CalcDist: Calculates predicted distance form lognormal pdf of positon for each 
# antenna at each teceived power within the enclosure.
#   Args
#     data: Data frame containg received Powers and uniquely identified antenna
#     model: Bayesian model coeffecients
#     
#   Returns
#     tables: A dataframe containing estimates of distance from the mode,
#     median, and mean of the posterior predictive of distace
CalcDist <- function(df, model){
  require(dplyr)
  coeffs <- data.frame(Ant = 1:32,
                       mean.p0 = model$beta.ant[,1],
                       mean.eta = model$beta.ant[,2],
                       mean.var = 1/model$tau.power)
  
  df <- inner_join(df, coeffs)
  
  df <- mutate(df, 
               logmu = (df$Power - mean.p0)/(10*mean.eta)*log(10),
               logsd =abs(sqrt(mean.var)/(10*mean.eta)*log(10)),
               mode = exp(logmu - logsd^2),
               median = exp(logmu),
               mean = exp((logmu + (logsd^2)/2))) 
}

CalcMode <- function(x){
  d <- density(x, na.rm = T)
  d$x[which(d$y == max(d$y))]
}

#Assigns Animal IDs and trims orion tracking data by the dates collars were in 
#the enclosure
# Args
#   data: Orion data with Date and Time in POSIXct format in a singe column 
#   named "Time"
#   id_key: A table associating id, collar, orion frequency and start and stop
#   of tracking. Start and stop formatted as POSIXct and named "start" and 
#   "stop"
# Returns
#   Orion data with ids filtered by start and stop
#   
AssignID <- function(data, id_key){
  #load required packages
  require(dplyr)
  require(dtplyr)
  require(data.table)
  
  data <- tbl_dt(data)
  id_key <- tbl_dt(id_key)
  
  #Create table of ids at time
  id <- with(id_key, split(id_key, 
                           list(as.factor(id), 
                                as.factor(Freq), 
                                as.factor(start)),
                           drop = TRUE))
  id <- lapply(id, function(x) {
    
    #Create a data table
    Time <- seq.POSIXt(x$start, x$stop, by = "sec")
    ID <- rep(x$id, length(Time))
    Freq <- rep(x$Freq, length(Time))
    data.table(Time, ID, Freq)})
  
  id <- rbindlist(id)
  
  #join id and data
  inner_join(id, data)
}

#Retreive Highest Posterior Density Contour
#Function from https://www.star.le.ac.uk/sav2/R/plot_contour.R
get.levels2d <- function(img, prob=0.95) {
  
  # check the inputs
  
  if (missing(img)) stop('Missing data IMG.')
  
  if (!exists("z", where=img)) stop('IMG input does not contain z array.')
  if (!exists("x", where=img)) stop('IMG input does not contain x vector.')
  if (!exists("y", where=img)) stop('IMG input does not contain y vector.')
  
  # probability should be between 0 and 1  
  
  if (max(prob) > 1) stop('PROB > 1 error.')
  if (min(prob) < 0) stop('PROB < 0 error.')
  
  dx <- diff(img$x[1:2])
  dy <- diff(img$y[1:2])
  norm <- sum(img$z) 
  sz <- sort(img$z)
  c1 <- cumsum(sz)/norm * dx * dy
  clevels <- approx(c1, sz, xout = 1 - prob)$y
  
  return( clevels )
}

#Takes a single isopleth generated from contourLines and reformats it as a SpatialPolygons object with appropriate islands and wholes
#If no isopleth is present the enclosure boundry is used
FormatContour <- function(contour, x_bounds, y_bounds){
  require(sp)
  require(maptools)
  require(cleangeo)
  
  level <- unique(sapply(contour, `[[`, 1))
  #Check that only a single contour level is being plotted
  if(length(level) > 1) {
    stop("More than one contour level selected")
  } 
  #If the contour contains no boundries use the enclosure boundry
  segs <- length(contour)
  if (segs == 0) {
    
    level <- 0
    coords <- cbind(x_bounds, y_bounds)
    contour <- list(Polygon(coords))
    
  } else {
    
    #Merge Contour line segments
    contour <- lapply(contour, function(x){
      coords <- cbind(x$x, x$y)
      Line(coords)
    })
    contour <- Lines(contour, ID = paste(level))
    contour <- SpatialLines(list(contour))
    contour <- gLineMerge(contour)
    
    #Generate Polygons that describe the isopleth
    contour <- lapply(contour@lines[[1]]@Lines, function(x){
      coords <- slot(x, "coords")
      Polygon(coords)
    }
    )
  }
  
  #Clean Polygons
  if(length(contour) > 1) {
    
    contour <- lapply(contour, function(x){
      
      #Format as spatial polygons for intersection testing and union later
      x <- SpatialPolygons(list(Polygons(list(x), ID = NA)))
      report <- clgeo_CollectionReport(x)
      if(report$valid == FALSE) {
        x <- clgeo_Clean(x)
      }
      x
    }
    )
    
    #Clear Null Elements
    contour <- contour[!sapply(contour, is.null)]
    
    #If the contour is NULL use the enclosure boundry
    if(length(contour) == 0){
      level <- 0
      coords <- cbind(x_bounds, y_bounds)
      contour <- list(Polygon(coords))
    }
    
    #Test for intersecting polygons
    n_polygons <- length(contour)
    if(n_polygons > 1){
      poly_comb <- combn(n_polygons, 2)
      i <- 1
      
      while(i <= ncol(poly_comb)) {
        poly1 <- contour[[poly_comb[1,i]]]
        poly2 <- contour[[poly_comb[2,i]]]
        
        
        if(gCrosses(poly1, poly2)) {
          
          contour[[n_polygons + 1]] <- gUnion(poly1, poly2)
          contour <-  contour[-poly_comb[,i]]
          
          #reset stop condition
          n_polygons <- length(contour)
          if(n_polygons == 1) {
            i <- ncol(poly_comb) + 1
          } else{
            poly_comb <- combn(n_polygons, 2)
            i <- 1
          }
        } else {
          #advance stop condition
          i <- i + 1
        }
      }
    } else {
      contour <- list(contour[[1]]@polygons[[1]]@Polygons[[1]])
    }
    
  }
  
  #Extract Polygons fom SpatialPolygons if required
  if(any(sapply(contour, class) == "SpatialPolygons")){
    contour <- lapply(contour, function(x){
      if(class(x) == "SpatialPolygons") {
        x@polygons[[1]]@Polygons[[1]]
      }
    }
    ) 
  }
  
  #Assemble all Polygon objects into one Polygons object
  contour <- Polygons(contour, ID = level)
  
  #Assign Island and Holes to polygons
  contour <- checkPolygonsHoles(contour, properly = FALSE)
  
  #Format as Spatial Polygons
  SpatialPolygons(list(contour))
}

ARTSPrediction <- function(data, model_tables) {
  #Splitting data by Freq and Time
  fixes <- split(data, data$Freq)
  fixes <- lapply(fixes, 
                  function(x) x[order(x$Time),])
  fixes <- lapply(fixes, function(x) {
    split(x, as.factor(as.character(x$Time)))
  })
  fixes <- lapply(fixes, function(x){
    x[which(sapply(x, function(x) {
      if(any(names(x) %in% c("trans_x", "trans_y"))){
        length(unique(x$Ant)) >= 3 &
        length(unique(interaction(x$trans_x, x$trans_y))) == 1
      } else {
        length(unique(x$Ant)) >= 3 
      }
    }
    ))]
  }
  )
  
  #Select 3 strongest contacts
  prediction <- lapply(fixes, function(x) {
    lapply(x, function(x) {
      prediction  <- Top3(x)
      prediction <- PositionPDF(prediction, tables = model_tables)
      MAP.est(prediction)
    }
    )
  }
  )
  #Format fitting varialbe names
  FormatMAP(prediction)
}

#Adapted from Seilmayer and Ratajczak (2017) and Zechmeister and Kuster (2009)
#Calculates the Generalized Lomb-Scargle periodogram.  
# Arguements
#   y: Numeric Response variable
#   t: Numeric time
#   f: frequency grid
#   weights: Numeric vector of observations weights/variances
#   intercept: Logical is observation window being fit (Classic LS)?  Note to 
#   calculate the LS periodogram for the observation window set weights to 1, 
#   code y = 1 for each t, fit model without interecpt.
# Returns
#   f: vector of fit frequencies
#   PSD:  Power Spectral Density normalized to sample variance.  Ranges from 0 
#   to 1, equivalent to model r^2.
#   a <- cosine term coefficeint
#   b <- sine term coefficeint
#   c <- interecpt term
#   A <- sine function amplitude
#   phi <- sine function phase 

GenLS <- function(y, t, f, weights = NULL, intercept = TRUE) {
  
  nt <- length(t)
  
  if(is.null(weights)) {
    weights <- rep(1, nt) 
  }  
  w <- (1/weights)/sum(1/weights)
  
  omega <- 2 * pi * f
  omega_t <- t %*% t(omega) 
  
  m =  nrow(omega_t)
  n = ncol(omega_t)
  
  cos <- cos(omega_t)
  
  #sin(x)^2 + cos(x)^2 = 1
  sin <- sqrt(1 - cos^2)
  
  #Calculate Tau
  #Classic LS Terms
  CC <- .colSums(w*cos^2, m = m, n = n)
  SS <- .colSums(w*sin^2, m = m, n = n)
  CS <- .colSums(w * cos * sin, m = m, n = n)
  
  #Generalized LS Terms
  if (intercept) {
    C <- .colSums(w*cos, m = m, n = n)
    S <- .colSums(w*sin,  m = m, n = n)
    
    CC <- CC - C^2
    SS <- SS - S^2
    CS <- CS - C * S
  } 
  
  omega_tau <- atan((2 * CS)/ (CC - SS)) / (2)
  
  #Calculate/Recalculate terms centered at tau
  cos <- cos(omega_t - omega_tau)
  sin <- sqrt(1 - cos^2)
  
  CC <- .colSums(w*cos^2, m = m, n = n)
  SS <- .colSums(w*sin^2, m = m, n = n)
  CS <- .colSums(w * cos * sin, m = m, n = n)
  
  YY <- sum(w*y^2)
  YC <- .colSums(w*y*cos, m = m, n = n)
  YS <- .colSums(w*y*sin, m = m, n = n)
  
  Y <- sum(w*y)
  C <- .colSums(w*cos, m = m, n = n)
  S <- .colSums(w*sin,  m = m, n = n)
  
  #Generalized LS Terms
  if (intercept) {
    CC <- CC - C^2
    SS <- SS - S^2
    CS <- CS - C * S
    
    YY <- YY - Y^2
    YC <- YC - Y * C
    YS<- YS - Y * S 
  } 
  
  D <- CC * SS - CS^2
  
  PSD <- 1/(YY) * (YC^2/CC + YS^2/SS)
  a <- 1/D * (YC * SS - YS * CS)
  b <- 1/D * (YS * CC - YC * CS)
  c <- Y - a * C - b * S
  A <- sqrt(a^2 + b^2)
  phi <- omega_tau
  
  data.frame(f, PSD, a, b, c, A, phi)
}

#Fourier Window
#Calculates the exact Fourier Window for a sample, normalized by 2/(y^2) to 
#compare to the normalized LSP, (equivalent to fitting the LSP to a uniformly
#sampled indicator fucntion (0 at no observation, 1 at observation), but much 
#faster to calculate)
FourierWindow <- function(t, f) {
  omega <- 2 * pi * f
  omega_t <- t %*% t(omega) 
  
  W <- exp(-1i*omega_t)
  W <- abs(colSums(W))
  (1/(length(t))*W)^2
} 

#Define functions to calculate FAP approximation from Baluev 2008 using
#Davies bound
#standard normalization is Z = 2/NH * z_1
FAP_single <- function(z, d_k, N){
  (1 - z)^((N-d_k)/2)
}

P_single <- function(z, d_k, N){
  1 - FAP_single(z, d_k, N)
}

FAP_approx <- function (z, N, d_k, fmax, T) {
  N_eff <- fmax * T
  1 - P_single(z, d_k, N)^N_eff
}

FAP_davies <- function(z, N, d_k, d_h, var_t, fmax) {
  T_eff <- sqrt(4 * pi * var_t)
  W <- fmax * T_eff
  A_fmax <- 2 * pi^(3/2) * W
  
  gamma <- exp(lgamma((N - d_h)/2) - lgamma((N - d_k+1)/2))
  
  Tau <- {
    gamma / (2 * pi) *
      A_fmax * 
      (z / pi)^((d_k - d_h - 1)/2) * 
      (1 - z)^((N - d_k -1)/2)
  }
  1 - P_single(z, d_k, N) * exp(-Tau)
}

#Search perodgram peaks for window peaks and 1st - 3rd harmonics and removes them
FilterHaromics <- function(var_ctps, win_ctps, n, ...) {
  require(dplyr)
  
  har <- seq(-n, n)
  har <- har[har != 0]
  
  index <- logical(length(var_ctps))
  for (i in seq_along(var_ctps)){
    
    index[i] <-
    
    #check if peak is in window
    !(any(near(var_ctps[i], win_ctps, ...)) |
    #or if there are peaks at multiple of window peaks
      any(near(var_ctps[i] - 3 * win_ctps, var_ctps, ...)) |
      any(near(var_ctps[i] - 2 * win_ctps, var_ctps, ...)) |
      any(near(var_ctps[i] - 1 * win_ctps, var_ctps, ...)) |
      any(near(var_ctps[i] + 1 * win_ctps, var_ctps, ...)) |
      any(near(var_ctps[i] + 2 * win_ctps, var_ctps, ...)) |
      any(near(var_ctps[i] + 3 * win_ctps, var_ctps, ...))) 
  }
  var_ctps[index]
}
  
  
  
  
  

  
```
