## Supplementary material for "An Automated Radio-Telemetry System (ARTS) for Monitoring Small Mammals": Analysis R Code: Filtering.R

```
#Load required packages 
library(parallel)
library(tidyverse)
library(data.table)
library(dtplyr)
library(lubridate)
library(here)

#Source functions
source(here::here("src", "ARTSFunctions.R"))

#Path to raw Orion data
rawdata_path <- here::here("data",
                           "Raw Orion Automated Tracking Data")

#Path to mapping key
mapkey_path <- here::here("data", 
                          "All Radiomapping Keys", 
                          "AVA2014 freq concatenated mapping key.csv")

#Path to antenna key
antennakey_path <- here::here("data", 
                              "All Radiomapping Keys", 
                              "Antenna Key.csv")

#Path to id key
idkey_path <- here::here("data", "2014 Animal Data", "freqs 2014.csv")

#Look at file structure of raw data
list.files(rawdata_path, pattern =".txt", recursive =TRUE, include.dirs = TRUE)

#1:284 tracking files
tracking.files <- list.files(rawdata_path, patter =".txt", 
                             recursive =TRUE, 
                             include.dirs = TRUE)[1:284]

#c(285:323, 336:370) mapping files (not including 2014-07-21 beacon mapping)
mapping.files <- list.files(rawdata_path, patter =".txt", 
                             recursive =TRUE, 
                             include.dirs = TRUE)[c(285:323, 336:370)]


#Set up cluster for parallel computation
cl <- makeCluster(detectCores() - 1)
clusterEvalQ(cl,
             {source(here::here("src","ARTSFunctions.R"))
              library(parallel)
              library(tidyverse)
              library(dtplyr)
              library(data.table)
              library(lubridate)})

#Load Mapping data
Mapping_2014 <- lapply(mapping.files,
                       function(x, path) {
                         read.table(paste(path, x, sep = "/"), 
                           stringsAsFactors = FALSE,
                           header = TRUE,
                           fill = TRUE,
                           sep = "")}, path = rawdata_path)

#Load Tracking data
Tracking_2014 <- lapply(tracking.files,
                           function(x, path) {
                             read.table(paste(path, x, sep ="/"), 
                               stringsAsFactors = FALSE,
                               header = TRUE,
                               fill = TRUE,
                               sep = "")}, path = rawdata_path)

#Load mapping key
map_key <- read_csv(file = mapkey_path, 
                     col_types = cols(Start = col_character(), 
                                      Stop = col_character()))

#Load antenna key
ant_key <- read_csv(antennakey_path) 

#Load id key
idkey <- read_csv(idkey_path, 
                  col_types = cols(manual_dead = col_character(), 
                  start = col_character(), stop = col_character()))

#Discard Empty Data tables
Tracking_2014 <- 
  Tracking_2014[lapply(Tracking_2014, function(x) nrow(x) != 0) == TRUE] 

Mapping_2014 <- 
  Mapping_2014[lapply(Mapping_2014, function(x) nrow(x) != 0) == TRUE] 
                                                  
#Format Times in Tracking_2014, Mapping_2014, map_key and idkey
Tracking_2014 <- parLapply(cl, Tracking_2014, function (x){
  x %>% 
    mutate(Time = ymd_hms(paste(Date, Time))) %>%
    mutate(Time = force_tz(Time, tz = "America/Chicago")) %>%
    dplyr::select(-Date)})

#Note 5 failures to parse in "10 28 2014/mmcdump 10-28-2014 SW PM.txt"
#At rows 723760 723799 723800 723801 723802 several variables are out of 
#normal ranges, these rows should be caught by FilterNoise()

Mapping_2014 <- parLapply(cl, Mapping_2014, function (x){
  x %>% 
    mutate(Time = ymd_hms(paste(Date, Time), tz = "America/Chicago")) %>%
    mutate(Time = force_tz(Time, tz = "America/Chicago")) %>%
    dplyr::select(-Date)
})

map_key <- map_key %>% 
  mutate(Start = mdy_hms(paste(Date, Start, tz = "America/Chicago"))) %>%
  mutate(Start = force_tz(Start, tz = "America/Chicago")) %>%
  dplyr::select(-Date, -Stop)

idkey <- idkey %>%
  mutate(start = ymd_hms(start, tz = "America/Chicago"),
         manual_dead = ymd_hms(manual_dead, tz = "America/Chicago"),
         stop = ymd_hms(stop)) %>%
  mutate(start = force_tz(start, tz = "America/Chicago"),
         manual_dead = force_tz(manual_dead, tz = "America/Chicago"),
         stop = force_tz(stop, tz = "America/Chicago")) 


#Filter Tracking_2014 and Mapping_2014
Tracking_2014 <- parLapply(cl,
                           Tracking_2014, 
                           FilterNoise, 
                           IPI.min = 1500, 
                           IPI.max = 2300,
                           PPM = 6)

Mapping_2014 <- parLapply(cl,
                          Mapping_2014, 
                          FilterNoise, 
                          IPI.min = 1500, 
                          IPI.max = 2300,
                          PPM = 6)

#Format tracking and mapping as data tables 
#may be be slow filtering but will allow removal of duplicate contact is 
#seperate files from MMC card writing errors
Mapping_2014 <- rbindlist(Mapping_2014)
Tracking_2014 <- rbindlist(Tracking_2014)

#Remove Duplicates
Mapping_2014 <- distinct(Mapping_2014)
Tracking_2014 <- distinct(Tracking_2014)

#Convert feet to meters in ant key and map key
ant_key <- mutate(ant_key, X = X * 0.3048, Y = Y * 0.3048)
map_key <- mutate(map_key, X = X * 0.3048, Y = Y * 0.3048)

#Switch antenna 1.6 and 1.7 in Map
Mapping_2014 <- FixAnt(Mapping_2014)

#Switch antenna 1.6 and 1.7 in Tracking data
Tracking_2014 <- FixAnt((Tracking_2014))

#Radiomapping key looks like Y (Easting) coordinate is shifted 10 feet in the 
#positive direction (East)
map_key_shift <- mutate(map_key, Y = ifelse(X > 90 * 0.3048, 
                                            Y - 10 * 0.3048,
                                            Y))

#Create Radiomaps 
Mapping_2014_raw <- CreateMap(Mapping_2014, map_key, ant_key, 36)
Mapping_2014_shift <- CreateMap(Mapping_2014, map_key_shift, ant_key, 36)

#Assign IDs to Tracking data
Tracking_2014 <- AssignID(Tracking_2014, idkey)

#Save Data (note that write_csv() converts datetimes to UTC)
write_csv(Tracking_2014,  here::here("results", "Tracking_2014.csv"))
write_csv(Mapping_2014_raw, here::here("results", "Mapping_2014_raw.csv"))
write_csv(Mapping_2014_shift, here::here("results","Mapping_2014_shift.csv"))

#stopCluster
stopCluster(cl)

 
```
