## Supplementary material for "An Automated Radio-Telemetry System (ARTS) for Monitoring Small Mammals": Analysis R Code: JAGS_Regression.R

```
#Load packages
library(here)
library(tidyverse)
library(rjags)
library(ggmcmc)
source(here::here("src", "ARTSFunctions.R"))

training_data <- c("raw", "shift", "TrackingAug", "TrackingAug2")
model_dir = here::here("src", "model.4.txt")

for (i in training_data){
  #Load Training data
  training_data <-
    switch(i,
           raw = here::here("results", "Mapping_2014_raw.csv"),
           shift = here::here("results", "Mapping_2014_shift.csv"),
           TrackingAug = here::here("results", "Mapping_2014_TrackingAug.csv"),
           TrackingAug2 = here::here("results", "Training_TrackingAug2.csv")
    ) %>%
    read_csv(.) %>%
    mutate(Ant = AntEdit(.))
  
  samples <- BuildJAGS(training_data, model_dir, seed = 1234)
  
  if(!dir.exists(here::here("docs", "JAGS_Regression"))) {
    dir.create(here::here("docs", "JAGS_Regression"))
  }
  
  diagnostic_dir <- here::here("docs", 
                               "JAGS_Regression", 
                               paste0(i, "_ggmcmc.pdf"))
  #Diagnostic plot in ggmcmc
  ggmcmc(ggs(samples),
         file = diagnostic_dir ,
         param_page = 6)
  
  assign(paste0(i, "_BHR_model_coeffs"),  JAGSCoefs(samples))
  save(list = paste0(i, "_BHR_model_coeffs"), 
       file = here::here("results", paste0(i, "_BHR_model_coeffs.RData")))
  rm(list = c("samples", paste0(i, "_BHR_model_coeffs")))
}
```
