## Supplementary material for "An Automated Radio-Telemetry System (ARTS) for Monitoring Small Mammals": Analysis R Code: Lomb-Scargle_Analysis.R

```
library(here)
library(data.table)
library(dtplyr)
library(tidyverse)
library(lubridate)
library(features)
source(here::here("src", "ARTSFunctions.R"))

# Load ARTS predictions
ARTS_predictions <- read_csv(here::here("results", "ARTS_predictions_2014.csv")) 

week2 <- ARTS_predictions %>%
  mutate(Time = with_tz(Time, tzone = "America/Chicago"),
         dawn = with_tz(dawn, tzone = "America/Chicago"),
         sunrise = with_tz(sunrise, tzone = "America/Chicago"),
         sunset = with_tz(sunset, tzone = "America/Chicago"),
         dusk = with_tz(dusk, tzone = "America/Chicago")) %>%
  filter(Time > ymd_hms("2014-10-7 00:00:00", tz = "America/Chicago"),
         Time < ymd_hms("2014-10-13 11:59:59", tz = "America/Chicago"))

#Frequency grid to scan, oversampling factor of 10, T = 1 week = 7*24*60*60 sec,
#f_max = 1/120 Hz, scan from f_min = 1/week to fmax with oversampling factor of
#5
fmax <- 1/120
T <- 7*24*60*60
ofac <- 5

f_grid <- seq(1/T, fmax, length.out= ofac*T*fmax)
T_grid <- 1/f_grid

LS_data <- list()
variables <- c("speed", "nest_dist")
for(i in variables) {
  #Let d_h = number of parameters in H0 and d_k = number of parameters in HA
  #Generalized LS Peroidograms fit to step(RobPer)
  LS_data[[paste0("step_" ,i)]] <-  week2 %>%
    mutate(Time = as.numeric(Time)) %>%
    split(.$ID) %>%
    lapply(function(x) { 
      require(RobPer)
      df <- data.frame(Time = x$Time, var = pull(x, i))
      RobPer(df,
             weighting = FALSE,
             periods = T_grid,
             model = "step",
             regression = "L2") %>%
        data.frame(PSD = ., 
                   ID = unique(x$ID),
                   period = T_grid,
                   method = "step",
                   variable = i,
                   d_h = 1,
                   d_k = 10) 
    }
    )
  
  #Generalized LS fit using GenLS
  LS_data[[paste0("GenLS_" ,i)]] <- 
    week2 %>%
    mutate(Time = as.numeric(Time)) %>%
    split(.$ID) %>%
    lapply(function(x) {
      GenLS(y = x[,i, drop = TRUE],
            t = x$Time,
            f = f_grid) %>%
        mutate(ID = unique(x$ID),
               period = T_grid,
               method = "GenLS",
               variable = i,
               d_h = 1,
               d_k = 3) 
    }
    )
} 
 
#Sampling Window generalized LS (RobPer)
LS_data[["step_window"]]<- 
  week2 %>%
  mutate(Time = as.numeric(Time)) %>%
  split(.$ID) %>%
  lapply(function(x) {
    require("RobPer")
    time = seq(min(x$Time), max(x$Time), by = 60)
    df <- data.frame(time,
                      contact = time %in% as.numeric(x$Time))
    RobPer(df,
            weighting = FALSE,
            periods = T_grid, 
            model = "step",
            regression = "L2") %>%
      data.frame(PSD = .,
                 ID = unique(x$ID),
                 period = T_grid,
                 method = "step",
                 variable = "window",
                 d_h = 1,
                 d_k = 10) 
    }
    )

#Fourier Window Function
LS_data[["fourier_window"]] <- 
  week2 %>%
  mutate(Time = as.numeric(Time)) %>%
  split(.$ID) %>%
  lapply(function(x) {
    FourierWindow(t = x$Time, f = f_grid) %>%
      data.frame(PSD = .,
                 ID = unique(x$ID),
                 period = T_grid,
                 method = "fourier",
                 variable = "window",
                 d_h = 1,
                 d_k = 3) 
  })


LS_data <- LS_data %>%
  lapply(function(x) {
    lapply(x, function(x) {
        x %>%
        as_tibble() %>%
        inner_join(distinct(dplyr::select(week2, ID, Sex, Density)), 
                   by = "ID") %>%
        select(ID, Sex, Density, period, PSD, method, variable, d_h, d_k)
      
    }
    )
  }
  ) %>%
  lapply(bind_rows) %>%
  bind_rows()

#Compute FAPs for each freq
LS_data <- week2 %>%
  group_by(ID) %>%
  summarize(N = n(),
            var_t = var(as.numeric(Time))) %>%
  inner_join(LS_data) %>%
  group_by(ID, variable, method) %>%
  mutate(FAP_approx = ifelse(!is.na(d_h) | !is.na(d_k),
                             FAP_approx(z = PSD, 
                                        N =  N, 
                                        d_k = d_k, 
                                        fmax = fmax, 
                                        T =  T),
                             NA),
         FAP_davies = ifelse(!is.na(d_h) | !is.na(d_k),
           FAP_davies(z = PSD, 
                      N = N, 
                      d_k = d_k, 
                      d_h = d_h,
                      var_t = var_t, 
                      fmax = fmax),
           NA)
         )

write_csv(LS_data, here::here("results","LS_data.csv"))

#Find Periodogram Minima/Maxima
#Long calculations
#Cache Results
# periodogram_methods <- c("step", "GenLS", "fourier")
# variables <- c("speed", "nest_dist", "window")
# LS_features <- list()
# PSD_sm <- list()
# for(i in periodogram_methods) {
#   for(j in variables){
#     if((i == "GenLS" & j == "window") | (i == "fourier" & j != "window")) {
#       next
#     }
#     for(k in unique(LS_data$ID)){
#       #Calcaulate smoothed LSP
#       feat_ls <- LS_data %>%
#         filter(method == i, variable == j, ID == k) %>%
#           {features(1/.$period, .$PSD, control = list(decim.out = 22))} 
# 
#       #Formate Featues as data_frames
#       LS_features[[paste(i, j, k, sep = "_")]] <- feat_ls %>%
#         {data_frame(ctps = .$cpts,
#                d2 = .$curvature,
#                method = i,
#                variable = j,
#                ID = k)}
#       
#       #Format Smoothed LSP as data_frames
#       PSD_sm[[paste(i, j, k, sep = "_")]] <- attributes(feat_ls)$fits$fn %>%
#         {data_frame(PSD_sm = ., 
#                 method = i,
#                 variable = j, 
#                 ID = k)}
#     }
#   }
# }
# LS_features <- bind_rows(LS_features)
# LS_data <- LS_data %>%
#   inner_join(bind_rows(LS_sm))
# 
# 
#   
#   periodogram_methods <- x %>%
#     dplyr::select(method) %>%
#     distinct() %>%
#     filter(method != "fourier") %>%
#     pull(method)
#   
#   variables <- x %>%
#     dplyr::select(method) %>%
#     distinct() %>%
#     filter(variable != "window") %>%
#     pull(variable)
#   
#   ids <- x %>%
#     dplyr::select(ID) %>%
#     distinct() %>%
#     pull(ID)
#   
#   for (i in periodogram_methods){
#     for (j in variables){
#       for (k in ids){
#         win_ctps <- filter(x, variable = "window",
#                             method == i,
#                             ID == k) %>%
#           pull(ctps)
#         var_ctps <- filter(x, variable != "window",
#                             method == i,
#                             ID == k)
#         index <- logical(nrow(var_ctps))
# 
# 
# #Plot LSP maxima  and smoothed LSP
# par(ask = TRUE)
# for (i in periodogram_methods) {
#   for(j in variables) {
#     for (k in unique(LS_data$ID)){
#       if((i == "GenLS" & j == "window") | (i == "fourier" & j != "window")) {
#         next
#       } 
#       index <- c(paste(i, j, sep ="_"), k)
#       
#       PSD_sm <- attributes(LS_features[[index]])$fits$fn 
#       names(PSD_sm) 
#       plot_df <- LS_data %>%
#         filter(method == i,
#                variable == j,
#                ID == k) %>%
#         bind_cols(PSD_sm)
#       
#       
#       n_maxima <- length(LS_featurues[[c(index, "ctps")]])
#       maxima_df <- data_frame(ctps = LS_featurues[[c(index, "ctps")]],
#                               d2 = LS_featurues[[c(index, "curvature")]],
#                               PSD = numeric(n_maxima),
#                               PSD_sm = numeric(n_maxima)) %>%
#         filter(d2 < 0)
#       
#       for (z in 1:n_maxima) {
#         maxima_df$PSD[z] <- plot_df$PSD[which.min(ctsp - plot_df$PSD)]
#         maxima_df$PSD_sm[z] <- plot_df$PSD[which.min(ctsp - plot_df$PSD_sm)]
#       }
#       
#         ggplot(ploft_df, aes(period, PSD)) +
#           geom_line() +
#           geom_line(aes(period, PSD_sm), color = "blue")+
#           geom_point(aes(ctsp, PSD), color = "red", data = maxima_df) +
#           labs(title = paste(i, j, k))
#     }
#   }
# }
# par(ask = FALSE)

#Check for 2nd and 3rd harmonics of window maxes

#Plot LSP minus window maxes and harmonics
```
