## Supplementary material for "An Automated Radio-Telemetry System (ARTS) for Monitoring Small Mammals": Analysis R Code: outlier_sensitivity.R

```
library(here)
library(tidyverse)
library(data.table)
library(dtplyr)
library(lubridate)
library(rjags)
library(ggmcmc)


func_dir <- here::here("src", "ARTSFunctions.R")
source(func_dir)

#Examining Model Sensitivity to "outliers". The strategy here is to refit the 
#model for the bayes_3 and TrackingAug2 methods excluding timepoints above a
#given precentile of error

training_predictions <- read_csv(here::here("results", 
                                            "training_residuals.csv")) %>%
  mutate(Time = with_tz(Time, tzone = "America/Chicago"))

#Load Antenna Coordinates for table calcs
antenna_dir <- here::here("data", "All Radiomapping Keys", "Antenna Key.csv")
ant_coor<- read_csv(file = antenna_dir) %>%
  
  #Label antenna 1-32, convert feet to meters
  mutate(Ant = AntEdit(.),
         X = X * 0.3048,
         Y = Y * 0.3048)

#Find and save time points falling within given quantiles
quantile_probs = c(0.8, 0.85, 0.9, 0.95, 0.99, 1)
training_data = c("raw", "TrackingAug2")
quant_error <- list()
model_dir = here::here("src", "model.4.txt")
for(i in training_data) {
  for(j in quantile_probs){
    quantile_times <- 
      training_predictions %>%
      filter(training == i, 
             method == "bayes",
             ant_selection == "top3") %>%
      filter(residual <= quantile(residual, j)) 

    training_df <-
      switch(i,
             raw = here::here("results", "Mapping_2014_raw.csv"),
             TrackingAug2 = here::here("results", 
                                       "Training_TrackingAug2.csv")) %>%
        read_csv() %>%
        mutate(Time = with_tz(Time, tzone = "America/Chicago"),
               Time = round_date(Time, unit = "min"),
               Ant = AntEdit(.)) %>%
      semi_join(quantile_times, by = c("Time", "Freq"))
   
    
    #Bayesian Model
    samples <- BuildJAGS(training_df, model_dir, seed = 1234)
    
    #Diagnostic plots in ggmcmc
    if(!dir.exists(here::here("docs", "Outlier_Sensitivity"))) {
      dir.create(here::here("docs", "Outlier_Sensitivity"))
    }
    plot_dir <- here::here("docs",
                           "Outlier_Sensitivity",
                           paste(i, 
                                 j,
                                 "ggmcmc.pdf",
                                 sep = "_"))
    ggmcmc(ggs(samples),
           file = plot_dir,
           param_page = 6)
    
    #Format Parameters
    BHR_quant_coeffs <- JAGSCoefs(samples)

    #Load test data
    testing_df <-
      switch(
        i,
        raw = here::here("results", "Manual_ARTS_2014.csv"),
        TrackingAug2 = here::here("results", "Testing_TrackingAug2.csv")
        ) %>%
      read_csv() %>%
    
      #bin times note that this decrease temporal resolutsion some locations in mapping
      #data are assigned to the same time
      mutate(Time = with_tz(Time, tzone = "America/Chicago"),
             Time = round_date(Time, unit = "min"),
             Ant = AntEdit(.)) %>%
      rename(trans_x = x, trans_y = y)
    
    #Calculatre or load Bayesian pdf tables for raw mapping
    bayes_tables <- CalcTablesBayes(BHR_quant_coeffs, ant_coor)
    
    prediction <- ARTSPrediction(testing_df, bayes_tables)
    
    prediction <- mutate(prediction , 
                             training = i,
                             quantile = j) %>%
      rename(pred_x = x, pred_y = y)%>%
      tbl_df() %>%
      inner_join(testing_df) %>%
      dplyr::select(pred_x, 
                    pred_y,
                    trans_x, 
                    trans_y, 
                    entropy, 
                    Time, 
                    Freq, 
                    training, 
                    quantile) %>%
      mutate(error = sqrt((trans_x - pred_x)^2 + (trans_y - pred_y)^2)) %>%
      distinct()
    
    quant_error[[paste(i,j, sep = "_")]] <- prediction
  }
}

quant_error <- rbindlist(quant_error)
write_csv(quant_error, here::here("results","outlier_errors.csv"))
```
