## Supplementary material for "An Automated Radio-Telemetry System (ARTS) for Monitoring Small Mammals": Analysis R Code: size_sensitivity.R

```
#Splitting the manual fixes into a training data set to be added to the mapping data and  and a testting data set 
library(tidyverse)
library(here)        
library(data.table)
library(dtplyr)
library(lubridate)
library(rjags)
library(ggmcmc)

func_dir <- here::here("src", "ARTSFunctions.R")
source(func_dir)

#Script Begins
mapping.dir <- here::here("results", "Mapping_2014_raw.csv")
Mapping_2014 <- read_csv(mapping.dir)
Mapping_2014 <- mutate(Mapping_2014,
                       Time = with_tz(Time, tzone = "America/Chicago"),
                       data = "mapping")

# Load Manual Fixes vs Automated Fixes from 2014 data filtering
Man_ARTS.dir <- here::here("results", "Manual_ARTS_2014.csv")
Manual_ARTS_2014 <- read_csv(Man_ARTS.dir)
Manual_ARTS_2014 <- mutate(Manual_ARTS_2014, 
                          manual_time = with_tz(manual_time, tzone = "America/Chicago"),
                          Time = with_tz(Time, tzone = "America/Chicago"),
                          data = "manual") %>%
  filter(time_diff <= 5)

set.seed(1234)
sample_fractions <- seq(0.1, 1, 0.1)
training_ls <- vector("list", length(sample_fractions))
testing_ls <- vector("list", length(sample_fractions))

for (i in seq_along(sample_fractions)){
  
  #Split manual fixes randomly into training and testing sets
  manual_training <- distinct(Manual_ARTS_2014, manual_time, Freq) %>%
    sample_frac(sample_fractions[i])
  manual_testing <- anti_join(Manual_ARTS_2014, manual_training)
  
  manual_training <- inner_join(Manual_ARTS_2014, manual_training)
  
  #Split radiomapping randomly into training and testing sets
  mapping_training <- distinct(Mapping_2014, trans_x, trans_y) %>%
    sample_frac(sample_fractions[i])
  mapping_testing <- anti_join(Mapping_2014, mapping_training) 
  
  mapping_training <- inner_join(Mapping_2014, mapping_training)
  
  #Concatenating Mapping training data and tracking training data
  training_ls[[i]] <- mutate(manual_training,
                        trans_x = x, 
                        trans_y = y,
                        distance = sqrt((trans_x - ant_x)^2 + (trans_y - ant_y)^2)) %>%
    dplyr::select(Time, 
                  Site,
                  Ant,
                  Freq,
                  Code, 
                  Power,
                  ant_x, 
                  ant_y,
                  trans_x,
                  trans_y,
                  distance,
                  data) %>%
    list(dplyr::select(mapping_training, -direction)) %>%
    rbindlist() %>%
    mutate(size = sample_fractions[i])
  
  #Concatenating Mapping "testing" data and tracking testing data
  testing_ls[[i]] <- mutate(manual_testing,
                                trans_x = x, 
                                trans_y = y,
                                distance = sqrt((trans_x - ant_x)^2 + (trans_y - ant_y)^2)) %>%
    dplyr::select(Time, 
                  Site,
                  Ant,
                  Freq,
                  Code, 
                  Power,
                  ant_x, 
                  ant_y,
                  trans_x,
                  trans_y,
                  distance,
                  data) %>%
    list(dplyr::select(mapping_testing, -direction)) %>%
    rbindlist() %>%
    mutate(size = sample_fractions[i])
}

model_dir = here::here("src", "model.4.txt")
BHR_size_coeffs <- vector("list", length(training_ls))

for(i in seq_along(training_ls)){
  training_df <- training_ls[[i]] %>%
    mutate(Ant = AntEdit(.))
  
  samples <- BuildJAGS(training_df, model_dir, seed = 1234)
  
  
  #Diagnostic plot in ggmcmc
  if(!dir.exists(here::here("docs", "Size_Sensitivity"))) {
    dir.create(here::here("docs", "Size_Sensitivity"))
  }
  plot_dir <- here::here("docs",
                         "Size_Sensitivity",
                         paste0(unique(training_df$size),
                                "_ggmcmc.pdf"))
  ggmcmc(ggs(samples),
         file = plot_dir,
         param_page = 6)
  
  #Format and Save Parameters
  BHR_size_coeffs[[i]] <- JAGSCoefs(samples)
  names(BHR_size_coeffs)[i] <- unique(training_df$size)
}

#Fit testing fixes for each sample size
names(testing_ls) <- lapply(testing_ls, function(x){
  unique(as.character(x$size))
}
)

size_errors <- vector("list", length(testing_ls) - 1)

#Load Antenna Coordinates
antenna_dir <- here::here("data", "All Radiomapping Keys", "Antenna Key.csv")
ant_coor<- read_csv(file = antenna_dir) %>%
  
  #Label antenna 1-32, convert feet to meters
  mutate(Ant = AntEdit(.),
         X = X * 0.3048,
         Y = Y * 0.3048)

#Loop over All training sets
for(i in 1:(length(testing_ls)-1)){
  
  #Select data source
  testing_df <- testing_ls[[i]] %>%
    mutate(Time = round_date(Time, unit = "min"),
           Ant = AntEdit(.),
           size = as.character(size)) %>%
    rename(data_source = data)
  
  #Calculate Bayesian pdf tables 
  model <- BHR_size_coeffs[[names(testing_ls)[i]]]
  bayes_tables <- CalcTablesBayes(model, ant_coor)
  
  prediction <- ARTSPrediction(testing_df, bayes_tables)
  
  prediction <- mutate(prediction, 
                           size = names(testing_ls)[i]) %>%
    rename(pred_x = x, pred_y = y)%>%
    tbl_df() %>%
    inner_join(testing_df) %>%
    dplyr::select(pred_x, 
                  pred_y,
                  trans_x, 
                  trans_y, 
                  entropy, 
                  Time, 
                  Freq,
                  data_source,
                  size) %>%
    mutate(error = sqrt((trans_x - pred_x)^2 + (trans_y - pred_y)^2)) %>%
    distinct()
  
  size_errors[[i]] <- prediction
}

size_errors <- rbindlist(size_errors)
write_csv(size_errors, here::here("results", "size_errors.csv"))
```
