## Supplementary material for "An Automated Radio-Telemetry System (ARTS) for Monitoring Small Mammals": Analysis R Code: split_training_testing.R

```
#Splitting the manual fixes into a training data set to be added to the mapping data and  and a testting data set 
library(here)
library(tidyverse)
library(data.table)
library(dtplyr)
library(lubridate)

mapping.dir <- here::here("results", "Mapping_2014_raw.csv")
Mapping_2014 <- read_csv(mapping.dir)
Mapping_2014 <- mutate(Mapping_2014,
                       Time = with_tz(Time, tzone = "America/Chicago"))

# Load Manual Fixes vs Automated Fixes from 2014 data filtering
Man_ARTS.dir <- here::here("results", "Manual_ARTS_2014.csv")
Manual_ARTS_2014 <- read_csv(Man_ARTS.dir)
Manual_ARTS_2014 <- mutate(Manual_ARTS_2014, 
                          manual_time = with_tz(manual_time, tzone = "America/Chicago"),
                          Time = with_tz(Time, 
                                              tzone = "America/Chicago")) %>%
                   filter(time_diff <= 5)

#Split manual fixes into training and testing sets
training_times <- distinct(Manual_ARTS_2014, manual_time, Freq) %>%
  slice(seq(1, nrow(.), by = 2))
Manual_ARTS_2014_TrackingAug <- anti_join(Manual_ARTS_2014, training_times)

training_data <- inner_join(Manual_ARTS_2014, training_times)
write_csv(Manual_ARTS_2014_TrackingAug, 
          here::here("results", "Manual_ARTS_2014_TrackingAug.csv"))

#Adding training data from 2014 tracking to mapping data
Mapping_2014_TrackingAug<- mutate(training_data,
                                  trans_x = x,
                                  trans_y = y,
                                  distance = sqrt((trans_x - ant_x)^2 + (trans_y - ant_y)^2)) %>%
                          dplyr::select(Time, 
                                         Site,
                                         Ant,
                                         Freq,
                                         Code, 
                                         Power,
                                         ant_x, 
                                         ant_y,
                                         trans_x,
                                         trans_y,
                                         distance) %>%
                          list(dplyr::select(Mapping_2014, -direction)) %>%
                          rbindlist()
write_csv(Mapping_2014_TrackingAug, 
          here::here("results", "Mapping_2014_TrackingAug.csv"))

#Split manual fixes randomly into training and testing sets
set.seed(1234)
training_times2 <- distinct(Manual_ARTS_2014, manual_time, Freq) %>%
  sample_frac(0.5)
Tracking_testing2<- anti_join(Manual_ARTS_2014, training_times2)

Tracking_training2 <- inner_join(Manual_ARTS_2014, training_times2)

#Split radiomapping randomly into training and testing sets
mapping_training_xy <- distinct(Mapping_2014, trans_x, trans_y) %>%
  sample_frac(0.5)
Mapping_testing <- anti_join(Mapping_2014, mapping_training_xy) %>%
  mutate()

Mapping_training <- inner_join(Mapping_2014, mapping_training_xy)

#Concatenating Mapping training data and tracking training data
Training_TrackingAug2<- mutate(Tracking_training2,
                               trans_x = x, 
                               trans_y = y,
                               distance = sqrt((trans_x - ant_x)^2 + (trans_y - ant_y)^2)) %>%
  dplyr::select(Time, 
               Site,
               Ant,
               Freq,
               Code, 
               Power,
               ant_x, 
               ant_y,
               trans_x,
               trans_y,
               distance) %>%
  list(dplyr::select(Mapping_training, -direction)) %>%
  rbindlist()
write_csv(Training_TrackingAug2, 
          here::here("results", "Training_TrackingAug2.csv"))

#Concatenating Mapping "testing" data and tracking testing data
Testing_TrackingAug2<- mutate(Tracking_testing2,
                              distance = sqrt((x - ant_x)^2 + (y - ant_y)^2)) %>%
  dplyr::select(Time, 
         Site,
         Ant,
         Freq,
         Code, 
         Power,
         ant_x, 
         ant_y,
         x,
         y,
         distance) %>%
  list(dplyr::select(Mapping_testing, -direction)) %>%
  rbindlist()
write_csv(Testing_TrackingAug2, 
          here::here("results", "Testing_TrackingAug2.csv"))
```
